# Reproducible Discovery of Cell-Binding Peptides “Lost” in Bulk Amplification via Emulsion Amplification in Phage Display Panning

**DOI:** 10.1101/2021.10.31.466683

**Authors:** Wadim L. Matochko, Frédérique Deiss, Yang Yang, Ratmir Derda

## Abstract

Many pharmaceutically-relevant cell surface receptors are functional only in the context of intact cells. Phage display, while being a powerful method for the discovery of ligands for purified proteins often fails to identify a diverse set of ligands to receptors on a cell membrane mosaic. To understand this deficiency, we examined growth bias in naïve phage display libraries and observed that it fundamentally changes selection outcomes: The presence of growth-biased (parasite) phage clones in a phage library is detrimental to selection and cell-based panning of such biased libraries is poised to yield ligands from within a small parasite population. Importantly, amplification of phage libraries in water-oil emulsions suppressed the amplification of parasites and steered the selection of biased phage libraries away from parasite population. Attenuation of the growth bias through the use of emulsion amplification reproducibly discovers the ligands for cell-surface receptors that cannot be identified in screen that use conventional ‘bulk’ amplification.

Find Ligands in Droplets
Canonical phage display selection of ligands for breast cancer cells, which uses bulk amplification (BA) of phage library, reproducibly identified peptide ligands from a ~0.0001% sub-population of the library, which harbors fast-growing phage. Replacing BA by emulsion-amplification (EmA) altered the selection landscape and yielded cell-binding ligands not accessible to conventional phage-display select

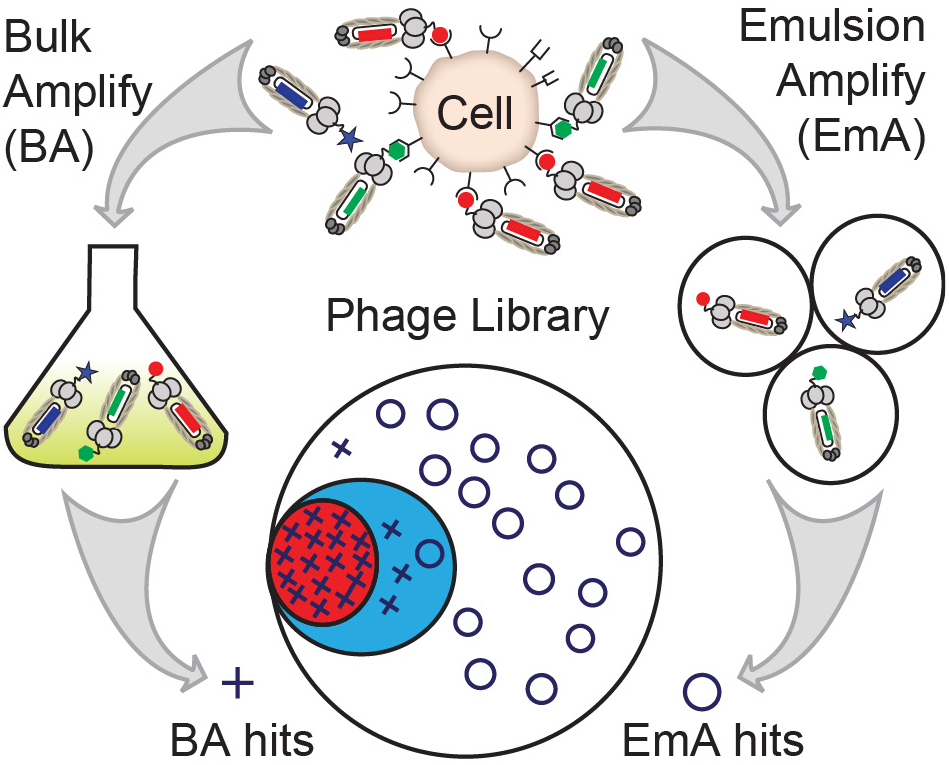

---

Genetically-encoded (GE) libraries displayed on phage,^[1]^ yeast,^[2]^ or RNA^[3]^ are powerful technologies for the discovery of ligands for virtually any molecular target, including many therapeutically-relevant targets.^[4]^ They also permit selection of ligands that bind multi-target entities such as cells and organs and the antibody repertoire (reviewed in^[5]^ ^[6]^). Functional ligands emanating from the multi-target screens can give rise to therapeutic candidates^[4, 7]^ or targeting probes^[5, 8]^ and instructive materials that control stem cell differentiation^[9]^ and self-renewal.^[10]^ All *in vitro* GE-selection strategies start from a diverse library of >10^9^ ligands, and increase the abundance of target-binding ‘hits’ in the sequence pool via rounds of panning—retention of binding and removal of non-binding ligands—and re-amplification of recovered ligands. These steps exert two orthogonal selection pressures: (i) panning selects for ligands that bind to the target; (ii) re-amplification selects the library clones that exhibit higher rates of amplification than the population average. Characterization of the amplification bias in the selection of ligands from GE-libraries is critical for improving the reproducibility and efficiency of discovery of ligands from these libraries.

Amplification biases have been characterized in oligonucleotide libraries,^[11]^ mRNA-displayed libraries,^[12]^ and in phage-displayed libraries of peptides.^[13]^ In phage-display specifically, bias originates from mutations in the ribosome-binding sequence^[14]^ or in the (+)-origin;^[15]^ clones that harbor these mutations (a.k.a., “parasites”) have higher propagation rates and these parasites often recur in selection procedures.^[13a]^ They can be identified prospectively by sequencing naïve and amplified libraries.^[13a]^ Unwanted enrichment of parasites in phage libraries,^[16]^ genomic DNA libraries,^[17]^ and in SELEX (ref^[18]^ and references within) can be minimized by employing emulsion amplification (EmA) of these libraries inside monodisperse aqueous droplets. Although the benefits of EmA in nucleotide libraries are documented, the effect of EmA on the outcome of selection of other GE-libraries is not know. In this manuscript, we characterize the role of amplification bias in the selection procedure against intact cells. As the cell contains several thousand distinct cell surface receptors, selection pressure in such screens is ‘weak’ and these selections are steered towards parasite sub-population. Reproducibility of such discovery is coupled to the amplification bias: clones that have amplification bias (“parasites”) are discovered reproducibly. Decreasing the amplification bias by employing EmA steers the selection landscape away from parasite population. Reproducibility of such discovery is driven by the binding preferences of the clones and it is not related to their amplification preferences.

As a model multi-receptor target, we employed the breast cancer cell line MDA-MB-231 (“MB-231”, Figure S1). Phage-displayed library of 10^9^ 7-mer peptides in M13KE vector (PhD-7) is a convenient library for such selection because the molecular mechanism for biases in this library are well characterized and this library has been used in over 1000 publications to date (source: BioPanning Database^[19]^). We categorized both libraries into: the ‘visible population’ (V) defined as 10^6^-10^7^ sequences identified by Illumina sequencing and the ‘invisible population’ (I) corresponding to the set of all possible library members excluding the visible population (Figure 1D). Within the V-population, we mapped the ‘parasite population’ (P) by re-amplification of the library in the absence of selection.^[13a]^ In the PhD-7 library (Figure S2), P-population consisted of ~10^3^ sequences that increased significantly (p<0.05) in such amplification; and the magnitude of bias defined as population average [Amp]/[Naive] was a factor of 10 (Figure 1B-D and Figure S2).

**Figure 1.**
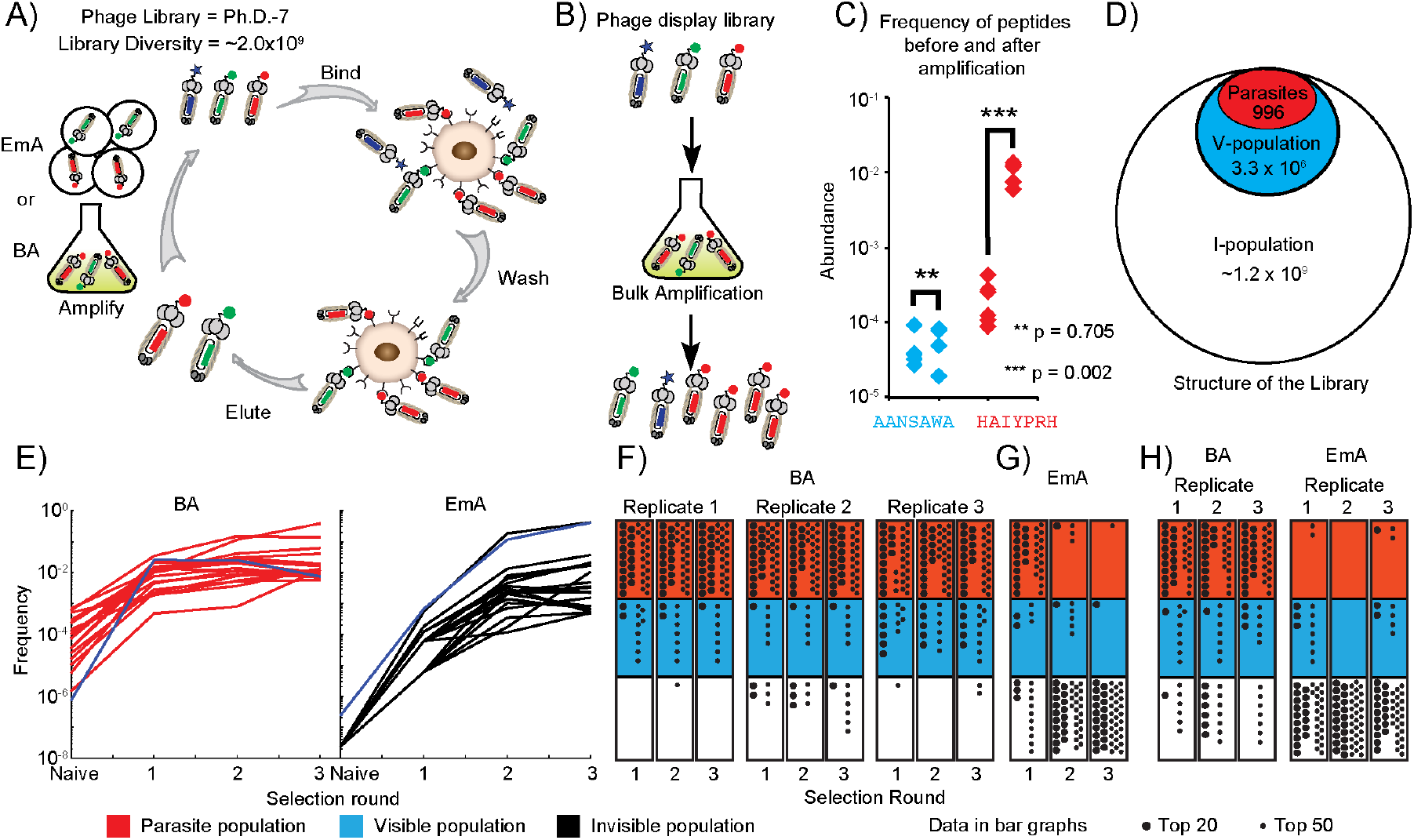
Mapping of enriched sequences to library diversity. (A) Schematic representation of two independent selections performed with a Ph.D.-7 library against breast cancer cell line, MDA-MB-231. In one selection, the eluted library was amplified in a bulk solution (BA), in the other; the eluted library was amplified in emulsion (EmA). (B) Phage display libraries contain clones with high growth rate (parasites). To identify parasites, a phage library was amplified, deep-sequenced and processed to select sequences that increased in abundance significantly (p < 0.05) when compared to the naïve population. We used volcano analysis to analyze all members of the naïve and amplified libraries. (C) The abundance of the representative non-parasite sequence (AANSAWA) and the parasite (HAIYPRH) before and after amplification. (D) We categorized the phage library as either visible (V-population), or invisible (I-population) in deep-sequence, or parasites (P-population). (E) Tracing the origin of the top 20 hits after 3 rounds of BA and EmA selections. Each trace describes a unique peptide sequence. The color of the sequences describes its origin (red **– ‘**parasite’; blue **– ‘**visible’; black **– ‘**invisible’). (F-G) The origin of the top 50 sequences enriched in EmA and three separate replicates of BA selection. Each ‘dot’ represents a unique sequence (large dot: top 20; small dot: top 50). (H) Analysis of one round of selection with a different lot of a Ph.D.-7 library (lot #0081212) and classification of the top 20/50 sequences. The EmA-screen enriches for sequences predominantly from the I-population, whereas the BA-screen enriches sequences from the P-population.

Three independent replicates from selection of strongly biased phage library against MB-231 reproducibly yielded binding peptide clones from the P-population (Figure 1F). In rounds 1 through 3, in a representative screen, 39 out of the top 50 unique sequences originated from P-population and 11 from V-population (Figure 1F and S3 show). The probability for 39/50 of unique sequences to originate from a population that has 10^3^/10^9^ = 0.0001% of the unique sequences is negligible (p~10^−39*6^). To show that the steering of the selection to the P-population was caused by a growth bias, we reduced the growth bias by using EmA in place of the conventional “bulk amplification” (BA) used in the selection. All other parameters in the selection—type of cell, type and amount of library, selection procedure (i.e., incubation time, number and stringency of washes, etc.)—remained unchanged. After three rounds of EmA-selections, contribution of clones from the P-population diminished to 4/50 and 1/50 in rounds 2 and 3 of panning; 48/50 sequences in round 3 originated from the I-population not accessible to BA-panning.

Steering of the selection towards or away from the parasite population was reproducible in other libraries. To demonstrate it, we used an independently manufactured lot of 7-mer peptide library. It is now well-established that the major mechanism for growth advantage in phage libraries build in M13KE plasmid result from mutations in a distal regulatory region of the phage genome (e.g., RBS of *pII* gene).^[14a]^ Growth bias is not related to the identity of the displayed sequences. At the time of the cloning of the library, a rare mutated vector is randomly matched with a specific peptide; the phage bearing such peptide is a “parasite”. As a result, identity of peptides in parasite populations in Ph.D-7 lot #2 termed {P_2_} was completely non-overlapping with {P_1_} (Figure 1D, S2B-C). Nevertheless, selection from lot#2 was steered to this {P_2_} population. In three independent instances of BA-screen from this library, 30-35 of the top 50 sequences originated from {P_2_} population (Figure 2H). Employing the EmA steered the selection away from {P_2_} yielding only 2/50, 0/50, and 3/50 sequences from the P-population in three independent screens. Steering towards {P_2_} population was reproducible for other targets: in screens against HEK-293 cell the top 50 enriched sequences were different from those identified in MDA-231 screen; however, 30-35 of these top 50 sequences originated from {P_2_} population (Table S3). These observations suggest the generality of the phenomena: libraries cloned in M13KE background contained a defined P-population and these libraries are biased to selecting ligands from the P-population in any multi-receptor screen. Suppressing this population by EmA, alleviates the bias in the screen.

**Figure 2.**
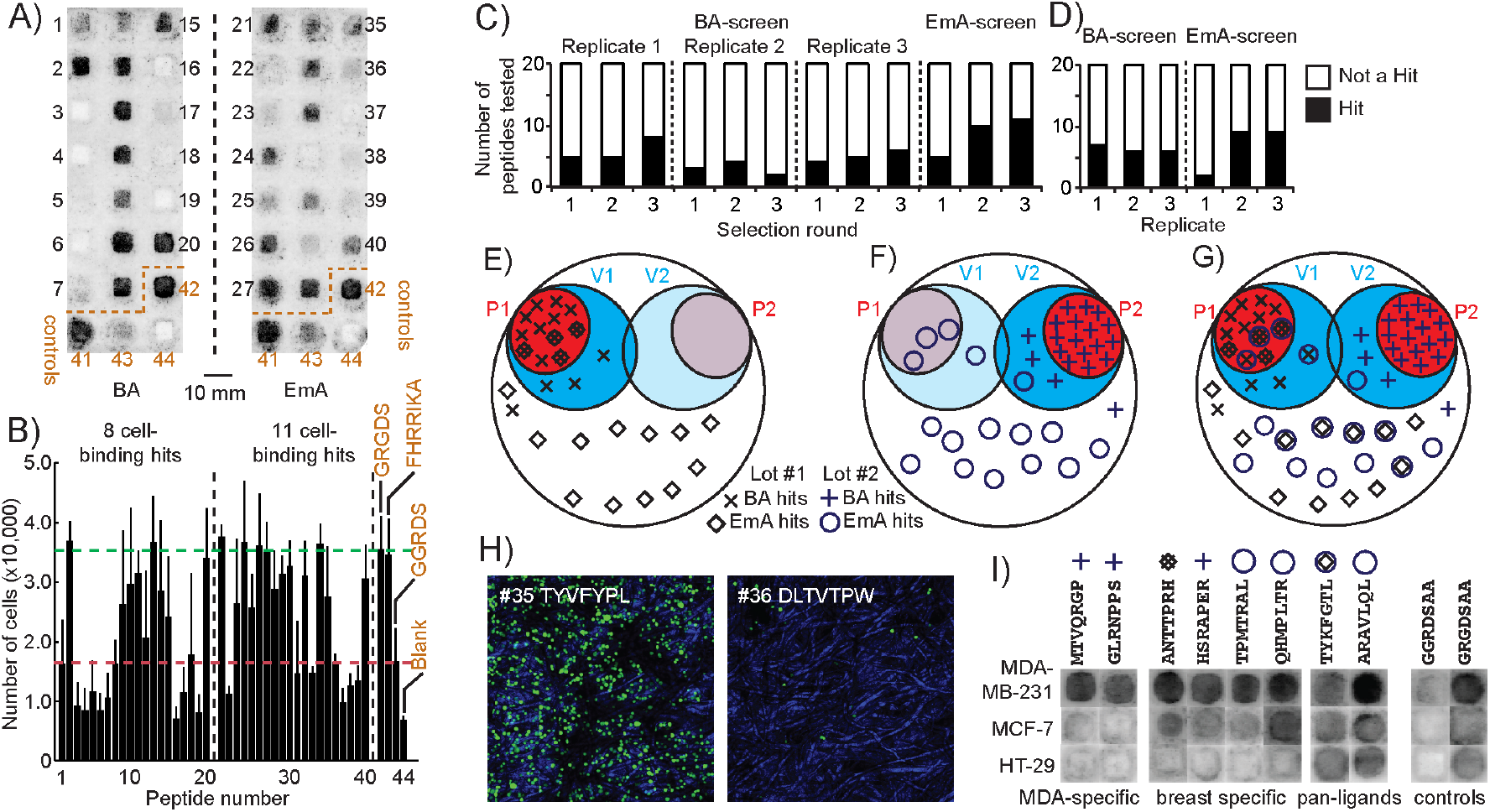
Validation of cell-binding hits. (A) We validated the top 20 sequences from each round and replicate of selection by measuring adhesion of MDA-MB-231-GFP cells to peptide arrays on paper. Representative fluorescent gel scanner image of cells adhering to the array displaying the top 20 peptide sequences after Round 3 of BA and EmA selection. (see Figure S2b for list of sequences). Integrin-(GRGDS) and Heparin-binding (FHRRIKA) peptides were the positive controls; unmodified paper and scrambled GGRDS sequence were the negative controls. (B) The greyscale intensity for each peptide was converted to number of cells using a standard calibrartion curve (Figure S7). Hit peptides supported adhesion of significantly more (p < 0.05) cells than the negative control (GGRDS, red line). (C-D) Summary of the hit and non-hit sequences among the top 20 peptides from each selection replicate and round performed with two Ph.D.-7 libraries, lot #0061101 (C) and #0081212 (D). (E-F) Location of all hits identified from BA and EmA selections in the P-population (red), V-population (blue), or I-population (white) of the Ph.D.-7 libraries, lot #0061101 (E) and #0081212 (F). The overlay of P- and V- populations from two lots indicate that the <u>validated hits</u> from the BA-screen are restricted to their respective P-populations, whereas many <u>validated hits</u> in the EmA-screen emanate from the I-population. (G) Describes an overlay of top 20 hits from sections (E) and (F). We observe significant reproducibility in discovery of the top 20 peptides when EmA is used: same sequences emanate from two different lots are marked as hits if and only if EmA is used. BA-hits are perfectly divergent and restricted to their respective P-populations. (H) Representativec confocal flourescent images of two peptides (TYVFYPL and DLTVTPW) that were confirmed as hit and not a in cell-adhesion assay (for images of all peptide-zones see Figure S5-S6). (I) The MDA-MB-231 binding peptides were re-tested with MCF-7 and HT-29 cancer cell lines. Peptides were either MDA-MB-231 specific, breast cancer specific or supporting adhesion of all three cell lines (for the latter, only two out of 36 examples are shown; for a complete set see Figure S14, S15, and S16).

To estimate the affinity of the cell-binding hits enriched in panning, we synthesized the peptide sequences on Teflon-patterned paper arrays and performed short-term adhesion of MB-231-GFP cells using a previously validated assay.^[20]^ Confocal fluorescent microscopy (Figures 2H, S5-S6) confirmed the presence of cells; fluorescent scanner and calibration curve (Figure S8) estimated the number of cells adhering to each peptide (Figure 2B, S7, S9-S13). Comparing the number cells adhered to the peptides of unknown affinity to peptides with known affinity (RGD, 10 μM)^[21]^ allowed preliminary estimation of binding strength (Figure S7D).

Testing top 20 peptides from every round and every replicate of the BA- and EmA-screens—360 peptides total, 209 unique—yielded 56 unique cell-binding hits that supported adhesion of significantly more cells than the negative control (GGRDS peptide, *p <* 0.05, Figure 2C-D). The fraction of validated cell-binding peptides was ~30% in BA-screens and up to 60% in EmA-screens (Figure 2C-D). The strength of peptide-cells interaction extrapolated from cell numbers EmA was significantly higher for a population of peptides discovered in EmA, when compared to BA selection.

To assess whether the identified peptides target different receptors on MB-231 cell surface, we estimated whether MB-231-binding peptides support adhesion of closely related MCF-7 breast and distally related HT29 colon carcinoma (Figure 2B, S14-S17). Of 36 tested peptides, two bound only to MB-231 cells and four bound to MB-231 and MCF-7 but not HT29. These observations suggested that population of peptides identified in the screen indeed target distinct receptors but this hypothesis can be further strengthened in the future using techniques such as affinity pull-down and proteomic analysis.

An average cell contains several thousand molecularly distinct receptors. Our results highlight that the GE-screens in such multi-receptor milieu are sensitive to intrinsic secondary selection pressures in the GE-library. Figure 2E-F summarizes the steering of the selection of cell-binding ligands in 7-mer peptide landscape. The confirmed cell-binding ligands selected from BA-screen from lot #1 and #2 of the Ph.D.-7 library are steered to non-overlapping P_1_ and P_2_ populations respectively (Figure 2E-G). The steering towards distinct populations that comprise less than 0.0001% of available diversity makes it impossible to discover the same sequence from two different preparations of the library. The irreproducibility makes it impossible to compare the outcomes of selections: if similar libraries are produced by two different research groups, selections from such two libraries is likely to diverge and yield two different set of binding ligands.

The observed steering and biases and linked to a well-characterized bias in M13KE originating from two factors: (i) proximity of cloned LacZ*a* gene to origin or replication and regulatory regions depresses replication and makes it possible for phage to acquire rare beneficial mutations that increase the growth rate;^[14b]^ (ii) amplification via continuous infection and secretion of mature phage converts even minor growth advantages (e.g., 20% increase) to major difference in composition (100-fold increase in one amplification).^[22]^

## Conclusion

We characterize discovery trajectories in phage-displayed libraries built on M13KE genome with well-characterized bias. Simple replacement of BA with EmA yields three major outcomes: (i) EmA-selection steers the selections away from the pre-defined parasite populations; (ii) Steering away from the “parasite” population discovers ligands from the regions of the library not accessible to parasite-biased BA-screens; (ii) most remarkably, EmA-selection from independently produced libraries L_1_ and L_2_ with two distinct P_1_ and P_2_ sets reproducibly converges on the same sequences. In contract BA-selections of same libraries L_1_ and L_2_ diverge to P_1_ and P_2_ sets. Convergence of discoveries from relatively small peptide libraries of <10^9^ is anticipated and we identify the circumstances in which convergence is maximized. Maximizing the reproducibility of discovery between different instances of production or selection of the same library will be critical for effecting communication of discovery observations between different researchers. Not every library is “stirred” by EmA: in large 10^11^-scale antibody libraries, in which bias towards “parasite population” was not detectable, EmA had no detectable effect either. Still, we anticipate that EmA will serve as useful tool for other genetically-encoded libraries in which the amplification biases are inherently present and cannot be readily removed by re-engineering of the system: for example EmA has been shown to steer SELEX away from parasites populations.^[18a]^ EmA or other technologies that remove growth bias will be critical in ensuring the reproducibility of discovery from GE-libraries. We also anticipate that the impact of EmA will be the most pronounced in screens against multi-site targets: cells, organs, mixtures of antibodies isolated from serum, mixtures of proteins, or even purified with a plurality of independent binding sites.

## Supporting information

Supplementary Information (Methods, Images)

Supporting Data (Sequencing, Scripts)

## Acknowledgements

We thank Prof. Jamie K. Scott for critical review of this manuscript. We thank Farah Hassan for help in the synthesis of Teflon-patterned peptide arrays, Dax Torti at the Donnelly Sequencing Center and Sophie Dang at the Molecular Biology Service Unit for help in Illumina and Ion Torrent sequencing, respectively. The research was supported by the Alberta Glycomics Centre, NSERC Discovery Grant (RGPIN-2016-402511 to R. D.), NSERC Accelerator Supplement (to R.D.), SENTINEL Bioactive Paper Network and Grand Challenge Canada (Rising Star award to F.D.). Canada Foundation for Innovation (CFI) provided infrastructure support.

## Notes

### Competing Interest Statement

The authors have declared no competing interest.

## References

[1] J. K. Scott, G. P. Smith, Science 1990, 249, 386–390.

[2] E. T. Boder, K. D. Wittrup, Nat. Biotechnol. 1997, 15, 553–557.

[3] R. W. Roberts, J. W. Szostak, Proc. Natl. Acad. Sci. USA 1997, 94, 12297–12302.

[4] A. L. Nelson, E. Dhimolea, J. M. Reichert, Nat. Rev. Drug Discov. 2010, 9, 767–774.

[5] B. P. Gray, K. C. Brown, Chem. Rev. 2014, 114, 1020–1081.

[6] J. T. Ballew, J. A. Murray, P. Collin, M. Maki, M. F. Kagnoff, K. Kaukinen, P. S. Daugherty, Proc. Natl. Acad. Sci. USA 2013, 110, 19330–19335.

[7] A. D. Keefe, S. Pai, A. Ellington, Nat. Rev. Drug Discov. 2010, 9, 537–550.

[8] K. F. Barnhart, D. R. Christianson, P. W. Hanley, W. H. P. Driessen, B. J. Bernacky, W. B. Baze, S. J. Wen, M. Tian, J. F. Ma, M. G. Kolonin, P. K. Saha, K. A. Do, J. F. Hulvat, J. G. Gelovani, L. Chan, W. Arap, R. Pasqualini, Science Translational Medicine 2011, 3.

[9] J. Xie, H. K. Zhang, K. Yea, R. A. Lerner, Proc. Natl. Acad. Sci. USA 2013, 110, 8099–8104.

[10] R. Derda, S. Musah, B. P. Orner, J. R. Klim, L. Y. Li, L. L. Kiessling, J. Am. Chem. Soc. 2010, 132, 1289–1295.

[11] aR. R. Breaker, G. F. Joyce, Proc. Natl. Acad. Sci. USA 1994, 91, 6093–6097; bB. Zimmermann, T. Gesell, D. Chen, C. Lorenz, R. Schroeder, Plos One 2010, 5.

[12] G. Kamalinia, B. J. Grindel, T. T. Takahashi, S. W. Millward, R. W. Roberts, Chem. Soc. Rev. 2021, 50, 9055–9103.

[13] aW. L. Matochko, S. C. Li, S. K. Y. Tang, R. Derda, Nucleic Acids Res. 2014, 42, 1784–1798; bD. J. Rodi, A. S. Soares, L. Makowski, J. Mol. Biol. 2002, 322, 1039–1052.

[14] aK. T. H. Nguyen, M. A. Adamkiewicz, L. E. Hebert, E. M. Zygiel, H. R. Boyle, C. M. Martone, C. B. Melendez-Rios, K. A. Noren, C. J. Noren, M. F. Hall, Anal. Biochem. 2014, 462, 35–43; bE. M. Zygiel, K. A. Noren, M. A. Adamkiewicz, R. J. Aprile, H. K. Bowditch, C. L. Carroll, M. A. S. Cerezo, A. M. Dagher, C. R. Hebert, L. E. Hebert, G. M. Mahame, S. C. Milne, K. M. Silvestri, S. E. Sutherland, A. M. Sylvia, C. N. Taveira, D. J. VanValkenburgh, C. J. Noren, M. F. Hall, 2017, -12.

[15] W. D. Thomas, M. Golomb, G. P. Smith, Anal. Biochem. 2010, 407, 237–240.

[16] R. Derda, S. K. Y. Tang, G. M. Whitesides, Angew. Chem. Int. Ed. 2010, 49, 5301–5304.

[17] aR. Williams, S. G. Peisajovich, O. J. Miller, S. Magdassi, D. S. Tawfik, A. D. Griffiths, Nat. Methods 2006, 3, 545–550; bD. Dressman, H. Yan, G. Traverso, K. W. Kinzler, B. Vogelstein, Proc. Natl. Acad. Sci. USA 2003, 100, 8817–8822.

[18] aS. Matsumura, A. Kun, M. Ryckelynck, F. Coldren, A. Szilagyi, F. Jossinet, C. Rick, P. Nghe, E. Szathmary, A. D. Griffiths, Science 2016, 354, 1293–1296; bM. Takahashi, X. W. Wu, M. Ho, P. Chomchan, J. J. Rossi, J. C. Burnett, J. H. Zhou, Scientific Reports 2016, 6.

[19] B. F. He, G. S. Chai, Y. C. Duan, Z. Q. Yan, L. Y. Qiu, H. X. Zhang, Z. C. Liu, Q. He, K. Han, B. B. Ru, F. B. Guo, H. Ding, H. Lin, X. L. Wang, N. N. Rao, P. Zhou, J. Huang, Nucleic Acids Res. 2016, 44, D1127–D1132.

[20] F. Deiss, W. L. Matochko, N. Govindasamy, E. Y. Lin, R. Derda, Angew. Chem. Int. Ed. 2014, 53, 6374–6377.

[21] J. R. Klim, A. J. Fowler, A. H. Courtney, P. J. Wrighton, R. T. C. Sheridan, M. L. Wong, L. L. Kiessling, Acs Chemical Biology 2012, 7, 518–525.

[22] R. Derda, S. K. Tang, S. C. Li, S. Ng, W. Matochko, M. R. Jafari, Molecules 2011, 16, 1776–1803.

