## Supplementary Information (Methods, Images) for "Reproducible Discovery of Cell-Binding Peptides “Lost” in Bulk Amplification via Emulsion Amplification in Phage Display Panning"

### Table of Contents

Methods....................................................................................................S2

Figure S1. Phage titers in selection with Ph.D.-7 library lot #1 ..................................S7

Figure S2. Characterization of parasites in Ph.D.-7 libraries ......................................S8

Figure S3. Tracing top 20 sequences through selection ..............................................S9

Figure S4. Phage titers in selection with Ph.D.-7 library lot #2 ..................................S10

Table S1. Top 20 sequences from each selection .....................................................S11

Table S2. Copy number for top 20 sequences from selection ...................................S12

Table S3. Top 50 sequences from selection against HEK cells..................................S17

Figure S5. Confocal images of cells to peptides from BA-screen .............................S18

Figure S6. Confocal images of cells to peptides from EmA-screen ..........................S19

Figure S7. Work flow of cell adhesion analysis to first peptide array........................S20

Figure S8. Calibration curves ....................................................................................S22

Figure S9. Adhesion of MDA-MB-231 peptide array (2nd dataset)............................S23

Figure S10. Adhesion of MDA-MB-231 peptide array (3rd dataset)..........................S24

Figure S11. Adhesion of MDA-MB-231 peptide array (4th dataset)..........................S25

Figure S12. Adhesion of MDA-MB-231 peptide array (5th dataset)..........................S26

Figure S13. Variability in adhesion of cells to peptide array ....................................S27

Figure S14. Adhesion of MDA-MB-231, MCF-7, and HT-29 (1st data set) .............S28

Figure S15. Adhesion of MDA-MB-231, MCF-7, and HT-29 (2nd data set) ............S29

Figure S16. Comparison of MDA-MB-231, MCF-7, and HT-29..............................S30

Figure S18. Scheme for the insert used in cell-adhesion assay .................................S31

Description of the files available as part of Supporting_Files.RAR..........................S32

Reference ...................................................................................................................S35

### Methods

#### Preparation of cDNA from phage libraries for Next-Generation Sequencing.

Phage cDNA was isolated from phage libraries using QIAprep Spin M13 kit. Isolated phage cDNA was subjected to PCR amplification with primers flanking the variable region. To avoid a second round of PCR amplification, the primers contained Ion Torrent or Illumina adapters at the 5’ ends. Ion Torrent (Life Technologies) was used to sequence phage libraries obtained from selections. The cDNA was amplified using the following primers:

Forward primer: 5′-CTCTCTATGGGCAGTCGGTGATCCTTTCTATTCTCACTCT-3′

Reverse primer: 5′-CCATCTCATCCCTCGCTGTCTCCGACTCAG**XXXXXXXX**CCGAACCTCCACC

Illumina was used to sequence naïve phage libraries (lot #1 and #2). The ssDNA was amplified using the following primers: Forward primer:

5′-CAAGCAGAAGACGGCATACGAGATCGGTCTCGGCATTCCTGCTGAACCGCTCTTCCGATCT**XXXX**CCTTTCTATTCTCACTCT-3′, and

Reverse primer:

5′-AATGATACGGCGACCACCGAGATCTACACTCTTTCCCTACACGACGCTCTTCCGATCT**XXXX**ACAGTTTCGGCCGA-3′

where XXXX denotes the barcode sequence. The temperature cycling protocol was as follows: 95 °C for 30 s, followed by 25 cycles of 95 °C for 10 s, 60.5 °C for 15 s and 72 °C for 30 s, and then a final extension at 72 °C for 5 min before holding at 4°C. The concentration of the PCR fragments that resulted from amplification of phage libraries was determined by analytical gel (2 % w/v agarose gel in TBE buffer) using a low molecular weight DNA ladder as a standard (NEB, Cat# N3233S). Multiple PCR products amplified with different barcoded primers were pooled and purified using E-gel with a SizeSelect 2% gel (Invitrogen). The fragments were extracted with RNAse free water. The concentration of purified product was determined using a Qubit Fluorimeter (Invitrogen). PCR amplification of the phage library region and data processing is further described in our previous publication.[1]

**Next generation sequencing of the library.**

Prior to Ion Torrent sequencing, the library was clonally amplified on Ion Sphere Particles (ISPs). The dsDNA fragments (3 pmol) were ligated onto ISPs and amplified by emulsion PCR according to Ion Torrent protocol. The concentration of ISPs with ligated dsDNA fragments after emulsion PCR was determined using Qubit Fluorimeter according to manufacturer's protocol. The ISPs with ligated dsDNA fragments were enriched for and loaded on an Ion 316 chip. The sequencing was performed using an Ion Torrent system (Life Technologies) with an Ion OneTouch 200 Template Kit. Ion Torrent sequencing was performed at the Molecular Biology Service Unit at the University of Alberta. The Donnelly Sequencing Center at the University of Toronto performed Illumina sequencing. Processing of deep-sequencing data and statistical analysis used for prospective identification of the parasite population was performed as described in our previous report.[1]

**Cell culture.**

MDA-MB-231, MCF-7, and HT-29 were cultured in Minimum Essential Media (MEM) (HyClone) supplemented with 10% fetal bovine serum (FBS) (HyClone), 1% Non-essential amino acids (HyClone), and 1% GlutaMAX (Gibco) at 37 °C in a 5% $CO_2$ incubator. Cells were passaged every 2-3 days using trypsin (HyClone). All culture reagents were purchased from Thermo Scientific.

**Cell panning.**

A commercially available library of M13KE phage displaying a random 7-mer peptide on the pIII protein was used in all panning experiments (Ph.D.$^{TM}$-7 kit, New England Biolabs, complexity of $1.1x10^9$ individual clones, Lot #0061101and Lot #0081212). An aliquot of the Ph.D.-7$^{TM}$ library

(1 μL from $10^{13}$ pfu/mL stock) was combined with 100 μL of CM-HBS-3% BSA (1 mM $CaCl_2$, 1 mM $MgCl_2$, 150 mM NaCl, 50 mM HEPES, 3% BSA, pH=7.0) and incubated for 30 min at 4 °C. A suspension of ~$10^7$ live MDA-MB-231-GFP cells in 100 μL of CM-HBS-3% BSA was combined with the phage solution and the mixture was rocked on a rotisserie for 1 h at 4 °C. The cell-phage suspension was centrifuged for 5 min at 2,000 rpm, and the supernatant removed. To remove unbound phage, 5-7 rounds of washing were performed. Each round of washing involved the following steps: 1) re-suspending the cell pellet in 50 mL of CM-HBS-1% BSA, 2) incubating for 5 min on ice, 3) centrifugation for 5 min at 2,000 rpm, and 4) discarding the supernatant. To elute the bound phage, the cell pellet was re-suspended in 200 μL of "Elution Buffer" (0.2 M Glycine-HCl, 0.1% BSA, pH=2.2) for no more than 10 min. 30 μL of "Neutralization Buffer" (1 M Tris-HCl, pH=9.1) was added to neutralize the solution and prevent loss of phage viability. 3-5 μL from each wash and elution solutions were used to determine the phage titer (Figure S1, S4). The eluted phage was amplified either in emulsion or in bulk solutions. The input and output titers were monitored in all selection procedures and they are summarized in Figures S1 and S4.

**Amplification of phage library in bulk solution.**

The eluted phage solution was combined with 25 mL of Lysogeny broth (LB) and 250 μL of *Escherichia coli* K12 ER2738 (Ph.D.-7 kit, New England Biolabs) in log phase growth. The phage were amplified for 4.5 h at 37 °C at ~225 rpm. The culture broth was centrifuged at 5,000 rpm for 15 min at 4 °C and the supernatant was combined with 12 mL of PEG/NaCl solution (15% (w/v) $PEG_{8,000}$, 1 M NaCl) and incubated for 2 h on ice to precipitate the phage. The phage was pelleted by centrifugation at 14,000 rpm for 15 min at 4 °C, and the pellet was re-suspended in 1 mL of CM-HBS-3% BSA buffer for further rounds of panning, Ion Torrent sequencing, and titering.

**Amplification of phage library in emulsions.**

The phage solution and 115 μL of *Escherichia coli* K12 ER2738 in log phase growth were combined with LB to a final volume of 3 mL and the mixture was emulsified using a microfluidics flow-focusing device as previously described.[1-2] The emulsion was incubated for 4.5 h at 37 °C at ~40 rpm. The emulsion was destabilized using 0.5% Krytox in HFE-7100 to combine all amplified phage clones into the aqueous layer. The aqueous-perfluoro solution was centrifuged at 14,000 rpm for 2 min. The aqueous layer was removed and combined with PEG/NaCl in a 1:1 ratio. The solution was incubated for 2 h on ice to precipitate the phage. The phage solution was pelleted at 14,000 rpm for 15 min at 4 °C, and the pellet was re-suspended in 100 μL of CM-HBS-3% BSA buffer for further rounds of panning, Ion Torrent sequencing, and titering.

**Adhesion of cells to arrays of peptides on paper.**

Arrays were synthesized as described in our previous publication.[3] The peptide functionalized on paper was soaked in MilliQ $H_2O$ for 30 min in a Nunc Omni-Tray. The paper was then washed twice with 13 mL of MEM media, followed by two washes with 13 mL of MEM media (2 x 5 min at 45 rpm). A custom made insert (for design of insert, see Figure S18) was added to hold the paper submerged and the paper and insert were washed twice with 13 mL of binding-media (0.5 % BSA-MEM media). A suspension of live MDA-MB-231-GFP cells ($0.3 \times 10^5$ cells/mL) in 25 mL of binding media was added to the array and incubated for 3 h at 37 °C in a $CO_2$ incubator. The array was subsequently washed with MEM (3 x 13 mL) and imaged using a fluorescent gel scanner (GE Healthcare, Typhoon FLA9500) and a confocal fluorescent microscope (Zeiss LSM 700). Cell-binding peptides were identified by comparing the number of cells between the test and control samples using one-sided, unequal variance Student's t-test with significance threshold of 0.05. The use of parametric statistics was justified because cell-binding data is normally distributed according to Lilliefors test (Figure S13E).

**Quantification of peptides on modified cellulose support.**

Three replicates of a peptide array were treated with 50% TFA:DCM for 5 min to remove cells after the cell-adhesion assay. The arrays were washed with DCM (3 x 10 mL), methanol (3 x 10 mL) and air-dried. The middle of each peptide zone was punched out using a 2 mm hole puncher and the 2 mm paper zone was placed in a 2 mL vial and treated with $NH_3$ gas overnight. The peptides were dissolved in 50 μL of $H_2O$ and analyzed by LC-MS (Agilent Technologies 6130 LCMS). The amount of each peptide was determined by comparing the peak areas of each peptide to the peak area of an internal standard peptide.

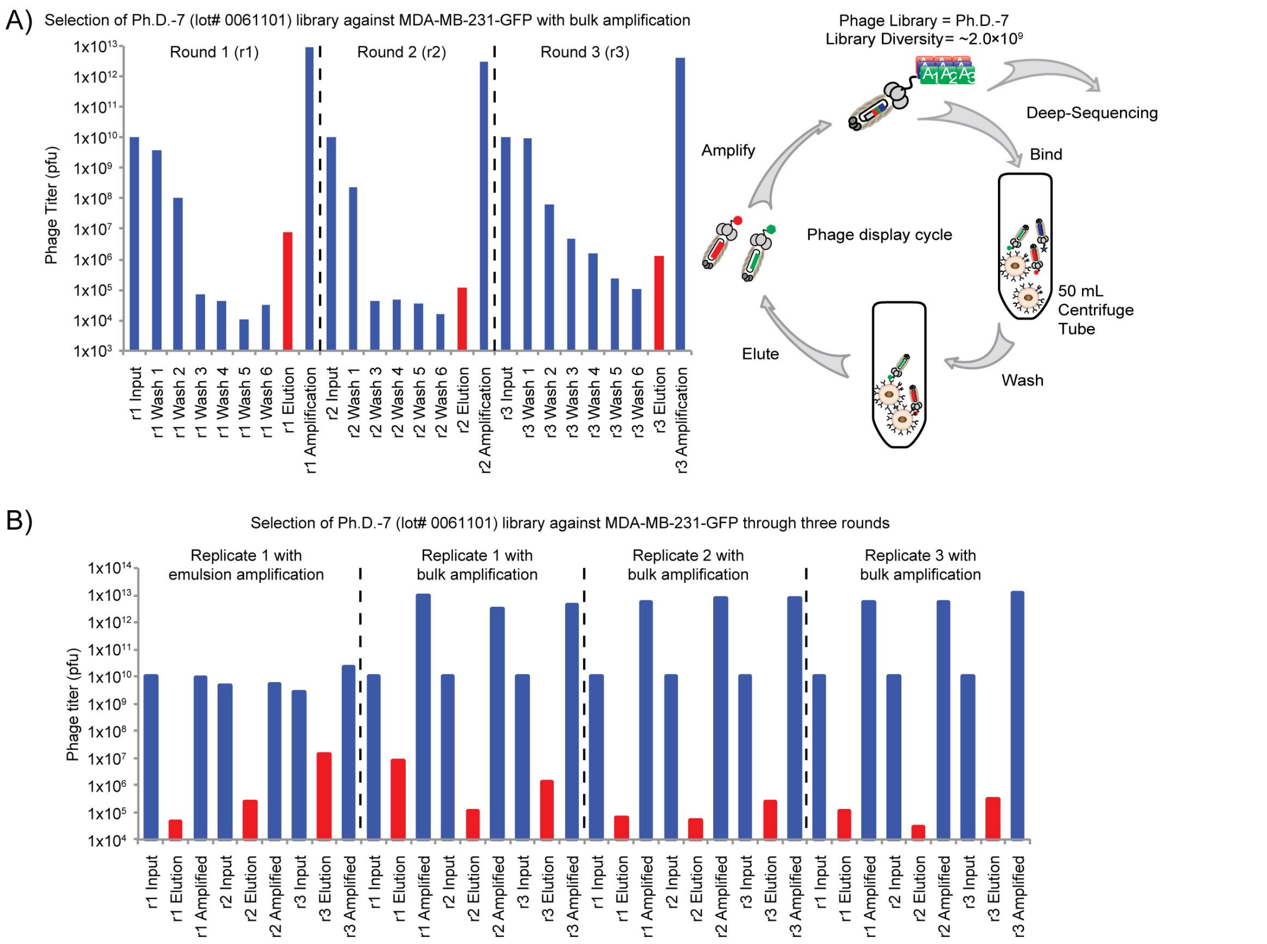


**Figure S1.** Monitoring phage titers in selection with Ph.D.-7 library lot #1. (A) Monitoring phage titers in the input selection, washes 1 through 6, elution and amplification (in BA) over three rounds of selection against MDA-MB-231-GFP cells. (B) Condensed summary of the input, output and amplified phage titers in all replicates of selection using BA and EmA. Replicate 1 with bulk amplification is the same as panel A. The output titer increased significantly through rounds of selection with EmA, but not with BA. Half of the amplified library after each round of selection was used for deep-sequencing.

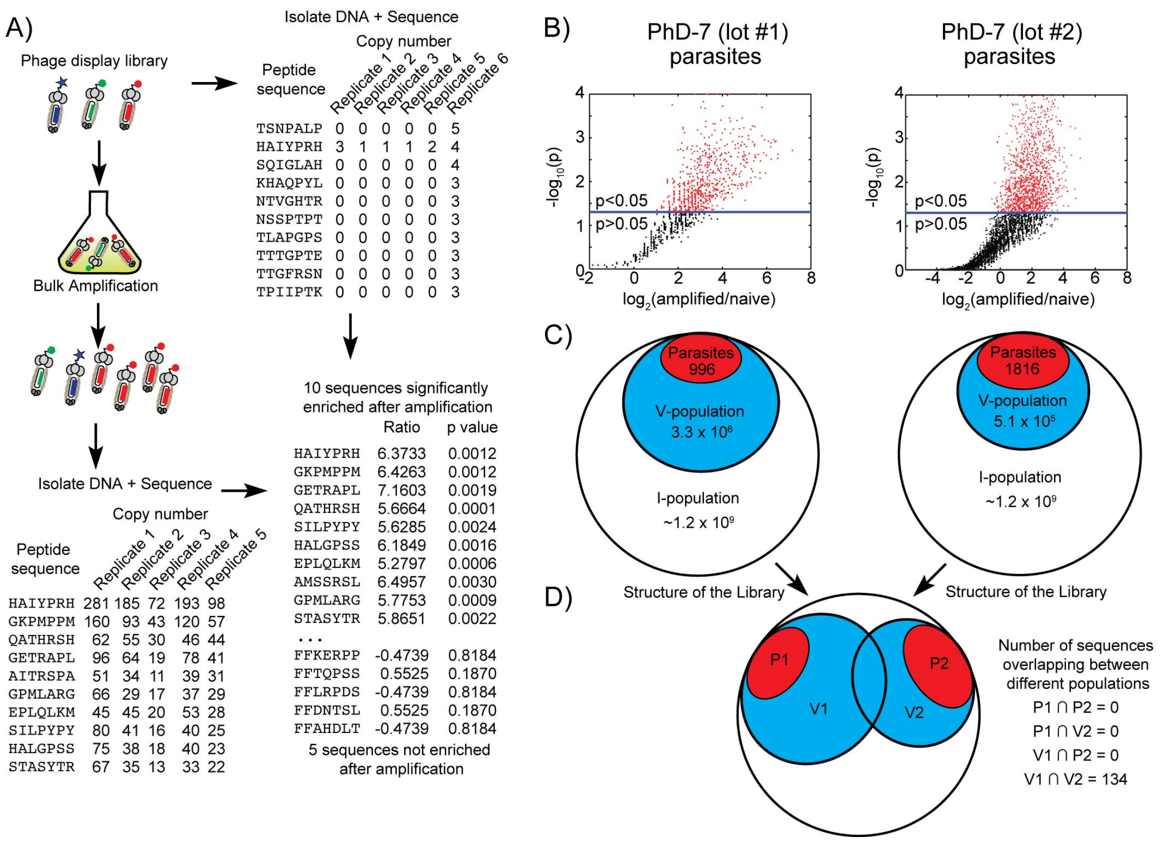


**Figure S2**. Characterization of parasites in Ph.D.-7 libraries. (A) To identify parasite sequences, the library was amplified; the cDNA from the naïve and amplified libraries was extracted and sequenced by Illumina. A representative list of 10 (out of ~3,000,000 from the naïve library) sequences sorted by copy number. For each sequence, we determined the normalized frequency, the average ratio before and after amplification and the p-value through a t-test. Sequences for which the frequency is increased significantly ($p < 0.05$) were classified as 'parasites'. (B) A volcano plot describing all parasites in the two lot of the PhD-7 libraries. (C) The structure of the two Ph.D.-7 libraries used in this report. The "invisible population" (I-population) is composed of all possible sequences of a 7-mer peptide library. The theoretical diversity is ~$1.2 \times 10^9$ sequences. Within the I-population, the "visible population" (V-population), contains all sequences that was identified by Illumina sequencing. The V-population contains a P-population with all parasite sequences identified in volcano plot (C). (D) Overlap of P1, P2, V1, and V2 populations. Interestingly, from the $3.3 \times 10^6 \cap 5 \times 10^5$ intersection of V1 and V2 populations, only 112 sequences were common to two sets.

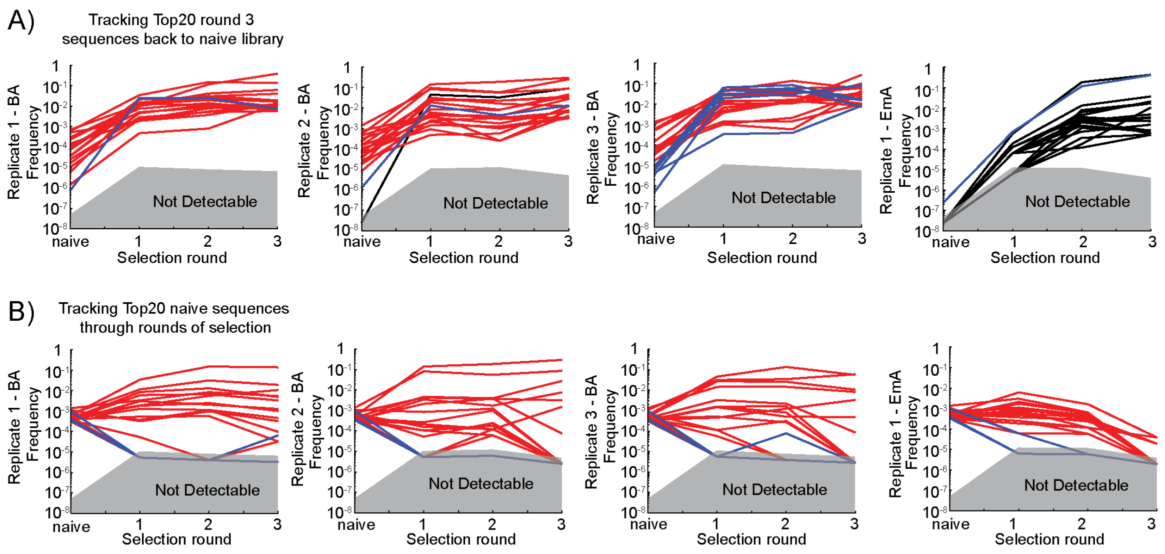


**Figure S3**. Tracing top 20 sequences through selection. (A) Top panels trace the origin of the top 20 sequences from the third round of selection to the naïve library. The colors for each line indicate the identity of each sequence (red – 'parasite'; blue – 'visible'; black – 'invisible'). Parasites were enriched when in the BA selection method, while sequences from the I-population were enriched when using the EmA selection. (B) Bottom panels trace the fate of the top 20 sequences in the naïve library through the selection rounds. In EmA selection is used, all top 20 sequences in the naïve library were depleted by the third round of selection. In BA selection, many sequences from the naïve library persist through rounds 1-3.

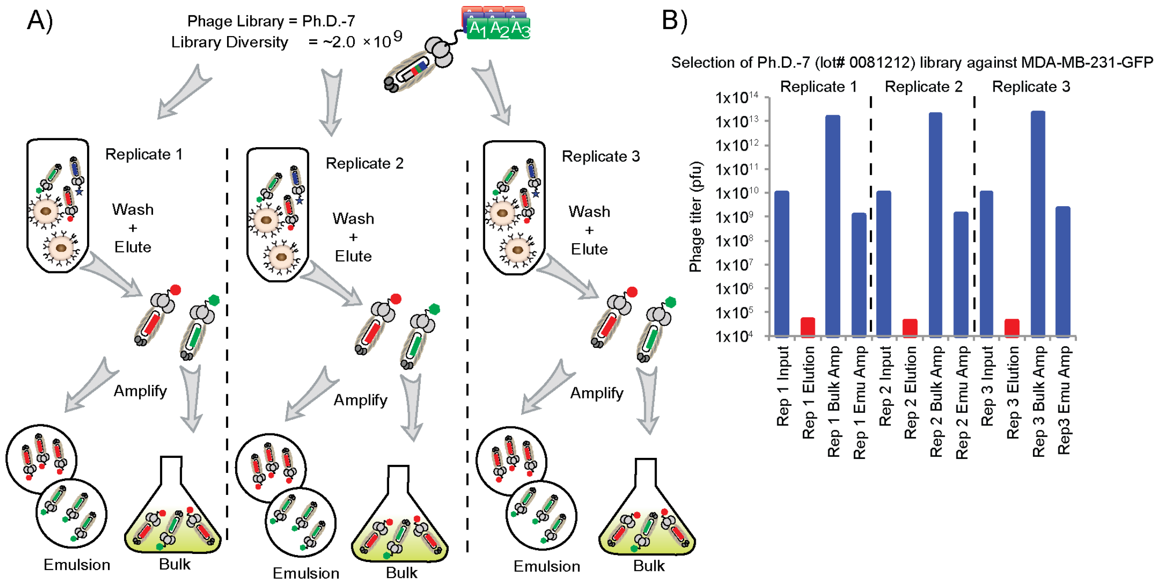


**Figure S4.** Monitoring phage titers in selection with Ph.D.-7 library lot #2. (A) Scheme of the selection performed with lot #2 of the Ph.D.-7 library. The three replicates of panning against MDA-MB-231-GFP cells. The eluted phage mixture was separated into two samples and amplified either by BA or EmA. (B) Summary of the input, output and amplified phage titers in all replicates of selection using the second lot of the PhD-7 library.

| L1-BA-R1-r1 | L1-BA-R1-r2 | L1-BA-R1-r3 | L1-BA-R2-r1 | L1-BA-R2-r2 | L1-BA-R2-r3 | L1-BA-R3-r1 | L1-BA-R3-r2 | L1-BA-R3-r3 |
|---|---|---|---|---|---|---|---|---|
| LOT 1 | LOT 1 | LOT 1 | LOT 1 | LOT 1 | LOT 1 | LOT 1 | LOT 1 | LOT 1 |
| BA-screen | BA-screen | BA-screen | BA-screen | BA-screen | BA-screen | BA-screen | BA-screen | BA-screen |
| Replicate 1 | Replicate 1 | Replicate 1 | Replicate 2 | Replicate 2 | Replicate 2 | Replicate 3 | Replicate 3 | Replicate 3 |
| Round 1 | Round 2 | Round 3 | Round 1 | Round 2 | Round 3 | Round 1 | Round 2 | Round 3 |
| Sequence | Sequence | Sequence | Sequence | Sequence | Sequence | Sequence | Sequence | Sequence |
| STASYTR | STASYTR | GETRAPL | HAIYPRH | HAIYPRH | HAIYPRH | MGLQTPY | STASYTR | SYHSFNL |
| QNTTTAL | GETRAPL | STASYTR | QPPRSTS | QPPRSTS | QPPRSTS | SILPYPY | MGLQTPY | APRTFNQ |
| MPGSLPS | YAGPYQH | YAGPYQH | GKPMPPM | GKPMPPM | QPTHPTR | STASYTR | HSTKVAF | TGHSAQG |
| TPQSSPT | EPLQLKM | YLTMPTP | QPTHPTR | QPTHPTR | GKPMPPM | NQLPLHA | SILPYPY | SILPYPY |
| QEPLTAR | YLTMPTP | SPWDARL | IPTLPSS | VTAHGGR | ALAHRIL | HSTKVAF | IPAPLRS | QPWPTSI |
| SPWDARL | SPWDARL | EPLQLKM | VTAHGGR | NHWASPR | QPSMLNP | MDAHHAL | NQLPLHA | STASYTR |
| HFRSGSL | QNTTTAL | VIPHVLS | NHWASPR | TWYFGPL | NHWASPR | QPWPTSI | QPWPTSI | HTIQFTP |
| VIPHVLS | TPQSSPT | SILPYPY | QPSMLNP | IPTLPSS | VTAHGGR | IPAPLRS | MDAHHAL | FPSTITP |
| GVKALST | VIPHVLS | YAAHRSH | TWYFGPL | YAGPYQH | SPTQPKS | HAIYPRH | TGHSAQG | ASYSGTA |
| ANTTPRH | QEPLTAR | QALSVYR | STPMQNL | QPSMLNP | STPMQNL | QAHTVGK | QAHTVGK | SPTGWAP |
| YAGPYQH | ANTTPRH | HAIYPRH | MDAHHAL | ALAHRIL | HHSLTVT | TGHSAQG | HAIYPRH | NQLPLHA |
| EPLQLKM | GVKALST | MPKYYLQ | YAGPYQH | MDAHHAL | IPTLPSS | MPTLTPT | SLHQPHL | SHSLLHH |
| TVRHLQL | DSHTPQR | GVKALST | KAVHPLR | STPMQNL | VLPGRSP | SLHQPHL | STTKLAL | STFTKSP |
| QTSMATV | SPQMTLS | ANTTPRH | QSLALQP | SPTQPKS | TARYPSW | MPRTPTD | SPTGWAP | MGLQTPY |
| GETRAPL | TTNLSPW | HFRSGSL | ALAHRIL | AMSSRSL | MHAPPFY | GKPMPPM | GKPMPPM | IPAPLRS |
| YLTMPTP | MPKYYLQ | QNTTTAL | SHTAPLR | HALGPSS | QLMNASR | SPTGWAP | MPRTPTD | HAIYPRH |
| QRLPQTA | TVRHLQL | TKTDTWL | SLSLIQT | MHAPPFY | STFTKSP | TLLPFQP | APRTFNQ | STPIQQP |
| SPQMTLS | HFRSGSL | DSHTPQR | GETRAPL | SLSLIQT | SLSLIQT | FPSTITP | STFTKSP | SHHQKPP |
| TTNLSPW | HIPPGSP | TPQSSPT | VIPHVLS | SSLVRTA | HALGPSS | AGNGTTP | SYHSFNL | GKPMPPM |
| KAVHPLR | QLHNDAT | SSLPLRK | TARYPSW | WSPHGLA | TPPTMDH | NAEQIAP | SHSLLHH | HSTKVAF |

| L1-EmA-R1-r1 | L1-EmA-R1-r2 | L1-EmA-R1-r3 | L2-EmA-R1-r1 | L2-EmA-R2-r1 | L2-EmA-R3-r1 | L2-BA-R1-r1 | L2-BA-R2-r1 | L1-BA-R3-r1 |
|---|---|---|---|---|---|---|---|---|
| LOT 1 | LOT 1 | LOT 1 | LOT 2 | LOT 2 | LOT 2 | LOT 2 | LOT 2 | LOT 2 |
| EmA-screen | EmA-screen | EmA-screen | EmA-screen | EmA-screen | EmA-screen | BA-screen | BA-screen | BA-screen |
| Replicate 1 | Replicate 1 | Replicate 1 | Replicate 1 | Replicate 2 | Replicate 3 | Replicate 1 | Replicate 2 | Replicate 3 |
| Round 1 | Round 2 | Round 3 | Round 1 | Round 1 | Round 1 | Round 1 | Round 1 | Round 1 |
| Sequence | Sequence | Sequence | Sequence | Sequence | Sequence | Sequence | Sequence | Sequence |
| HAIYPRH | SWQYGKL | SWQYGKL | SWQYGKL | HAIYPRH | EQGRPLP | STPATLI | AGSVIDT | HSRAPER |
| GKPMPPM | NQLAGSG | NQLAGSG | NQLAGSG | YLTMPTP | MAANGAR | WSLSELH | QAYHVSA | TTLGVWT |
| GPMLARG | HWHFGPL | HWHFGPL | TSSESES | STASYTR | QVLLTAA | LPVRLDW | SNMTRWH | YSEPAVT |
| AMSSRSL | TYRFGPL | TYKFGTL | ERTVLHT | TPQSSPT | AGRELCC | QTWLEMG | GRLDTGI | ESRVMSR |
| SSALLLP | SWKFGPL | TYRFGPL | TYKFGTL | EPLQLKM | ARAVLQL | GPHNPTQ | ALQPQKH | GPLHAQF |
| DSHTPQR | ALEVTFW | SWKFGPL | HWHFGPL | GVKALST | QNMQQQI | NDRPHMP | NAYGGRI | NNTLSRT |
| AASSLTI | TYKFGTL | HWKFGIL | TYRFGPL | TPFMAYH | AWSAVMR | VPNIVTQ | TKTVLER | DHAVPRY |
| QPPRSTS | VQNEWRS | TFKFGPL | RFTVDWD | IPAPLRS | APIWMHV | LRSDPVV | GWETRME | AFPPVTA |
| HALGPSS | LTVEPWL | TWKFSPL | SWKFGPL | RLPSWHE | ATWQLGT | VPASPWT | WNQRATG | MTVQRGP |
| STASYTR | ELWVSPL | TYKYYPL | TLTVQAW | AYPEPYV | LHRQSSA | SSVSWLN | VDMIVPS | HLNQQNH |
| EPLQLKM | LEVYALV | TYLFQPL | LAGPLMT | QPTHPTR | ASWIPLP | TTQVLEA | LPGNRLL | QLPFTIK |
| SSSVVTH | TYKYYPL | TYQYGKL | TWKFSPL | SWQYGKL | QQQYMAH | QFTQLHQ | QLYREFN | SQPTWMF |
| QATHRSH | TFKFGPL | LTVEPWL | NVSGSHS | QPPRSTS | STAMDGR | DAIPTSV | GTSTTAQ | YNGSANQ |
| GKVQAQS | TYQYGKL | HWKYWPL | SVLLPHR | SILPYPY | GWRTTWP | VGKTSFQ | NTQLHPS | TTQVLEA |
| SILPYPY | TYLFQPL | TYVFYPL | DAGQVSQ | TQTMRST | ATHQRPA | SLDVRMW | YRNHVTY | FSLQTTR |
| SPTQPKS | DLTVTPW | DLTVTPW | TYKYYPL | TKTDTWL | WMASMAV | GQVALLD | MNSNIPI | GLRNPPS |
| TYQNPVH | TWKFSPL | MEVFPYY | HHQYVPA | TPMTRAL | HEQPMHR | VENVHVR | ATLVPAA | TEKFRVT |
| ANTTPRH | QLTVMSW | ELWVSPL | WTTTSRL | VMSQPHP | SLDVRMW | VPVTMYW | ILAHSIM | TVISQNM |
| QTGYATR | MTVQPWP | KVWELHP | QLMPMMM | SPWDARL | QHMPLTR | ELGTTQT | IDGNGTH | ASSMPTQ |
| GETRAPL | HAIYPRH | TYRFLPL | RPYDTAH | GSTVFTA | TMTEHRQ | AMTALDL | IDNSHTH | AFTTSYM |

Legend L#: Lot Number BA: Bulk Amplification EmA: Emulsion Amplification R#: Replicate Number r#: Round Number

**Table S1**. List of the top 20 sequences from each round and replicate of selection using both BA and EmA methods. The abbreviation and color for each selection is used to distinguish the origin of hits in Figures S4, S6-S10.

### Table S2

| | Lot1-BA-Rep1-Round1 | Lot1-BA-Rep1-Round2 | Lot1-BA-Rep1-Round3 | Lot1-BA-Rep2-Round1 | Lot1-BA-Rep2-Round2 | Lot1-BA-Rep2-Round3 | Lot1-BA-Rep3-Round1 | Lot1-BA-Rep3-Round2 | Lot1-BA-Rep3-Round3 | Lot1-EmA-Rep1-Round1 | Lot1-EmA-Rep1-Round2 | Lot1-EmA-Rep1-Round3 | Lot2-EmA-Rep1-Round1 | Lot2-EmA-Rep2-Round1 | Lot2-EmA-Rep3-Round1 | Lot2-BA-Rep1-Round1 | Lot2-BA-Rep2-Round1 | Lot2-BA-Rep3-Round1 | Lot1-Parasites | Lot2-Parasites |
|---|---|---|---|---|---|---|---|---|---|---|---|---|---|---|---|---|---|---|---|---|
| STASYTR | 645 | 3722 | 4203 | 1 | 4 | 0 | 807 | 3490 | 1957 | 17 | 6 | 1 | 38 | 1695 | 53 | 0 | 0 | 0 | 1 | 0 |
| QNTTTAL | 482 | 629 | 229 | 0 | 0 | 0 | 0 | 0 | 0 | 3 | 0 | 0 | 7 | 575 | 32 | 0 | 0 | 0 | 0 | 0 |
| LPGSLPS | 416 | 143 | 23 | 4 | 0 | 0 | 0 | 0 | 0 | 0 | 0 | 0 | 0 | 151 | 38 | 0 | 0 | 0 | 1 | 0 |
| TPQSSPT | 400 | 612 | 207 | 0 | 0 | 0 | 0 | 0 | 0 | 3 | 8 | 1 | 22 | 1408 | 50 | 0 | 0 | 0 | 1 | 0 |
| QEPLTAR | 317 | 549 | 97 | 0 | 1 | 0 | 0 | 0 | 0 | 0 | 0 | 0 | 10 | 413 | 5 | 0 | 0 | 0 | 0 | 0 |
| SPWDARL | 281 | 705 | 593 | 3 | 6 | 0 | 0 | 1 | 0 | 1 | 0 | 0 | 41 | 697 | 39 | 0 | 0 | 0 | 1 | 0 |
| HFRSGSL | 263 | 266 | 264 | 0 | 0 | 0 | 0 | 0 | 0 | 4 | 0 | 1 | 0 | 563 | 12 | 0 | 0 | 0 | 1 | 0 |
| VIPHVLS | 262 | 569 | 503 | 101 | 29 | 7 | 1 | 0 | 0 | 4 | 0 | 0 | 27 | 207 | 44 | 0 | 0 | 0 | 1 | 0 |
| GVKALST | 236 | 497 | 295 | 42 | 7 | 22 | 0 | 0 | 0 | 6 | 1 | 0 | 5 | 1139 | 10 | 0 | 0 | 0 | 1 | 0 |
| ANTTPRH | 236 | 523 | 289 | 0 | 0 | 0 | 2 | 8 | 1 | 12 | 0 | 3 | 0 | 475 | 32 | 0 | 0 | 0 | 1 | 0 |
| YAGPYQH | 215 | 1028 | 1907 | 190 | 141 | 63 | 1 | 2 | 0 | 4 | 2 | 0 | 4 | 437 | 16 | 0 | 0 | 0 | 1 | 0 |
| EPLQLKM | 214 | 777 | 586 | 3 | 2 | 0 | 2 | 0 | 0 | 16 | 6 | 0 | 7 | 1159 | 6 | 1 | 0 | 0 | 1 | 0 |
| TVRHLQL | 213 | 273 | 124 | 9 | 3 | 0 | 0 | 0 | 0 | 2 | 0 | 0 | 5 | 266 | 8 | 0 | 0 | 0 | 1 | 0 |
| QTSMATV | 200 | 203 | 47 | 35 | 5 | 0 | 0 | 0 | 0 | 2 | 0 | 0 | 0 | 0 | 22 | 0 | 0 | 0 | 1 | 0 |
| GETRAPL | 191 | 2868 | 11859 | 111 | 36 | 1 | 5 | 0 | 0 | 11 | 6 | 14 | 29 | 478 | 40 | 0 | 0 | 0 | 1 | 0 |
| YLTMPTP | 191 | 749 | 1254 | 4 | 4 | 0 | 71 | 51 | 0 | 7 | 4 | 1 | 18 | 2296 | 15 | 0 | 0 | 0 | 1 | 0 |
| QRLPQTA | 166 | 178 | 21 | 1 | 1 | 0 | 31 | 27 | 0 | 2 | 2 | 0 | 0 | 73 | 0 | 0 | 0 | 0 | 1 | 0 |
| SPQMTLS | 159 | 322 | 151 | 5 | 16 | 0 | 58 | 56 | 3 | 10 | 6 | 0 | 0 | 7 | 4 | 0 | 0 | 0 | 1 | 0 |
| TTNLSPW | 157 | 309 | 147 | 0 | 0 | 0 | 0 | 0 | 0 | 1 | 0 | 0 | 0 | 4 | 0 | 0 | 0 | 0 | 1 | 0 |
| KAVHPLR | 152 | 220 | 118 | 173 | 32 | 39 | 0 | 0 | 0 | 2 | 0 | 0 | 6 | 133 | 20 | 0 | 0 | 0 | 1 | 0 |
| STASYTR | 645 | 3722 | 4203 | 1 | 4 | 0 | 807 | 3490 | 1957 | 17 | 6 | 1 | 38 | 1695 | 53 | 0 | 0 | 0 | 1 | 0 |
| GETRAPL | 191 | 2868 | 11859 | 111 | 36 | 1 | 5 | 0 | 0 | 11 | 6 | 14 | 29 | 478 | 40 | 0 | 0 | 0 | 1 | 0 |
| YAGPYQH | 215 | 1028 | 1907 | 190 | 141 | 63 | 1 | 2 | 0 | 4 | 2 | 0 | 4 | 437 | 16 | 0 | 0 | 0 | 1 | 0 |
| EPLQLKM | 214 | 777 | 586 | 3 | 2 | 0 | 2 | 0 | 0 | 16 | 6 | 0 | 7 | 1159 | 6 | 1 | 0 | 0 | 1 | 0 |
| YLTMPTP | 191 | 749 | 1254 | 4 | 4 | 0 | 71 | 51 | 0 | 7 | 4 | 1 | 18 | 2296 | 15 | 0 | 0 | 0 | 1 | 0 |
| SPWDARL | 281 | 705 | 593 | 3 | 6 | 0 | 0 | 1 | 0 | 1 | 0 | 0 | 41 | 697 | 39 | 0 | 0 | 0 | 1 | 0 |
| QNTTTAL | 482 | 629 | 229 | 0 | 0 | 0 | 0 | 0 | 0 | 3 | 0 | 0 | 7 | 575 | 32 | 0 | 0 | 0 | 0 | 0 |
| TPQSSPT | 400 | 612 | 207 | 0 | 0 | 0 | 0 | 0 | 0 | 3 | 8 | 1 | 22 | 1408 | 50 | 0 | 0 | 0 | 1 | 0 |
| VIPHVLS | 262 | 569 | 503 | 101 | 29 | 7 | 1 | 0 | 0 | 4 | 0 | 0 | 27 | 207 | 44 | 0 | 0 | 0 | 1 | 0 |
| QEPLTAR | 317 | 549 | 97 | 0 | 1 | 0 | 0 | 0 | 0 | 0 | 0 | 0 | 10 | 413 | 5 | 0 | 0 | 0 | 0 | 0 |
| ANTTPRH | 236 | 523 | 289 | 0 | 0 | 0 | 2 | 8 | 1 | 12 | 0 | 3 | 0 | 475 | 32 | 0 | 0 | 0 | 1 | 0 |
| GVKALST | 236 | 497 | 295 | 42 | 7 | 22 | 0 | 0 | 0 | 6 | 1 | 0 | 5 | 1139 | 10 | 0 | 0 | 0 | 1 | 0 |
| DSHTPQR | 141 | 349 | 208 | 12 | 9 | 0 | 11 | 24 | 3 | 21 | 8 | 0 | 0 | 4 | 0 | 0 | 0 | 0 | 1 | 0 |
| SPQMTLS | 159 | 322 | 151 | 5 | 16 | 0 | 58 | 56 | 3 | 10 | 6 | 0 | 0 | 7 | 4 | 0 | 0 | 0 | 1 | 0 |
| TTNLSPW | 157 | 309 | 147 | 0 | 0 | 0 | 0 | 0 | 0 | 1 | 0 | 0 | 0 | 4 | 0 | 0 | 0 | 0 | 1 | 0 |
| MPKYYLQ | 85 | 284 | 327 | 16 | 8 | 0 | 0 | 0 | 0 | 0 | 0 | 0 | 0 | 560 | 0 | 0 | 0 | 0 | 1 | 0 |
| TVRHLQL | 213 | 273 | 124 | 9 | 3 | 0 | 0 | 0 | 0 | 2 | 0 | 0 | 5 | 266 | 8 | 0 | 0 | 0 | 1 | 0 |
| HFRSGSL | 263 | 266 | 264 | 0 | 0 | 0 | 0 | 0 | 0 | 4 | 0 | 1 | 0 | 563 | 12 | 0 | 0 | 0 | 1 | 0 |
| HIPPGSP | 134 | 264 | 123 | 12 | 2 | 0 | 15 | 10 | 0 | 3 | 1 | 1 | 16 | 38 | 0 | 0 | 0 | 0 | 1 | 0 |
| QLHNDAT | 133 | 239 | 89 | 0 | 0 | 1 | 0 | 0 | 0 | 2 | 0 | 0 | 9 | 23 | 0 | 0 | 0 | 0 | 1 | 0 |
| GETRAPL | 191 | 2868 | 11859 | 111 | 36 | 1 | 5 | 0 | 0 | 11 | 6 | 14 | 29 | 478 | 40 | 0 | 0 | 0 | 1 | 0 |
| STASYTR | 645 | 3722 | 4203 | 1 | 4 | 0 | 807 | 3490 | 1957 | 17 | 6 | 1 | 38 | 1695 | 53 | 0 | 0 | 0 | 1 | 0 |
| YAGPYQH | 215 | 1028 | 1907 | 190 | 141 | 63 | 1 | 2 | 0 | 4 | 2 | 0 | 4 | 437 | 16 | 0 | 0 | 0 | 1 | 0 |
| YLTMPTP | 191 | 749 | 1254 | 4 | 4 | 0 | 71 | 51 | 0 | 7 | 4 | 1 | 18 | 2296 | 15 | 0 | 0 | 0 | 1 | 0 |
| SPWDARL | 281 | 705 | 593 | 3 | 6 | 0 | 0 | 1 | 0 | 1 | 0 | 0 | 41 | 697 | 39 | 0 | 0 | 0 | 1 | 0 |
| EPLQLKM | 214 | 777 | 586 | 3 | 2 | 0 | 2 | 0 | 0 | 16 | 6 | 0 | 7 | 1159 | 6 | 1 | 0 | 0 | 1 | 0 |
| VIPHVLS | 262 | 569 | 503 | 101 | 29 | 7 | 1 | 0 | 0 | 4 | 0 | 0 | 27 | 207 | 44 | 0 | 0 | 0 | 1 | 0 |
| SILPYPY | 36 | 92 | 409 | 16 | 10 | 12 | 857 | 1488 | 2536 | 13 | 8 | 1 | 10 | 847 | 22 | 0 | 0 | 0 | 1 | 0 |
| YAAHRSH | 9 | 20 | 390 | 0 | 0 | 0 | 0 | 0 | 0 | 1 | 0 | 0 | 0 | 0 | 0 | 0 | 0 | 0 | 1 | 0 |
| QALSVYR | 42 | 192 | 380 | 0 | 1 | 1 | 0 | 0 | 0 | 2 | 4 | 0 | 20 | 17 | 7 | 0 | 0 | 0 | 1 | 0 |
| HAIYPRH | 35 | 137 | 328 | 2659 | 3032 | 11887 | 481 | 661 | 370 | 99 | 28 | 2 | 148 | 8381 | 19 | 0 | 0 | 10 | 1 | 0 |
| MPKYYLQ | 85 | 284 | 327 | 16 | 8 | 0 | 0 | 0 | 0 | 0 | 0 | 0 | 0 | 560 | 0 | 0 | 0 | 0 | 1 | 0 |
| GVKALST | 236 | 497 | 295 | 42 | 7 | 22 | 0 | 0 | 0 | 6 | 1 | 0 | 5 | 1139 | 10 | 0 | 0 | 0 | 1 | 0 |
| ANTTPRH | 236 | 523 | 289 | 0 | 0 | 0 | 2 | 8 | 1 | 12 | 0 | 3 | 0 | 475 | 32 | 0 | 0 | 0 | 1 | 0 |
| HFRSGSL | 263 | 266 | 264 | 0 | 0 | 0 | 0 | 0 | 0 | 4 | 0 | 1 | 0 | 563 | 12 | 0 | 0 | 0 | 1 | 0 |
| QNTTTAL | 482 | 629 | 229 | 0 | 0 | 0 | 0 | 0 | 0 | 3 | 0 | 0 | 7 | 575 | 32 | 0 | 0 | 0 | 0 | 0 |
| TKTDTWL | 95 | 216 | 219 | 2 | 1 | 0 | 3 | 2 | 0 | 11 | 5 | 1 | 21 | 725 | 34 | 0 | 0 | 0 | 1 | 0 |
| DSHTPQR | 141 | 349 | 208 | 12 | 9 | 0 | 11 | 24 | 3 | 21 | 8 | 0 | 0 | 4 | 0 | 0 | 0 | 0 | 1 | 0 |
| TPQSSPT | 400 | 612 | 207 | 0 | 0 | 0 | 0 | 0 | 0 | 3 | 8 | 1 | 22 | 1408 | 50 | 0 | 0 | 0 | 1 | 0 |
| SSLPLRK | 53 | 130 | 175 | 5 | 2 | 0 | 0 | 0 | 0 | 2 | 1 | 0 | 0 | 474 | 15 | 0 | 0 | 0 | 1 | 0 |
| HAIYPRH | 35 | 137 | 328 | 2659 | 3032 | 11887 | 481 | 661 | 370 | 99 | 28 | 2 | 148 | 8381 | 19 | 0 | 0 | 10 | 1 | 0 |
| QPPRSTS | 23 | 33 | 11 | 1775 | 938 | 10032 | 8 | 4 | 0 | 19 | 5 | 0 | 0 | 847 | 13 | 0 | 0 | 0 | 1 | 0 |
| GKPMPPM | 63 | 208 | 165 | 1540 | 899 | 3479 | 254 | 375 | 287 | 49 | 11 | 0 | 3 | 24 | 0 | 0 | 0 | 0 | 1 | 0 |
| QPTHPTR | 0 | 0 | 0 | 827 | 524 | 3585 | 3 | 0 | 0 | 0 | 0 | 0 | 0 | 889 | 0 | 0 | 0 | 0 | 0 | 0 |
| IPTLPSS | 1 | 3 | 8 | 589 | 259 | 485 | 86 | 190 | 221 | 4 | 2 | 0 | 7 | 239 | 0 | 0 | 0 | 4 | 1 | 0 |
| VTAHGGR | 0 | 0 | 3 | 524 | 367 | 1240 | 2 | 1 | 5 | 1 | 0 | 0 | 0 | 0 | 0 | 0 | 0 | 0 | 1 | 0 |
| NHWASPR | 0 | 0 | 0 | 443 | 365 | 1431 | 72 | 189 | 17 | 9 | 2 | 0 | 0 | 233 | 0 | 0 | 0 | 0 | 1 | 0 |
| QPSMLNP | 2 | 0 | 0 | 348 | 132 | 1457 | 74 | 87 | 121 | 0 | 0 | 0 | 0 | 0 | 0 | 0 | 0 | 0 | 1 | 0 |
| TWYFGPL | 0 | 0 | 0 | 297 | 364 | 0 | 76 | 30 | 0 | 0 | 1 | 0 | 6 | 0 | 16 | 0 | 0 | 0 | 0 | 0 |
| STPMQNL | 1 | 0 | 0 | 242 | 71 | 528 | 0 | 0 | 0 | 0 | 0 | 0 | 0 | 0 | 0 | 0 | 0 | 0 | 0 | 0 |
| MDAHHAL | 2 | 0 | 0 | 235 | 97 | 8 | 702 | 870 | 4 | 10 | 3 | 0 | 0 | 0 | 0 | 0 | 0 | 0 | 1 | 0 |
| YAGPYQH | 215 | 1028 | 1907 | 190 | 141 | 63 | 1 | 2 | 0 | 4 | 2 | 0 | 4 | 437 | 16 | 0 | 0 | 0 | 1 | 0 |
| KAVHPLR | 152 | 220 | 118 | 173 | 32 | 39 | 0 | 0 | 0 | 2 | 0 | 0 | 6 | 133 | 20 | 0 | 0 | 0 | 1 | 0 |
| QSLALQP | 9 | 9 | 1 | 163 | 20 | 1 | 2 | 0 | 0 | 0 | 0 | 0 | 0 | 0 | 0 | 0 | 0 | 0 | 1 | 0 |
| ALAHRIL | 0 | 0 | 0 | 156 | 130 | 1817 | 0 | 1 | 0 | 4 | 1 | 0 | 0 | 0 | 0 | 0 | 0 | 0 | 1 | 0 |
| SHTAPLR | 92 | 144 | 97 | 112 | 19 | 5 | 1 | 0 | 0 | 3 | 2 | 0 | 27 | 0 | 0 | 0 | 0 | 0 | 1 | 0 |
| SLSLIQT | 2 | 0 | 0 | 112 | 46 | 136 | 0 | 2 | 0 | 1 | 1 | 0 | 0 | 0 | 0 | 0 | 0 | 0 | 1 | 0 |
| GETRAPL | 191 | 2868 | 11859 | 111 | 36 | 1 | 5 | 0 | 0 | 11 | 6 | 14 | 29 | 478 | 40 | 0 | 0 | 0 | 1 | 0 |
| VIPHVLS | 262 | 569 | 503 | 101 | 29 | 7 | 1 | 0 | 0 | 4 | 0 | 0 | 27 | 207 | 44 | 0 | 0 | 0 | 1 | 0 |
| TARYPSW | 78 | 147 | 33 | 89 | 4 | 301 | 19 | 5 | 0 | 2 | 0 | 0 | 0 | 0 | 0 | 0 | 0 | 0 | 1 | 0 |

### Table S2 (cont.)

| | Lot1-BA-Rep1-Round1 | Lot1-BA-Rep1-Round2 | Lot1-BA-Rep1-Round3 | Lot1-BA-Rep2-Round1 | Lot1-BA-Rep2-Round2 | Lot1-BA-Rep2-Round3 | Lot1-BA-Rep3-Round1 | Lot1-BA-Rep3-Round2 | Lot1-BA-Rep3-Round3 | Lot1-EmA-Rep1-Round1 | Lot1-EmA-Rep1-Round2 | Lot1-EmA-Rep1-Round3 | Lot2-EmA-Rep1-Round1 | Lot2-EmA-Rep2-Round1 | Lot2-EmA-Rep3-Round1 | Lot2-BA-Rep1-Round1 | Lot2-BA-Rep2-Round1 | Lot2-BA-Rep3-Round1 | Lot1-Parasites | Lot2-Parasites |
|---|---|---|---|---|---|---|---|---|---|---|---|---|---|---|---|---|---|---|---|---|
| HAIYPRH | 35 | 137 | 328 | 2659 | 3032 | 11887 | 481 | 661 | 370 | 99 | 28 | 2 | 148 | 8381 | 19 | 0 | 0 | 10 | 1 | 0 |
| QPPRSTS | 23 | 33 | 11 | 1775 | 938 | 10032 | 8 | 4 | 0 | 19 | 5 | 0 | 0 | 847 | 13 | 0 | 0 | 0 | 1 | 0 |
| GKPMPPM | 63 | 208 | 165 | 1540 | 899 | 3479 | 254 | 375 | 287 | 49 | 11 | 0 | 3 | 24 | 0 | 0 | 0 | 0 | 1 | 0 |
| QPTHPTR | 0 | 0 | 0 | 827 | 524 | 3585 | 3 | 0 | 0 | 0 | 0 | 0 | 0 | 889 | 0 | 0 | 0 | 0 | 0 | 0 |
| VTAHGGR | 0 | 0 | 3 | 524 | 367 | 1240 | 2 | 1 | 5 | 1 | 0 | 0 | 0 | 0 | 0 | 0 | 0 | 0 | 1 | 0 |
| NHWASPR | 0 | 0 | 0 | 443 | 365 | 1431 | 72 | 189 | 17 | 9 | 2 | 0 | 0 | 233 | 0 | 0 | 0 | 0 | 1 | 0 |
| TWYFGPL | 0 | 0 | 0 | 297 | 364 | 0 | 76 | 30 | 0 | 0 | 1 | 0 | 6 | 0 | 16 | 0 | 0 | 0 | 0 | 0 |
| IPTLPSS | 1 | 3 | 8 | 589 | 259 | 485 | 86 | 190 | 221 | 4 | 2 | 0 | 7 | 239 | 0 | 0 | 0 | 4 | 1 | 0 |
| YAGPYQH | 215 | 1028 | 1907 | 190 | 141 | 63 | 1 | 2 | 0 | 4 | 2 | 0 | 4 | 437 | 16 | 0 | 0 | 0 | 1 | 0 |
| QPSMLNP | 2 | 0 | 0 | 348 | 132 | 1457 | 74 | 87 | 121 | 0 | 0 | 0 | 0 | 0 | 0 | 0 | 0 | 0 | 1 | 0 |
| ALAHRIL | 0 | 0 | 0 | 156 | 130 | 1817 | 0 | 1 | 0 | 4 | 1 | 0 | 0 | 0 | 0 | 0 | 0 | 0 | 1 | 0 |
| MDAHHAL | 2 | 0 | 0 | 235 | 97 | 8 | 702 | 870 | 4 | 10 | 3 | 0 | 0 | 0 | 0 | 0 | 0 | 0 | 1 | 0 |
| STPMQNL | 1 | 0 | 0 | 242 | 71 | 528 | 0 | 0 | 0 | 0 | 0 | 0 | 0 | 0 | 0 | 0 | 0 | 0 | 0 | 0 |
| SPTQPKS | 34 | 61 | 31 | 88 | 64 | 1120 | 9 | 7 | 0 | 13 | 2 | 0 | 0 | 0 | 0 | 0 | 0 | 0 | 1 | 0 |
| AMSSRSL | 0 | 0 | 0 | 40 | 61 | 3 | 8 | 32 | 117 | 28 | 10 | 0 | 14 | 245 | 0 | 0 | 0 | 0 | 1 | 0 |
| HALGPSS | 7 | 27 | 3 | 81 | 60 | 127 | 0 | 2 | 0 | 18 | 8 | 0 | 0 | 0 | 10 | 0 | 0 | 0 | 1 | 0 |
| MHAPPFY | 0 | 0 | 1 | 45 | 52 | 249 | 2 | 1 | 0 | 1 | 0 | 0 | 0 | 0 | 0 | 0 | 0 | 0 | 1 | 0 |
| SLSLIQT | 2 | 0 | 0 | 112 | 46 | 136 | 0 | 2 | 0 | 1 | 1 | 0 | 0 | 0 | 0 | 0 | 0 | 0 | 1 | 0 |
| SSLVRTA | 13 | 38 | 35 | 40 | 44 | 32 | 26 | 33 | 0 | 1 | 1 | 0 | 0 | 209 | 22 | 0 | 0 | 0 | 1 | 0 |
| WSPHGLA | 0 | 0 | 0 | 36 | 43 | 0 | 10 | 4 | 0 | 0 | 0 | 0 | 0 | 0 | 0 | 0 | 0 | 0 | 0 | 0 |
| HAIYPRH | 35 | 137 | 328 | 2659 | 3032 | 11887 | 481 | 661 | 370 | 99 | 28 | 2 | 148 | 8381 | 19 | 0 | 0 | 10 | 1 | 0 |
| QPPRSTS | 23 | 33 | 11 | 1775 | 938 | 10032 | 8 | 4 | 0 | 19 | 5 | 0 | 0 | 847 | 13 | 0 | 0 | 0 | 1 | 0 |
| QPTHPTR | 0 | 0 | 0 | 827 | 524 | 3585 | 3 | 0 | 0 | 0 | 0 | 0 | 0 | 889 | 0 | 0 | 0 | 0 | 0 | 0 |
| GKPMPPM | 63 | 208 | 165 | 1540 | 899 | 3479 | 254 | 375 | 287 | 49 | 11 | 0 | 3 | 24 | 0 | 0 | 0 | 0 | 1 | 0 |
| ALAHRIL | 0 | 0 | 0 | 156 | 130 | 1817 | 0 | 1 | 0 | 4 | 1 | 0 | 0 | 0 | 0 | 0 | 0 | 0 | 1 | 0 |
| QPSMLNP | 2 | 0 | 0 | 348 | 132 | 1457 | 74 | 87 | 121 | 0 | 0 | 0 | 0 | 0 | 0 | 0 | 0 | 0 | 1 | 0 |
| NHWASPR | 0 | 0 | 0 | 443 | 365 | 1431 | 72 | 189 | 17 | 9 | 2 | 0 | 0 | 233 | 0 | 0 | 0 | 0 | 1 | 0 |
| VTAHGGR | 0 | 0 | 3 | 524 | 367 | 1240 | 2 | 1 | 5 | 1 | 0 | 0 | 0 | 0 | 0 | 0 | 0 | 0 | 1 | 0 |
| SPTQPKS | 34 | 61 | 31 | 88 | 64 | 1120 | 9 | 7 | 0 | 13 | 2 | 0 | 0 | 0 | 0 | 0 | 0 | 0 | 1 | 0 |
| STPMQNL | 1 | 0 | 0 | 242 | 71 | 528 | 0 | 0 | 0 | 0 | 0 | 0 | 0 | 0 | 0 | 0 | 0 | 0 | 0 | 0 |
| HHSLTVT | 0 | 0 | 2 | 9 | 9 | 492 | 0 | 0 | 0 | 2 | 0 | 0 | 0 | 0 | 0 | 0 | 0 | 0 | 1 | 0 |
| IPTLPSS | 1 | 3 | 8 | 589 | 259 | 485 | 86 | 190 | 221 | 4 | 2 | 0 | 7 | 239 | 0 | 0 | 0 | 4 | 1 | 0 |
| VLPGRSP | 0 | 0 | 0 | 65 | 28 | 309 | 0 | 0 | 0 | 1 | 0 | 0 | 0 | 0 | 0 | 0 | 0 | 0 | 1 | 0 |
| TARYPSW | 78 | 147 | 33 | 89 | 4 | 301 | 19 | 5 | 0 | 2 | 0 | 0 | 0 | 0 | 0 | 0 | 0 | 0 | 1 | 0 |
| MHAPPFY | 0 | 0 | 1 | 45 | 52 | 249 | 2 | 1 | 0 | 1 | 0 | 0 | 0 | 0 | 0 | 0 | 0 | 0 | 1 | 0 |
| QLMNASR | 0 | 0 | 0 | 17 | 4 | 167 | 2 | 0 | 0 | 2 | 2 | 0 | 0 | 0 | 0 | 0 | 0 | 0 | 1 | 0 |
| STFTKSP | 1 | 0 | 4 | 27 | 20 | 141 | 83 | 335 | 626 | 7 | 3 | 0 | 0 | 0 | 0 | 0 | 0 | 0 | 1 | 0 |
| SLSLIQT | 2 | 0 | 0 | 112 | 46 | 136 | 0 | 2 | 0 | 1 | 1 | 0 | 0 | 0 | 0 | 0 | 0 | 0 | 1 | 0 |
| HALGPSS | 7 | 27 | 3 | 81 | 60 | 127 | 0 | 2 | 0 | 18 | 8 | 0 | 0 | 0 | 10 | 0 | 0 | 0 | 1 | 0 |
| TPPTMDH | 28 | 46 | 40 | 39 | 16 | 125 | 1 | 1 | 0 | 4 | 3 | 0 | 0 | 67 | 0 | 0 | 0 | 0 | 1 | 0 |
| MGLQTPY | 0 | 0 | 0 | 0 | 0 | 0 | 1133 | 2162 | 564 | 0 | 1 | 0 | 0 | 0 | 0 | 0 | 0 | 0 | 0 | 0 |
| SILPYPY | 36 | 92 | 409 | 16 | 10 | 12 | 857 | 1488 | 2536 | 13 | 8 | 1 | 10 | 847 | 22 | 0 | 0 | 0 | 1 | 0 |
| STASYTR | 645 | 3722 | 4203 | 1 | 4 | 0 | 807 | 3490 | 1957 | 17 | 6 | 1 | 38 | 1695 | 53 | 0 | 0 | 0 | 1 | 0 |
| NQLPLHA | 24 | 54 | 34 | 2 | 3 | 0 | 806 | 1085 | 879 | 10 | 4 | 1 | 0 | 312 | 12 | 0 | 0 | 0 | 1 | 0 |
| HSTKVAF | 0 | 0 | 0 | 0 | 0 | 0 | 772 | 1534 | 256 | 0 | 0 | 0 | 0 | 0 | 0 | 0 | 0 | 0 | 0 | 0 |
| MDAHHAL | 2 | 0 | 0 | 235 | 97 | 8 | 702 | 870 | 4 | 10 | 3 | 0 | 0 | 0 | 0 | 0 | 0 | 0 | 1 | 0 |
| QPWPTSI | 6 | 24 | 10 | 16 | 20 | 0 | 556 | 888 | 2104 | 1 | 1 | 0 | 0 | 0 | 19 | 0 | 0 | 1 | 1 | 0 |
| IPAPLRS | 0 | 0 | 0 | 0 | 0 | 0 | 496 | 1219 | 535 | 0 | 0 | 0 | 0 | 983 | 0 | 0 | 0 | 0 | 0 | 0 |
| HAIYPRH | 35 | 137 | 328 | 2659 | 3032 | 11887 | 481 | 661 | 370 | 99 | 28 | 2 | 148 | 8381 | 19 | 0 | 0 | 10 | 1 | 0 |
| QAHTVGK | 0 | 0 | 0 | 0 | 0 | 0 | 452 | 783 | 35 | 0 | 0 | 0 | 0 | 84 | 0 | 0 | 0 | 0 | 0 | 0 |
| TGHSAQG | 0 | 0 | 0 | 0 | 0 | 0 | 336 | 790 | 2946 | 1 | 0 | 0 | 25 | 78 | 0 | 0 | 0 | 0 | 0 | 0 |
| MPTLTPT | 0 | 0 | 0 | 0 | 0 | 0 | 330 | 147 | 251 | 2 | 0 | 0 | 0 | 0 | 0 | 0 | 0 | 0 | 1 | 0 |
| SLHQPHL | 0 | 0 | 0 | 0 | 0 | 0 | 316 | 651 | 9 | 6 | 0 | 0 | 0 | 0 | 0 | 0 | 0 | 0 | 1 | 0 |
| LPRTPTD | 0 | 0 | 0 | 0 | 0 | 0 | 260 | 355 | 44 | 2 | 1 | 0 | 3 | 0 | 0 | 0 | 0 | 0 | 1 | 0 |
| GKPMPPM | 63 | 208 | 165 | 1540 | 899 | 3479 | 254 | 375 | 287 | 49 | 11 | 0 | 3 | 24 | 0 | 0 | 0 | 0 | 1 | 0 |
| SPTGWAP | 0 | 0 | 0 | 1 | 0 | 0 | 226 | 417 | 1088 | 1 | 0 | 1 | 2 | 29 | 7 | 0 | 0 | 0 | 1 | 0 |
| TLLPFQP | 0 | 0 | 0 | 0 | 0 | 0 | 216 | 124 | 55 | 0 | 0 | 0 | 8 | 195 | 0 | 0 | 0 | 0 | 0 | 0 |
| FPSTITP | 17 | 8 | 21 | 2 | 5 | 105 | 179 | 280 | 1309 | 2 | 1 | 0 | 0 | 265 | 12 | 0 | 0 | 0 | 1 | 0 |
| AGNGTTP | 0 | 0 | 0 | 2 | 2 | 0 | 153 | 186 | 133 | 0 | 1 | 0 | 0 | 0 | 0 | 0 | 0 | 0 | 1 | 0 |
| QLHMDYR | 6 | 26 | 29 | 0 | 2 | 0 | 145 | 78 | 1 | 3 | 7 | 0 | 0 | 31 | 0 | 0 | 0 | 0 | 1 | 0 |
| STASYTR | 645 | 3722 | 4203 | 1 | 4 | 0 | 807 | 3490 | 1957 | 17 | 6 | 1 | 38 | 1695 | 53 | 0 | 0 | 0 | 1 | 0 |
| MGLQTPY | 0 | 0 | 0 | 0 | 0 | 0 | 1133 | 2162 | 564 | 0 | 1 | 0 | 0 | 0 | 0 | 0 | 0 | 0 | 0 | 0 |
| HSTKVAF | 0 | 0 | 0 | 0 | 0 | 0 | 772 | 1534 | 256 | 0 | 0 | 0 | 0 | 0 | 0 | 0 | 0 | 0 | 0 | 0 |
| SILPYPY | 36 | 92 | 409 | 16 | 10 | 12 | 857 | 1488 | 2536 | 13 | 8 | 1 | 10 | 847 | 22 | 0 | 0 | 0 | 1 | 0 |
| IPAPLRS | 0 | 0 | 0 | 0 | 0 | 0 | 496 | 1219 | 535 | 0 | 0 | 0 | 0 | 983 | 0 | 0 | 0 | 0 | 0 | 0 |
| NQLPLHA | 24 | 54 | 34 | 2 | 3 | 0 | 806 | 1085 | 879 | 10 | 4 | 1 | 0 | 312 | 12 | 0 | 0 | 0 | 1 | 0 |
| QPWPTSI | 6 | 24 | 10 | 16 | 20 | 0 | 556 | 888 | 2104 | 1 | 1 | 0 | 0 | 0 | 19 | 0 | 0 | 1 | 1 | 0 |
| MDAHHAL | 2 | 0 | 0 | 235 | 97 | 8 | 702 | 870 | 4 | 10 | 3 | 0 | 0 | 0 | 0 | 0 | 0 | 0 | 1 | 0 |
| TGHSAQG | 0 | 0 | 0 | 0 | 0 | 0 | 336 | 790 | 2946 | 1 | 0 | 0 | 25 | 78 | 0 | 0 | 0 | 0 | 0 | 0 |
| QAHTVGK | 0 | 0 | 0 | 0 | 0 | 0 | 452 | 783 | 35 | 0 | 0 | 0 | 0 | 84 | 0 | 0 | 0 | 0 | 0 | 0 |
| HAIYPRH | 35 | 137 | 328 | 2659 | 3032 | 11887 | 481 | 661 | 370 | 99 | 28 | 2 | 148 | 8381 | 19 | 0 | 0 | 10 | 1 | 0 |
| SLHQPHL | 0 | 0 | 0 | 0 | 0 | 0 | 316 | 651 | 9 | 6 | 0 | 0 | 0 | 0 | 0 | 0 | 0 | 0 | 1 | 0 |
| STTKLAL | 0 | 0 | 0 | 0 | 1 | 0 | 128 | 456 | 22 | 1 | 0 | 0 | 0 | 0 | 0 | 0 | 0 | 0 | 1 | 0 |
| SPTGWAP | 0 | 0 | 0 | 1 | 0 | 0 | 226 | 417 | 1088 | 1 | 0 | 1 | 2 | 29 | 7 | 0 | 0 | 0 | 1 | 0 |
| GKPMPPM | 63 | 208 | 165 | 1540 | 899 | 3479 | 254 | 375 | 287 | 49 | 11 | 0 | 3 | 24 | 0 | 0 | 0 | 0 | 1 | 0 |
| LPRTPTD | 0 | 0 | 0 | 0 | 0 | 0 | 260 | 355 | 44 | 2 | 1 | 0 | 3 | 0 | 0 | 0 | 0 | 0 | 1 | 0 |
| APRTFNQ | 0 | 0 | 0 | 0 | 0 | 0 | 67 | 343 | 3528 | 0 | 0 | 0 | 6 | 98 | 0 | 0 | 0 | 0 | 1 | 0 |
| STFTKSP | 1 | 0 | 4 | 27 | 20 | 141 | 83 | 335 | 626 | 7 | 3 | 0 | 0 | 0 | 0 | 0 | 0 | 0 | 1 | 0 |
| SYHSFNL | 0 | 0 | 0 | 0 | 0 | 0 | 120 | 334 | 9361 | 0 | 3 | 0 | 0 | 339 | 0 | 0 | 0 | 0 | 1 | 0 |
| SHSLLHH | 0 | 0 | 0 | 0 | 1 | 0 | 114 | 333 | 691 | 7 | 0 | 0 | 0 | 196 | 4 | 0 | 0 | 0 | 1 | 0 |

**Table S2 (cont.)**

| | Lot1-BA-Rep1-Round1 | Lot1-BA-Rep1-Round2 | Lot1-BA-Rep1-Round3 | Lot1-BA-Rep2-Round1 | Lot1-BA-Rep2-Round2 | Lot1-BA-Rep2-Round3 | Lot1-BA-Rep3-Round1 | Lot1-BA-Rep3-Round2 | Lot1-BA-Rep3-Round3 | Lot1-EmA-Rep1-Round1 | Lot1-EmA-Rep1-Round2 | Lot1-EmA-Rep1-Round3 | Lot2-EmA-Rep1-Round1 | Lot2-EmA-Rep2-Round1 | Lot2-EmA-Rep3-Round1 | Lot2-BA-Rep1-Round1 | Lot2-BA-Rep2-Round1 | Lot2-BA-Rep3-Round1 | Lot1-Parasites | Lot2-Parasites |
|---|---|---|---|---|---|---|---|---|---|---|---|---|---|---|---|---|---|---|---|---|
| SYHSFNL | 0 | 0 | 0 | 0 | 0 | 0 | 120 | 334 | 9361 | 0 | 3 | 0 | 0 | 339 | 0 | 0 | 0 | 0 | 1 | 0 |
| APRTFNQ | 0 | 0 | 0 | 0 | 0 | 0 | 67 | 343 | 3528 | 0 | 0 | 0 | 6 | 98 | 0 | 0 | 0 | 0 | 1 | 0 |
| TGHSAQG | 0 | 0 | 0 | 0 | 0 | 0 | 336 | 790 | 2946 | 1 | 0 | 0 | 25 | 78 | 0 | 0 | 0 | 0 | 0 | 0 |
| SILPYPY | 36 | 92 | 409 | 16 | 10 | 12 | 857 | 1488 | 2536 | 13 | 8 | 1 | 10 | 847 | 22 | 0 | 0 | 0 | 1 | 0 |
| QPWPTSI | 6 | 24 | 10 | 16 | 20 | 0 | 556 | 888 | 2104 | 1 | 1 | 0 | 0 | 0 | 19 | 0 | 0 | 1 | 1 | 0 |
| STASYTR | 645 | 3722 | 4203 | 1 | 4 | 0 | 807 | 3490 | 1957 | 17 | 6 | 1 | 38 | 1695 | 53 | 0 | 0 | 0 | 1 | 0 |
| HTIQFTP | 0 | 0 | 0 | 10 | 14 | 17 | 18 | 15 | 1564 | 4 | 0 | 0 | 5 | 18 | 0 | 0 | 0 | 0 | 1 | 0 |
| FPSTITP | 17 | 8 | 21 | 2 | 5 | 105 | 179 | 280 | 1309 | 2 | 1 | 0 | 0 | 265 | 12 | 0 | 0 | 0 | 1 | 0 |
| ASYSGTA | 15 | 18 | 23 | 33 | 26 | 9 | 19 | 55 | 1121 | 8 | 3 | 1 | 3 | 227 | 0 | 0 | 0 | 0 | 1 | 0 |
| SPTGWAP | 0 | 0 | 0 | 1 | 0 | 0 | 226 | 417 | 1088 | 1 | 0 | 1 | 2 | 29 | 7 | 0 | 0 | 0 | 1 | 0 |
| NQLPLHA | 24 | 54 | 34 | 2 | 3 | 0 | 806 | 1085 | 879 | 10 | 4 | 1 | 0 | 312 | 12 | 0 | 0 | 0 | 1 | 0 |
| SHSLLHH | 0 | 0 | 0 | 0 | 1 | 0 | 114 | 333 | 691 | 7 | 0 | 0 | 0 | 196 | 4 | 0 | 0 | 0 | 1 | 0 |
| STFTKSP | 1 | 0 | 4 | 27 | 20 | 141 | 83 | 335 | 626 | 7 | 3 | 0 | 0 | 0 | 0 | 0 | 0 | 0 | 1 | 0 |
| MGLQTPY | 0 | 0 | 0 | 0 | 0 | 0 | 1133 | 2162 | 564 | 0 | 1 | 0 | 0 | 0 | 0 | 0 | 0 | 0 | 0 | 0 |
| IPAPLRS | 0 | 0 | 0 | 0 | 0 | 0 | 496 | 1219 | 535 | 0 | 0 | 0 | 0 | 983 | 0 | 0 | 0 | 0 | 0 | 0 |
| HAIYPRH | 35 | 137 | 328 | 2659 | 3032 | 11887 | 481 | 661 | 370 | 99 | 28 | 2 | 148 | 8381 | 19 | 0 | 0 | 10 | 1 | 0 |
| STPIQQP | 5 | 6 | 2 | 0 | 0 | 0 | 25 | 37 | 354 | 8 | 4 | 0 | 0 | 246 | 0 | 0 | 0 | 0 | 1 | 0 |
| SHHQKPP | 0 | 0 | 0 | 0 | 0 | 0 | 6 | 10 | 294 | 1 | 0 | 0 | 0 | 0 | 0 | 0 | 0 | 0 | 0 | 0 |
| GKPMPPM | 63 | 208 | 165 | 1540 | 899 | 3479 | 254 | 375 | 287 | 49 | 11 | 0 | 3 | 24 | 0 | 0 | 0 | 0 | 1 | 0 |
| HSTKVAF | 0 | 0 | 0 | 0 | 0 | 0 | 772 | 1534 | 256 | 0 | 0 | 0 | 0 | 0 | 0 | 0 | 0 | 0 | 0 | 0 |
| HAIYPRH | 35 | 137 | 328 | 2659 | 3032 | 11887 | 481 | 661 | 370 | 99 | 28 | 2 | 148 | 8381 | 19 | 0 | 0 | 10 | 1 | 0 |
| GKPMPPM | 63 | 208 | 165 | 1540 | 899 | 3479 | 254 | 375 | 287 | 49 | 11 | 0 | 3 | 24 | 0 | 0 | 0 | 0 | 1 | 0 |
| GPMLARG | 112 | 166 | 101 | 0 | 0 | 0 | 0 | 0 | 0 | 34 | 10 | 0 | 0 | 43 | 0 | 0 | 0 | 0 | 1 | 0 |
| AMSSRSL | 0 | 0 | 0 | 40 | 61 | 3 | 8 | 32 | 117 | 28 | 10 | 0 | 14 | 245 | 0 | 0 | 0 | 0 | 1 | 0 |
| SSALLLP | 0 | 0 | 0 | 0 | 0 | 0 | 0 | 0 | 0 | 24 | 10 | 2 | 0 | 0 | 0 | 0 | 0 | 0 | 0 | 0 |
| DSHTPQR | 141 | 349 | 208 | 12 | 9 | 0 | 11 | 24 | 3 | 21 | 8 | 0 | 0 | 4 | 0 | 0 | 0 | 0 | 1 | 0 |
| QPPRSTS | 23 | 33 | 11 | 1775 | 938 | 10032 | 8 | 4 | 0 | 19 | 5 | 0 | 0 | 847 | 13 | 0 | 0 | 0 | 1 | 0 |
| AASSLTI | 0 | 0 | 0 | 0 | 0 | 0 | 0 | 0 | 0 | 19 | 8 | 0 | 0 | 0 | 0 | 0 | 0 | 0 | 0 | 0 |
| HALGPSS | 7 | 27 | 3 | 81 | 60 | 127 | 0 | 2 | 0 | 18 | 8 | 0 | 0 | 0 | 10 | 0 | 0 | 0 | 1 | 0 |
| STASYTR | 645 | 3722 | 4203 | 1 | 4 | 0 | 807 | 3490 | 1957 | 17 | 6 | 1 | 38 | 1695 | 53 | 0 | 0 | 0 | 1 | 0 |
| EPLQLKM | 214 | 777 | 586 | 3 | 2 | 0 | 2 | 0 | 0 | 16 | 6 | 0 | 7 | 1159 | 6 | 1 | 0 | 0 | 1 | 0 |
| QATHRSH | 128 | 229 | 147 | 23 | 8 | 1 | 6 | 3 | 0 | 14 | 8 | 1 | 7 | 556 | 0 | 0 | 0 | 0 | 1 | 0 |
| GKVQAQS | 57 | 51 | 39 | 0 | 1 | 0 | 28 | 41 | 0 | 14 | 7 | 0 | 0 | 439 | 0 | 0 | 0 | 0 | 1 | 0 |
| SSSVVTH | 111 | 92 | 0 | 0 | 0 | 0 | 0 | 0 | 0 | 14 | 0 | 0 | 0 | 0 | 0 | 0 | 0 | 0 | 0 | 0 |
| SILPYPY | 36 | 92 | 409 | 16 | 10 | 12 | 857 | 1488 | 2536 | 13 | 8 | 1 | 10 | 847 | 22 | 0 | 0 | 0 | 1 | 0 |
| SPTQPKS | 34 | 61 | 31 | 88 | 64 | 1120 | 9 | 7 | 0 | 13 | 2 | 0 | 0 | 0 | 0 | 0 | 0 | 0 | 1 | 0 |
| ANTTPRH | 236 | 523 | 289 | 0 | 0 | 0 | 2 | 8 | 1 | 12 | 0 | 3 | 0 | 475 | 32 | 0 | 0 | 0 | 1 | 0 |
| QTGYATR | 0 | 0 | 0 | 0 | 0 | 0 | 0 | 0 | 0 | 12 | 10 | 3 | 0 | 0 | 0 | 0 | 0 | 0 | 0 | 0 |
| TYQNPVH | 0 | 0 | 0 | 0 | 0 | 0 | 0 | 0 | 0 | 12 | 6 | 2 | 0 | 0 | 0 | 0 | 0 | 0 | 0 | 0 |
| NQLAGSG | 0 | 0 | 0 | 1 | 1 | 0 | 1 | 1 | 1 | 11 | 1984 | 21310 | 4055 | 83 | 19 | 0 | 0 | 0 | 0 | 0 |
| SWQYGKL | 0 | 0 | 0 | 1 | 0 | 0 | 0 | 2 | 0 | 9 | 3143 | 22060 | 5814 | 848 | 42 | 0 | 1 | 0 | 0 | 0 |
| NQLAGSG | 0 | 0 | 0 | 1 | 1 | 0 | 1 | 1 | 1 | 11 | 1984 | 21310 | 4055 | 83 | 19 | 0 | 0 | 0 | 0 | 0 |
| HWHFGPL | 0 | 0 | 0 | 0 | 0 | 0 | 1 | 1 | 0 | 3 | 228 | 1929 | 510 | 0 | 26 | 0 | 0 | 0 | 0 | 0 |
| TYRFGPL | 0 | 0 | 0 | 0 | 0 | 0 | 0 | 0 | 0 | 0 | 136 | 923 | 347 | 12 | 0 | 0 | 0 | 0 | 0 | 0 |
| SWKFGPL | 0 | 0 | 0 | 0 | 0 | 0 | 0 | 2 | 0 | 1 | 109 | 899 | 296 | 15 | 0 | 0 | 0 | 0 | 0 | 0 |
| ALEVTFW | 0 | 0 | 0 | 0 | 0 | 0 | 0 | 0 | 0 | 3 | 80 | 25 | 9 | 0 | 0 | 0 | 0 | 0 | 0 | 0 |
| TYKFGTL | 0 | 0 | 0 | 0 | 0 | 0 | 0 | 0 | 0 | 2 | 78 | 1107 | 545 | 0 | 7 | 0 | 0 | 0 | 0 | 0 |
| VQNEWRS | 0 | 0 | 0 | 0 | 0 | 0 | 0 | 0 | 0 | 4 | 70 | 21 | 13 | 0 | 0 | 0 | 0 | 0 | 0 | 0 |
| LTVEPWL | 0 | 0 | 0 | 0 | 0 | 0 | 0 | 0 | 0 | 2 | 63 | 118 | 108 | 0 | 0 | 1 | 0 | 0 | 0 | 0 |
| ELWVSPL | 0 | 0 | 0 | 0 | 0 | 0 | 0 | 0 | 0 | 2 | 54 | 28 | 21 | 0 | 0 | 0 | 0 | 0 | 0 | 0 |
| LEVYALV | 0 | 0 | 0 | 0 | 0 | 0 | 0 | 0 | 0 | 2 | 45 | 16 | 24 | 0 | 0 | 0 | 0 | 0 | 0 | 0 |
| TYKYYPL | 0 | 0 | 0 | 0 | 0 | 0 | 0 | 0 | 0 | 1 | 43 | 177 | 169 | 1 | 0 | 0 | 0 | 0 | 0 | 0 |
| TFKFGPL | 0 | 0 | 0 | 0 | 0 | 0 | 0 | 0 | 0 | 0 | 41 | 257 | 123 | 0 | 0 | 0 | 0 | 0 | 0 | 0 |
| TYQYGKL | 0 | 0 | 0 | 0 | 0 | 0 | 0 | 0 | 0 | 1 | 40 | 122 | 107 | 0 | 12 | 0 | 0 | 0 | 0 | 0 |
| TYLFQPL | 0 | 0 | 0 | 0 | 0 | 0 | 0 | 0 | 0 | 1 | 35 | 172 | 146 | 12 | 0 | 0 | 0 | 0 | 0 | 0 |
| DLTVTPW | 0 | 0 | 0 | 0 | 0 | 0 | 0 | 0 | 0 | 0 | 35 | 41 | 36 | 0 | 0 | 0 | 0 | 0 | 0 | 0 |
| TWKFSPL | 0 | 0 | 0 | 0 | 0 | 0 | 0 | 0 | 0 | 1 | 34 | 187 | 183 | 0 | 0 | 0 | 0 | 0 | 0 | 0 |
| QLTVMSW | 0 | 0 | 0 | 0 | 0 | 0 | 0 | 0 | 0 | 0 | 34 | 21 | 13 | 0 | 0 | 0 | 0 | 0 | 0 | 0 |
| LTVQPWP | 0 | 0 | 0 | 0 | 0 | 0 | 0 | 1 | 0 | 2 | 32 | 17 | 17 | 0 | 0 | 0 | 0 | 0 | 0 | 0 |
| HAIYPRH | 35 | 137 | 328 | 2659 | 3032 | 11887 | 481 | 661 | 370 | 99 | 28 | 2 | 148 | 8381 | 19 | 0 | 0 | 10 | 1 | 0 |
| SWQYGKL | 0 | 0 | 0 | 1 | 0 | 0 | 0 | 2 | 0 | 9 | 3143 | 22060 | 5814 | 848 | 42 | 0 | 1 | 0 | 0 | 0 |
| NQLAGSG | 0 | 0 | 0 | 1 | 1 | 0 | 1 | 1 | 1 | 11 | 1984 | 21310 | 4055 | 83 | 19 | 0 | 0 | 0 | 0 | 0 |
| HWHFGPL | 0 | 0 | 0 | 0 | 0 | 0 | 1 | 1 | 0 | 3 | 228 | 1929 | 510 | 0 | 26 | 0 | 0 | 0 | 0 | 0 |
| TYKFGTL | 0 | 0 | 0 | 0 | 0 | 0 | 0 | 0 | 0 | 2 | 78 | 1107 | 545 | 0 | 7 | 0 | 0 | 0 | 0 | 0 |
| TYRFGPL | 0 | 0 | 0 | 0 | 0 | 0 | 0 | 0 | 0 | 0 | 136 | 923 | 347 | 12 | 0 | 0 | 0 | 0 | 0 | 0 |
| SWKFGPL | 0 | 0 | 0 | 0 | 0 | 0 | 0 | 2 | 0 | 1 | 109 | 899 | 296 | 15 | 0 | 0 | 0 | 0 | 0 | 0 |
| HWKFGIL | 0 | 0 | 0 | 0 | 0 | 0 | 0 | 0 | 0 | 0 | 3 | 556 | 51 | 0 | 3 | 0 | 0 | 0 | 0 | 0 |
| TFKFGPL | 0 | 0 | 0 | 0 | 0 | 0 | 0 | 0 | 0 | 0 | 41 | 257 | 123 | 0 | 0 | 0 | 0 | 0 | 0 | 0 |
| TWKFSPL | 0 | 0 | 0 | 0 | 0 | 0 | 0 | 0 | 0 | 1 | 34 | 187 | 183 | 0 | 0 | 0 | 0 | 0 | 0 | 0 |
| TYKYYPL | 0 | 0 | 0 | 0 | 0 | 0 | 0 | 0 | 0 | 1 | 43 | 177 | 169 | 1 | 0 | 0 | 0 | 0 | 0 | 0 |
| TYLFQPL | 0 | 0 | 0 | 0 | 0 | 0 | 0 | 0 | 0 | 1 | 35 | 172 | 146 | 12 | 0 | 0 | 0 | 0 | 0 | 0 |
| TYQYGKL | 0 | 0 | 0 | 0 | 0 | 0 | 0 | 0 | 0 | 1 | 40 | 122 | 107 | 0 | 12 | 0 | 0 | 0 | 0 | 0 |
| LTVEPWL | 0 | 0 | 0 | 0 | 0 | 0 | 0 | 0 | 0 | 2 | 63 | 118 | 108 | 0 | 0 | 1 | 0 | 0 | 0 | 0 |
| HWKYWPL | 0 | 0 | 0 | 0 | 0 | 0 | 0 | 0 | 0 | 1 | 12 | 81 | 35 | 0 | 0 | 0 | 0 | 0 | 0 | 0 |
| TYVFYPL | 0 | 0 | 0 | 0 | 0 | 0 | 0 | 0 | 0 | 0 | 18 | 44 | 40 | 0 | 0 | 0 | 0 | 0 | 0 | 0 |
| DLTVTPW | 0 | 0 | 0 | 0 | 0 | 0 | 0 | 0 | 0 | 0 | 35 | 41 | 36 | 0 | 0 | 0 | 0 | 0 | 0 | 0 |
| LEVFPYY | 0 | 0 | 0 | 0 | 0 | 0 | 0 | 0 | 0 | 1 | 24 | 35 | 15 | 0 | 0 | 0 | 0 | 0 | 0 | 0 |
| ELWVSPL | 0 | 0 | 0 | 0 | 0 | 0 | 0 | 0 | 0 | 2 | 54 | 28 | 21 | 0 | 0 | 0 | 0 | 0 | 0 | 0 |
| KVWELHP | 0 | 0 | 0 | 0 | 0 | 0 | 0 | 0 | 0 | 0 | 6 | 27 | 17 | 0 | 0 | 0 | 0 | 0 | 0 | 0 |
| TYRFLPL | 0 | 0 | 0 | 0 | 0 | 0 | 0 | 0 | 0 | 1 | 2 | 26 | 11 | 0 | 0 | 0 | 0 | 0 | 0 | 0 |

**Table S2 (cont.)**

| | Lot1-BA-Rep1-Round1 | Lot1-BA-Rep1-Round2 | Lot1-BA-Rep1-Round3 | Lot1-BA-Rep2-Round1 | Lot1-BA-Rep2-Round2 | Lot1-BA-Rep2-Round3 | Lot1-BA-Rep3-Round1 | Lot1-BA-Rep3-Round2 | Lot1-BA-Rep3-Round3 | Lot1-EmA-Rep1-Round1 | Lot1-EmA-Rep1-Round2 | Lot1-EmA-Rep1-Round3 | Lot2-EmA-Rep1-Round1 | Lot2-EmA-Rep2-Round1 | Lot2-EmA-Rep3-Round1 | Lot2-BA-Rep1-Round1 | Lot2-BA-Rep2-Round1 | Lot2-BA-Rep3-Round1 | Lot1-Parasites | Lot2-Parasites |
|---|---|---|---|---|---|---|---|---|---|---|---|---|---|---|---|---|---|---|---|---|
| SWQYGKL | 0 | 0 | 0 | 1 | 0 | 0 | 0 | 2 | 0 | 9 | 3143 | 22060 | 5814 | 848 | 42 | 0 | 1 | 0 | 0 | 0 |
| NQLAGSG | 0 | 0 | 0 | 1 | 1 | 0 | 1 | 1 | 1 | 11 | 1984 | 21310 | 4055 | 83 | 19 | 0 | 0 | 0 | 0 | 0 |
| TSSESES | 0 | 0 | 0 | 0 | 0 | 0 | 0 | 0 | 0 | 0 | 0 | 0 | 665 | 0 | 0 | 0 | 0 | 0 | 0 | 0 |
| ERTVLHT | 0 | 0 | 0 | 0 | 0 | 0 | 0 | 0 | 0 | 0 | 0 | 0 | 554 | 0 | 0 | 0 | 0 | 0 | 0 | 0 |
| TYKFGTL | 0 | 0 | 0 | 0 | 0 | 0 | 0 | 0 | 0 | 2 | 78 | 1107 | 545 | 0 | 7 | 0 | 0 | 0 | 0 | 0 |
| HWHFGPL | 0 | 0 | 0 | 0 | 0 | 0 | 1 | 1 | 0 | 3 | 228 | 1929 | 510 | 0 | 26 | 0 | 0 | 0 | 0 | 0 |
| TYRFGPL | 0 | 0 | 0 | 0 | 0 | 0 | 0 | 0 | 0 | 0 | 136 | 923 | 347 | 12 | 0 | 0 | 0 | 0 | 0 | 0 |
| RFTVDWD | 0 | 0 | 0 | 0 | 0 | 0 | 0 | 0 | 0 | 0 | 0 | 0 | 309 | 0 | 0 | 0 | 0 | 0 | 0 | 0 |
| SWKFGPL | 0 | 0 | 0 | 0 | 0 | 0 | 0 | 2 | 0 | 1 | 109 | 899 | 296 | 15 | 0 | 0 | 0 | 0 | 0 | 0 |
| LAGPLMT | 0 | 0 | 0 | 0 | 0 | 0 | 0 | 0 | 0 | 0 | 0 | 0 | 192 | 0 | 0 | 0 | 0 | 0 | 0 | 0 |
| TWKFSPL | 0 | 0 | 0 | 0 | 0 | 0 | 0 | 0 | 0 | 1 | 34 | 187 | 183 | 0 | 0 | 0 | 0 | 0 | 0 | 0 |
| NVSGSHS | 0 | 0 | 0 | 0 | 0 | 0 | 0 | 0 | 0 | 0 | 0 | 0 | 180 | 0 | 0 | 0 | 0 | 0 | 0 | 0 |
| SVLLPHR | 0 | 0 | 0 | 0 | 0 | 0 | 0 | 0 | 0 | 0 | 0 | 0 | 177 | 0 | 0 | 0 | 0 | 0 | 0 | 0 |
| TYKYYPL | 0 | 0 | 0 | 0 | 0 | 0 | 0 | 0 | 0 | 1 | 43 | 177 | 169 | 1 | 0 | 0 | 0 | 0 | 0 | 0 |
| HHQYVPA | 0 | 0 | 0 | 0 | 0 | 0 | 0 | 0 | 0 | 0 | 0 | 0 | 161 | 0 | 0 | 0 | 0 | 0 | 0 | 0 |
| WTTTSRL | 0 | 0 | 0 | 0 | 0 | 0 | 0 | 0 | 0 | 0 | 0 | 0 | 161 | 0 | 0 | 0 | 0 | 0 | 0 | 0 |
| QLMPMMM | 0 | 0 | 0 | 0 | 0 | 0 | 0 | 0 | 0 | 0 | 0 | 0 | 159 | 0 | 0 | 0 | 0 | 0 | 0 | 0 |
| RPYDTAH | 0 | 0 | 0 | 0 | 0 | 0 | 0 | 0 | 0 | 0 | 0 | 0 | 157 | 0 | 0 | 0 | 0 | 0 | 0 | 0 |
| SAGSMHL | 0 | 0 | 0 | 0 | 0 | 0 | 0 | 0 | 0 | 0 | 0 | 0 | 154 | 0 | 0 | 0 | 0 | 0 | 0 | 0 |
| GLYHSAT | 0 | 0 | 0 | 0 | 0 | 0 | 0 | 0 | 0 | 0 | 0 | 0 | 153 | 0 | 0 | 0 | 0 | 0 | 0 | 0 |
| HAIYPRH | 35 | 137 | 328 | 2659 | 3032 | 11887 | 481 | 661 | 370 | 99 | 28 | 2 | 148 | 8381 | 19 | 0 | 0 | 10 | 1 | 0 |
| YLTMPTP | 191 | 749 | 1254 | 4 | 4 | 0 | 71 | 51 | 0 | 7 | 4 | 1 | 18 | 2296 | 15 | 0 | 0 | 0 | 1 | 0 |
| STASYTR | 645 | 3722 | 4203 | 1 | 4 | 0 | 807 | 3490 | 1957 | 17 | 6 | 1 | 38 | 1695 | 53 | 0 | 0 | 0 | 1 | 0 |
| TPQSSPT | 400 | 612 | 207 | 0 | 0 | 0 | 0 | 0 | 0 | 3 | 8 | 1 | 22 | 1408 | 50 | 0 | 0 | 0 | 1 | 0 |
| EPLQLKM | 214 | 777 | 586 | 3 | 2 | 0 | 2 | 0 | 0 | 16 | 6 | 0 | 7 | 1159 | 6 | 1 | 0 | 0 | 1 | 0 |
| GVKALST | 236 | 497 | 295 | 42 | 7 | 22 | 0 | 0 | 0 | 6 | 1 | 0 | 5 | 1139 | 10 | 0 | 0 | 0 | 1 | 0 |
| TPFMAYH | 10 | 15 | 13 | 0 | 0 | 0 | 0 | 0 | 0 | 3 | 0 | 0 | 0 | 1083 | 0 | 0 | 0 | 0 | 1 | 0 |
| IPAPLRS | 0 | 0 | 0 | 0 | 0 | 0 | 496 | 1219 | 535 | 0 | 0 | 0 | 0 | 983 | 0 | 0 | 0 | 0 | 0 | 0 |
| RLPSWHE | 34 | 25 | 6 | 0 | 0 | 0 | 6 | 1 | 8 | 4 | 0 | 0 | 0 | 912 | 0 | 0 | 0 | 0 | 1 | 0 |
| AYPEPYV | 0 | 0 | 0 | 0 | 0 | 0 | 0 | 0 | 0 | 1 | 0 | 0 | 0 | 910 | 0 | 0 | 0 | 0 | 0 | 0 |
| QPTHPTR | 0 | 0 | 0 | 827 | 524 | 3585 | 3 | 0 | 0 | 0 | 0 | 0 | 0 | 889 | 0 | 0 | 0 | 0 | 0 | 0 |
| SWQYGKL | 0 | 0 | 0 | 1 | 0 | 0 | 0 | 2 | 0 | 9 | 3143 | 22060 | 5814 | 848 | 42 | 0 | 1 | 0 | 0 | 0 |
| QPPRSTS | 23 | 33 | 11 | 1775 | 938 | 10032 | 8 | 4 | 0 | 19 | 5 | 0 | 0 | 847 | 13 | 0 | 0 | 0 | 1 | 0 |
| SILPYPY | 36 | 92 | 409 | 16 | 10 | 12 | 857 | 1488 | 2536 | 13 | 8 | 1 | 10 | 847 | 22 | 0 | 0 | 0 | 1 | 0 |
| TQTMRST | 92 | 126 | 48 | 0 | 0 | 0 | 0 | 0 | 0 | 0 | 0 | 0 | 11 | 749 | 0 | 0 | 0 | 0 | 0 | 0 |
| TKTDTWL | 95 | 216 | 219 | 2 | 1 | 0 | 3 | 2 | 0 | 11 | 5 | 1 | 21 | 725 | 34 | 0 | 0 | 0 | 1 | 0 |
| TPMTRAL | 4 | 0 | 0 | 0 | 0 | 0 | 0 | 0 | 0 | 0 | 0 | 0 | 0 | 711 | 0 | 0 | 0 | 0 | 0 | 0 |
| VMSQPHP | 0 | 0 | 0 | 0 | 0 | 0 | 0 | 0 | 0 | 0 | 0 | 0 | 0 | 708 | 0 | 0 | 0 | 0 | 0 | 0 |
| SPWDARL | 281 | 705 | 593 | 3 | 6 | 0 | 0 | 1 | 0 | 1 | 0 | 0 | 41 | 697 | 39 | 0 | 0 | 0 | 1 | 0 |
| GSTVFTA | 0 | 0 | 0 | 0 | 0 | 0 | 0 | 1 | 0 | 0 | 0 | 0 | 0 | 675 | 0 | 0 | 0 | 0 | 0 | 0 |
| EQGRPLP | 0 | 0 | 0 | 0 | 0 | 0 | 0 | 0 | 0 | 0 | 0 | 0 | 0 | 0 | 349 | 0 | 0 | 0 | 0 | 0 |
| MAANGAR | 0 | 0 | 0 | 0 | 0 | 0 | 0 | 0 | 0 | 0 | 0 | 0 | 0 | 0 | 262 | 0 | 0 | 0 | 0 | 0 |
| QVLLTAA | 0 | 0 | 0 | 0 | 0 | 0 | 0 | 0 | 0 | 0 | 0 | 0 | 0 | 0 | 243 | 0 | 0 | 0 | 0 | 0 |
| AGRELCC | 0 | 0 | 0 | 0 | 0 | 0 | 0 | 0 | 0 | 0 | 0 | 0 | 0 | 0 | 220 | 0 | 0 | 0 | 0 | 0 |
| ARAVLQL | 0 | 0 | 0 | 0 | 0 | 0 | 0 | 0 | 0 | 0 | 0 | 0 | 0 | 0 | 191 | 0 | 0 | 0 | 0 | 0 |
| QNMQQQI | 0 | 0 | 0 | 0 | 0 | 0 | 0 | 0 | 0 | 0 | 0 | 0 | 0 | 0 | 190 | 0 | 0 | 0 | 0 | 0 |
| AWSAVMR | 0 | 0 | 0 | 0 | 0 | 0 | 0 | 0 | 0 | 0 | 0 | 0 | 0 | 0 | 190 | 0 | 0 | 0 | 0 | 0 |
| APIWMHV | 0 | 0 | 0 | 0 | 0 | 0 | 0 | 0 | 0 | 0 | 0 | 0 | 0 | 0 | 186 | 0 | 0 | 0 | 0 | 0 |
| ATWQLGT | 0 | 0 | 0 | 0 | 0 | 0 | 0 | 0 | 0 | 0 | 0 | 0 | 0 | 0 | 181 | 0 | 0 | 0 | 0 | 0 |
| ASWIPLP | 0 | 0 | 0 | 0 | 0 | 0 | 0 | 0 | 0 | 0 | 0 | 0 | 0 | 0 | 172 | 0 | 0 | 0 | 0 | 0 |
| QQQYMAH | 0 | 0 | 0 | 0 | 0 | 0 | 0 | 0 | 0 | 0 | 0 | 0 | 0 | 0 | 164 | 0 | 0 | 0 | 0 | 0 |
| STAMDGR | 0 | 0 | 0 | 0 | 0 | 0 | 0 | 0 | 0 | 0 | 0 | 0 | 0 | 0 | 163 | 0 | 0 | 0 | 0 | 0 |
| GWRTTWP | 0 | 0 | 0 | 0 | 0 | 0 | 0 | 0 | 0 | 0 | 0 | 0 | 0 | 0 | 161 | 0 | 0 | 0 | 0 | 0 |
| WMASMAV | 0 | 0 | 0 | 0 | 0 | 0 | 0 | 0 | 0 | 0 | 0 | 0 | 0 | 0 | 141 | 0 | 0 | 0 | 0 | 0 |
| HEQPMHR | 0 | 0 | 0 | 0 | 0 | 0 | 0 | 0 | 0 | 0 | 0 | 0 | 0 | 0 | 132 | 0 | 0 | 0 | 0 | 0 |
| SLDVRMW | 0 | 0 | 0 | 0 | 0 | 0 | 0 | 0 | 0 | 0 | 0 | 0 | 4 | 19 | 126 | 931 | 25 | 0 | 0 | 1 |
| QHMPLTR | 0 | 0 | 0 | 0 | 0 | 0 | 0 | 0 | 0 | 0 | 0 | 0 | 0 | 0 | 124 | 0 | 0 | 0 | 0 | 0 |
| GEELPTL | 0 | 0 | 0 | 0 | 0 | 0 | 0 | 0 | 0 | 0 | 0 | 0 | 0 | 0 | 118 | 0 | 0 | 0 | 0 | 0 |
| HAMWFSV | 0 | 0 | 0 | 0 | 0 | 0 | 0 | 0 | 0 | 0 | 0 | 0 | 0 | 0 | 117 | 0 | 0 | 0 | 0 | 0 |
| APSGLSR | 0 | 0 | 0 | 0 | 0 | 0 | 0 | 0 | 0 | 0 | 0 | 0 | 0 | 0 | 114 | 0 | 0 | 0 | 0 | 0 |
| STPATLI | 0 | 0 | 0 | 0 | 0 | 0 | 0 | 0 | 0 | 0 | 0 | 0 | 0 | 0 | 0 | 3298 | 10 | 0 | 0 | 1 |
| WSLSELH | 0 | 0 | 0 | 0 | 0 | 0 | 0 | 0 | 0 | 0 | 0 | 0 | 3 | 8 | 21 | 3240 | 151 | 0 | 0 | 1 |
| LPVRLDW | 0 | 0 | 0 | 0 | 0 | 0 | 0 | 0 | 0 | 0 | 0 | 0 | 0 | 0 | 0 | 3042 | 0 | 0 | 0 | 1 |
| QTWLEMG | 0 | 0 | 0 | 0 | 0 | 0 | 0 | 0 | 0 | 0 | 0 | 0 | 0 | 0 | 0 | 2864 | 0 | 4 | 0 | 1 |
| GPHNPTQ | 0 | 0 | 0 | 0 | 0 | 0 | 0 | 0 | 0 | 0 | 0 | 0 | 0 | 0 | 0 | 2807 | 0 | 0 | 0 | 1 |
| NDRPHMP | 0 | 0 | 0 | 0 | 0 | 0 | 0 | 0 | 0 | 0 | 0 | 0 | 32 | 0 | 1 | 2565 | 0 | 0 | 0 | 1 |
| VPNIVTQ | 0 | 0 | 0 | 0 | 0 | 0 | 0 | 0 | 0 | 0 | 0 | 0 | 48 | 0 | 5 | 1972 | 38 | 0 | 0 | 1 |
| LRSDPVV | 0 | 0 | 0 | 0 | 0 | 0 | 0 | 0 | 0 | 0 | 0 | 0 | 0 | 0 | 1 | 1921 | 0 | 0 | 0 | 1 |
| VPASPWT | 0 | 0 | 0 | 0 | 0 | 0 | 0 | 0 | 0 | 0 | 0 | 0 | 0 | 0 | 31 | 1753 | 21 | 0 | 0 | 1 |
| SSVSWLN | 0 | 0 | 0 | 0 | 0 | 0 | 0 | 0 | 0 | 0 | 0 | 0 | 0 | 0 | 0 | 1586 | 0 | 0 | 0 | 1 |
| TTQVLEA | 0 | 0 | 0 | 0 | 0 | 0 | 0 | 0 | 0 | 0 | 0 | 0 | 10 | 0 | 103 | 1413 | 6 | 1693 | 0 | 1 |
| DAIPTSV | 0 | 0 | 0 | 0 | 0 | 0 | 0 | 0 | 0 | 0 | 0 | 0 | 10 | 40 | 0 | 1197 | 39 | 38 | 0 | 1 |
| VGKTSFQ | 0 | 0 | 0 | 0 | 0 | 0 | 0 | 0 | 0 | 0 | 0 | 0 | 0 | 0 | 0 | 1067 | 1 | 0 | 0 | 1 |
| SLDVRMW | 0 | 0 | 0 | 0 | 0 | 0 | 0 | 0 | 0 | 0 | 0 | 0 | 4 | 19 | 126 | 931 | 25 | 0 | 0 | 1 |
| GQVALLD | 0 | 0 | 0 | 0 | 0 | 0 | 0 | 0 | 0 | 0 | 0 | 0 | 0 | 0 | 0 | 850 | 0 | 0 | 0 | 0 |
| VENVHVR | 0 | 0 | 0 | 0 | 0 | 0 | 0 | 0 | 0 | 0 | 0 | 0 | 0 | 0 | 0 | 848 | 0 | 0 | 0 | 1 |
| VPVTMYW | 0 | 0 | 0 | 0 | 0 | 0 | 0 | 0 | 0 | 0 | 0 | 0 | 6 | 0 | 0 | 835 | 0 | 10 | 0 | 1 |
| ELGTTQT | 0 | 0 | 0 | 0 | 0 | 0 | 0 | 0 | 0 | 0 | 0 | 0 | 0 | 0 | 0 | 823 | 0 | 0 | 0 | 0 |
| AMTALDL | 0 | 0 | 0 | 0 | 0 | 0 | 0 | 0 | 0 | 0 | 0 | 0 | 0 | 7 | 0 | 745 | 0 | 5 | 0 | 1 |
| MAVTQKY | 0 | 0 | 0 | 0 | 0 | 0 | 0 | 0 | 0 | 0 | 0 | 0 | 0 | 0 | 0 | 707 | 0 | 0 | 0 | 1 |

| | Lot1-BA-Rep1-Round1 | Lot1-BA-Rep1-Round2 | Lot1-BA-Rep1-Round3 | Lot1-BA-Rep2-Round1 | Lot1-BA-Rep2-Round2 | Lot1-BA-Rep2-Round3 | Lot1-BA-Rep3-Round1 | Lot1-BA-Rep3-Round2 | Lot1-BA-Rep3-Round3 | Lot1-EmA-Rep1-Round1 | Lot1-EmA-Rep1-Round2 | Lot1-EmA-Rep1-Round3 | Lot2-EmA-Rep1-Round1 | Lot2-EmA-Rep2-Round1 | Lot2-EmA-Rep3-Round1 | Lot2-BA-Rep1-Round1 | Lot2-BA-Rep2-Round1 | Lot2-BA-Rep3-Round1 | Lot1-Parasites | Lot2-Parasites |
|---|---|---|---|---|---|---|---|---|---|---|---|---|---|---|---|---|---|---|---|---|
| AGSVIDT | 0 | 0 | 0 | 0 | 0 | 0 | 0 | 0 | 0 | 0 | 0 | 0 | 0 | 0 | 0 | 0 | 4722 | 396 | 0 | 1 |
| QAYHVSA | 0 | 0 | 0 | 0 | 0 | 0 | 0 | 0 | 0 | 0 | 0 | 0 | 0 | 0 | 0 | 0 | 4028 | 0 | 0 | 0 |
| SNMTRWH | 0 | 0 | 0 | 0 | 0 | 0 | 0 | 0 | 0 | 0 | 0 | 0 | 0 | 0 | 0 | 0 | 3593 | 0 | 0 | 1 |
| GRLDTGI | 0 | 0 | 0 | 0 | 0 | 0 | 0 | 0 | 0 | 0 | 0 | 0 | 0 | 0 | 1 | 1 | 3058 | 0 | 0 | 1 |
| NAYGGRI | 0 | 0 | 0 | 0 | 0 | 0 | 0 | 0 | 0 | 0 | 0 | 0 | 0 | 0 | 0 | 0 | 2829 | 0 | 0 | 1 |
| TKTVLER | 0 | 0 | 0 | 0 | 0 | 0 | 0 | 0 | 0 | 0 | 0 | 0 | 0 | 0 | 1 | 98 | 2578 | 93 | 0 | 1 |
| GWETRME | 0 | 0 | 0 | 0 | 0 | 0 | 0 | 0 | 0 | 0 | 0 | 0 | 39 | 0 | 0 | 702 | 2251 | 43 | 0 | 1 |
| WNQRATG | 0 | 0 | 0 | 0 | 0 | 0 | 0 | 0 | 0 | 0 | 0 | 0 | 0 | 0 | 0 | 0 | 1783 | 0 | 0 | 1 |
| VDMIVPS | 0 | 0 | 0 | 0 | 0 | 0 | 0 | 0 | 0 | 0 | 0 | 0 | 0 | 0 | 4 | 0 | 1613 | 0 | 0 | 1 |
| LPGNRLL | 0 | 0 | 0 | 0 | 0 | 0 | 0 | 0 | 0 | 0 | 0 | 0 | 0 | 0 | 0 | 0 | 1510 | 0 | 0 | 0 |
| QLYREFN | 0 | 0 | 0 | 0 | 0 | 0 | 0 | 0 | 0 | 0 | 0 | 0 | 110 | 0 | 2 | 328 | 1481 | 582 | 0 | 1 |
| GTSTTAQ | 0 | 0 | 0 | 0 | 0 | 0 | 0 | 0 | 0 | 0 | 0 | 0 | 1 | 0 | 0 | 0 | 1137 | 0 | 0 | 1 |
| NTQLHPS | 0 | 0 | 0 | 0 | 0 | 0 | 0 | 0 | 0 | 0 | 0 | 0 | 0 | 0 | 0 | 0 | 1124 | 0 | 0 | 1 |
| YRNHVTY | 0 | 0 | 0 | 0 | 0 | 0 | 0 | 0 | 0 | 0 | 0 | 0 | 0 | 0 | 0 | 0 | 1100 | 0 | 0 | 0 |
| MNSNIPI | 0 | 0 | 0 | 0 | 0 | 0 | 0 | 0 | 0 | 0 | 0 | 0 | 0 | 0 | 0 | 0 | 865 | 0 | 0 | 1 |
| ATLVPAA | 0 | 0 | 0 | 0 | 0 | 0 | 0 | 0 | 0 | 0 | 0 | 0 | 0 | 0 | 0 | 0 | 842 | 0 | 0 | 1 |
| ILAHSIM | 0 | 0 | 0 | 0 | 0 | 0 | 0 | 0 | 0 | 0 | 0 | 0 | 0 | 0 | 1 | 0 | 801 | 0 | 0 | 0 |
| IDGNGTH | 0 | 0 | 0 | 0 | 0 | 0 | 0 | 0 | 0 | 0 | 0 | 0 | 0 | 0 | 0 | 0 | 721 | 0 | 0 | 0 |
| IDNSHTH | 0 | 0 | 0 | 0 | 0 | 0 | 0 | 0 | 0 | 0 | 0 | 0 | 0 | 0 | 0 | 490 | 714 | 0 | 0 | 1 |
| WGRISHV | 0 | 0 | 0 | 0 | 0 | 0 | 0 | 0 | 0 | 0 | 0 | 0 | 0 | 0 | 6 | 0 | 699 | 0 | 0 | 1 |
| HSRAPER | 0 | 0 | 0 | 0 | 0 | 0 | 0 | 0 | 0 | 0 | 0 | 0 | 0 | 0 | 0 | 0 | 0 | 13100 | 0 | 1 |
| TTLGVWT | 0 | 0 | 0 | 0 | 0 | 0 | 0 | 0 | 0 | 0 | 0 | 0 | 0 | 0 | 19 | 0 | 0 | 8649 | 0 | 1 |
| YSEPAVT | 0 | 0 | 0 | 0 | 0 | 0 | 0 | 0 | 0 | 0 | 0 | 0 | 56 | 0 | 1 | 0 | 0 | 6577 | 0 | 1 |
| ESRVMSR | 0 | 0 | 0 | 0 | 0 | 0 | 0 | 0 | 0 | 0 | 0 | 0 | 0 | 8 | 10 | 360 | 0 | 4618 | 0 | 1 |
| GPLHAQF | 0 | 0 | 0 | 0 | 0 | 0 | 0 | 0 | 0 | 0 | 0 | 0 | 0 | 1 | 0 | 0 | 0 | 3220 | 0 | 1 |
| NNTLSRT | 0 | 0 | 0 | 0 | 0 | 0 | 0 | 0 | 0 | 0 | 0 | 0 | 0 | 0 | 0 | 0 | 0 | 2417 | 0 | 1 |
| DHAVPRY | 0 | 0 | 0 | 0 | 0 | 0 | 0 | 0 | 0 | 0 | 0 | 0 | 0 | 0 | 0 | 40 | 0 | 2343 | 0 | 1 |
| AFPPVTA | 0 | 0 | 0 | 0 | 0 | 0 | 0 | 0 | 0 | 0 | 0 | 0 | 0 | 0 | 0 | 0 | 0 | 2038 | 0 | 0 |
| MTVQRGP | 0 | 0 | 0 | 0 | 0 | 0 | 0 | 0 | 0 | 0 | 0 | 0 | 0 | 0 | 0 | 0 | 0 | 2029 | 0 | 0 |
| HLNQQNH | 0 | 0 | 0 | 0 | 0 | 0 | 0 | 0 | 0 | 0 | 0 | 0 | 1 | 0 | 22 | 44 | 0 | 1929 | 0 | 1 |
| QLPFTIK | 0 | 0 | 0 | 0 | 0 | 0 | 0 | 0 | 0 | 0 | 0 | 0 | 0 | 0 | 0 | 0 | 0 | 1840 | 0 | 1 |
| SQPTWMF | 0 | 0 | 0 | 0 | 0 | 0 | 0 | 0 | 0 | 0 | 0 | 0 | 0 | 0 | 0 | 0 | 0 | 1710 | 0 | 1 |
| YNGSANQ | 0 | 0 | 0 | 0 | 0 | 0 | 0 | 0 | 0 | 0 | 0 | 0 | 57 | 0 | 1 | 19 | 0 | 1694 | 0 | 1 |
| TTQVLEA | 0 | 0 | 0 | 0 | 0 | 0 | 0 | 0 | 0 | 0 | 0 | 0 | 10 | 0 | 103 | 1413 | 6 | 1693 | 0 | 1 |
| FSLQTTR | 0 | 0 | 0 | 0 | 0 | 0 | 0 | 0 | 0 | 0 | 0 | 0 | 0 | 0 | 0 | 0 | 0 | 1415 | 0 | 1 |
| GLRNPPS | 0 | 0 | 0 | 0 | 0 | 0 | 0 | 0 | 0 | 0 | 0 | 0 | 18 | 0 | 0 | 2 | 0 | 1114 | 0 | 1 |
| TEKFRVT | 0 | 0 | 0 | 0 | 0 | 0 | 0 | 0 | 0 | 0 | 0 | 0 | 0 | 0 | 0 | 0 | 0 | 882 | 0 | 1 |
| TVISQNM | 0 | 0 | 0 | 0 | 0 | 0 | 0 | 0 | 0 | 0 | 0 | 0 | 8 | 0 | 0 | 15 | 0 | 854 | 0 | 1 |
| AFTTSYM | 0 | 0 | 0 | 0 | 0 | 0 | 0 | 0 | 0 | 0 | 0 | 0 | 0 | 0 | 0 | 0 | 0 | 727 | 0 | 1 |
| MSVITKP | 0 | 0 | 0 | 0 | 0 | 0 | 0 | 0 | 0 | 0 | 0 | 0 | 20 | 26 | 6 | 253 | 19 | 688 | 0 | 0 |
| Total Reads | 18718 | 25065 | 31095 | 18591 | 15878 | 40390 | 17387 | 25513 | 35198 | 15350 | 16656 | 35198 | 64196 | 131881 | 57009 | 58445 | 66750 | 91056 | | |

**Table S2**. Copy number for top 20 sequences from each round and replicate of selection using both BA and EmA methods. The color for each selection is used to distinguish between different screens. The numbers indicate the copy number of each sequence. The sequences are ordered by decreasing copy number in the corresponding screen. If sequences are found in other screens, the corresponding copy number is indicated. The last two columns identify if the sequence is a parasite ('1') or not ('0') for the two studied lots. The total number of reads for each screen is indicated.

| | Lot2-BA-Rep1-Round1 | Lot2-BA-Rep2-Round1 | Lot2-BA-Rep3-Round1 | Found in MDA-MB-231 screen | Lot2-Parasites |
|---|---|---|---|---|---|
| SLLGQTP | 2917 | 1 | 26 | 0 | 1 |
| WSSHKNV | 1649 | 0 | 0 | 0 | 0 |
| AYVARQN | 1445 | 584 | 0 | 0 | 1 |
| RPPLINT | 712 | 0 | 0 | 0 | 1 |
| SYTDLLR | 654 | 0 | 0 | 0 | 0 |
| LATLTAC | 586 | 0 | 0 | 0 | 1 |
| IWKQALI | 579 | 0 | 0 | 0 | 0 |
| VVYVSFF | 480 | 0 | 0 | 0 | 1 |
| INTLSRT | 457 | 0 | 0 | 0 | 1 |
| TVISQNM | 455 | 0 | 0 | ** | 1 |
| AHKHDPT | 451 | 0 | 0 | 0 | 1 |
| ILDKLTH | 435 | 0 | 0 | 0 | 0 |
| VDIGDSR | 376 | 0 | 0 | 0 | 1 |
| FQWELYS | 359 | 0 | 0 | 0 | 1 |
| STIANTK | 357 | 0 | 0 | 0 | 0 |
| SGAVWPF | 326 | 0 | 131 | 0 | 1 |
| LTSVAGA | 303 | 0 | 0 | 0 | 0 |
| NWKQALI | 297 | 0 | 0 | 0 | 0 |
| DVYVSFF | 287 | 0 | 0 | 0 | 0 |
| QATLTAC | 284 | 0 | 0 | 0 | 1 |
| FQVALHS | 264 | 0 | 0 | 0 | 0 |
| LGGVRLY | 261 | 14 | 49 | 0 | 1 |
| LVVHRTS | 257 | 0 | 0 | 0 | 1 |
| MDGAAGF | 234 | 0 | 69 | 0 | 1 |
| NNTLSRT | 232 | 0 | 0 | ** | 1 |
| GMNTTWT | 228 | 0 | 79 | 0 | 1 |
| NLDKLTH | 227 | 0 | 0 | 0 | 0 |
| IKSAWFL | 220 | 218 | 0 | 0 | 0 |
| SLLGQTP | 214 | 0 | 1 | 0 | 1 |
| GQHVWWI | 213 | 0 | 0 | 0 | 0 |
| VPGLGLN | 208 | 0 | 0 | 0 | 1 |
| VFKINSK | 205 | 0 | 0 | 0 | 0 |
| WSSLKNV | 200 | 0 | 0 | 0 | 0 |
| SFNLPNT | 188 | 0 | 0 | 0 | 1 |
| AYVARQK | 181 | 61 | 0 | 0 | 1 |
| LHHKVTI | 181 | 0 | 0 | 0 | 0 |
| YQWELYS | 180 | 0 | 0 | 0 | 1 |
| SSSSHVM | 176 | 0 | 0 | 0 | 1 |
| VHAGLQV | 175 | 16 | 0 | 0 | 1 |
| EPGLGLN | 169 | 0 | 0 | 0 | 1 |
| AIDFARN | 168 | 9 | 0 | 0 | 1 |
| LLADMHA | 168 | 0 | 0 | 0 | 1 |
| MTGSVQM | 161 | 0 | 0 | 0 | 0 |
| ILLPFAS | 161 | 0 | 0 | 0 | 0 |
| IDRTQFM | 156 | 105 | 6 | 0 | 1 |
| LMHREPA | 156 | 0 | 0 | 0 | 1 |
| GLRNPPS | 145 | 27 | 0 | ** | 1 |
| IMENSYV | 144 | 0 | 0 | 0 | 0 |
| VALLSTT | 139 | 0 | 0 | 0 | 0 |
| DHAGLQV | 135 | 9 | 0 | 0 | 1 |

| | Lot2-BA-Rep1-Round1 | Lot2-BA-Rep2-Round1 | Lot2-BA-Rep3-Round1 | Found in MDA-MB-231 screen | Lot2-Parasites |
|---|---|---|---|---|---|
| LSHTLTW | 39 | 7429 | 5 | 0 | 1 |
| HSHTLTW | 20 | 4589 | 3 | 0 | 1 |
| TTANVRI | 1 | 2461 | 0 | 0 | 1 |
| GTSIYLH | 0 | 2036 | 2 | 0 | 1 |
| LRSDPVV | 0 | 1795 | 670 | ** | 1 |
| AHINVPS | 0 | 1776 | 11 | 0 | 1 |
| AGMWNAT | 0 | 1266 | 0 | 0 | 1 |
| LSHSLTW | 8 | 904 | 0 | 0 | 1 |
| VENVHVR | 0 | 705 | 0 | ** | 1 |
| HSHTLTC | 7 | 650 | 1 | 0 | 1 |
| FNGSANQ | 15 | 594 | 2 | 0 | 1 |
| AYVARQN | 1445 | 584 | 0 | 0 | 1 |
| WEIGSGP | 0 | 580 | 0 | 0 | 1 |
| GQGQTIP | 0 | 540 | 0 | 0 | 1 |
| LLYREFN | 33 | 518 | 808 | 0 | 1 |
| AEIPHRG | 0 | 515 | 0 | 0 | 0 |
| YNGSANQ | 5 | 508 | 4 | ** | 1 |
| FMTAPGF | 0 | 371 | 0 | 0 | 0 |
| STMKTGC | 0 | 354 | 0 | 0 | 0 |
| FARANAA | 0 | 352 | 0 | 0 | 1 |
| HSHTLTR | 4 | 343 | 0 | 0 | 1 |
| GPLHAQF | 20 | 326 | 0 | 0 | 1 |
| VTAVKTG | 0 | 305 | 0 | 0 | 1 |
| VPQILPY | 0 | 303 | 21 | 0 | 1 |
| AHIYVPS | 0 | 300 | 2 | 0 | 0 |
| LTMPFSG | 0 | 296 | 23 | 0 | 1 |
| AHTSTWP | 0 | 286 | 0 | 0 | 1 |
| QLYREFN | 26 | 283 | 380 | ** | 1 |
| SPTMVNA | 0 | 278 | 0 | 0 | 0 |
| HSHTLTF | 1 | 266 | 0 | 0 | 1 |
| GTSFYLH | 0 | 265 | 0 | 0 | 1 |
| WGRISHV | 26 | 262 | 1436 | ** | 1 |
| LLYFAAP | 0 | 257 | 1 | 0 | 1 |
| LANATSL | 2 | 256 | 0 | 0 | 1 |
| TTAYVRI | 0 | 244 | 0 | 0 | 0 |
| IAYGGRI | 0 | 243 | 0 | 0 | 1 |
| VQYKPMK | 0 | 241 | 129 | 0 | 1 |
| WQAHGIS | 0 | 241 | 0 | 0 | 0 |
| ASDGKVA | 0 | 224 | 0 | 0 | 0 |
| IKSAWFL | 220 | 218 | 0 | 0 | 0 |
| WPNKFMY | 0 | 216 | 0 | 0 | 0 |
| TTANVRI | 1 | 212 | 0 | 0 | 1 |
| SYASYYL | 0 | 209 | 0 | 0 | 0 |
| THEDVNL | 0 | 205 | 0 | 0 | 0 |
| MRTSMPH | 0 | 191 | 0 | 0 | 0 |
| SHVNVPS | 1 | 187 | 0 | 0 | 1 |
| VKNVPSQ | 0 | 187 | 0 | 0 | 0 |
| VSRVMSR | 0 | 185 | 0 | 0 | 1 |
| AYDKFTP | 0 | 184 | 0 | 0 | 1 |
| TTQVLEA | 115 | 183 | 155 | ** | 1 |

| | Lot2-BA-Rep1-Round1 | Lot2-BA-Rep2-Round1 | Lot2-BA-Rep3-Round1 | Found in MDA-MB-231 screen | Lot2-Parasites |
|---|---|---|---|---|---|
| SAAWNKS | 0 | 0 | 14055 | 0 | 1 |
| WSLSELH | 47 | 168 | 8439 | ** | 1 |
| SLDVRMW | 1 | 49 | 5970 | ** | 1 |
| VPWSVTR | 0 | 0 | 2644 | 0 | 0 |
| ALPAKRE | 0 | 0 | 2548 | 0 | 0 |
| TATLPRM | 0 | 0 | 1920 | 0 | 1 |
| LQLAVDT | 0 | 0 | 1882 | 0 | 1 |
| WGRISHV | 26 | 262 | 1436 | ** | 1 |
| WDPRVNV | 0 | 0 | 1311 | 0 | 1 |
| FQLPWAG | 0 | 0 | 1260 | 0 | 0 |
| GMHETHV | 0 | 0 | 1146 | 0 | 1 |
| LSPSLRN | 0 | 0 | 1104 | 0 | 1 |
| LWTNNWL | 0 | 0 | 1046 | 0 | 0 |
| TKPNLYN | 54 | 0 | 969 | 0 | 1 |
| QQLAVDT | 0 | 0 | 838 | 0 | 1 |
| LLYREFN | 33 | 518 | 808 | 0 | 1 |
| MFDMVKL | 26 | 0 | 792 | 0 | 1 |
| SAAWNKS | 0 | 0 | 747 | 0 | 1 |
| LAAWHFI | 0 | 0 | 741 | 0 | 1 |
| LTWLEMG | 72 | 0 | 715 | 0 | 1 |
| AGHNLVP | 0 | 0 | 709 | 0 | 0 |
| LRSDPVV | 0 | 1795 | 670 | ** | 1 |
| LRAQVTP | 0 | 0 | 583 | 0 | 1 |
| WSLSELQ | 4 | 16 | 578 | 0 | 1 |
| SLLSLNF | 0 | 0 | 546 | 0 | 0 |
| HWTNNWL | 0 | 0 | 489 | 0 | 0 |
| KFDMVKL | 21 | 0 | 473 | 0 | 1 |
| LHRPANC | 0 | 0 | 461 | 0 | 0 |
| TVNFKLY | 2 | 0 | 419 | 0 | 1 |
| QLYREFN | 26 | 283 | 380 | ** | 1 |
| SLDVRMC | 0 | 1 | 357 | 0 | 0 |
| QTWLEMG | 38 | 0 | 355 | ** | 1 |
| SAAWNKT | 0 | 0 | 355 | 0 | 0 |
| TVGNSVG | 0 | 0 | 352 | 0 | 0 |
| HAAWHFI | 0 | 0 | 341 | 0 | 1 |
| AKSLMFQ | 0 | 2 | 317 | 0 | 1 |
| LGEWIKY | 0 | 0 | 295 | 0 | 1 |
| FASAARV | 0 | 0 | 293 | 0 | 1 |
| SFSRAES | 9 | 16 | 289 | 0 | 1 |
| GKVAKQE | 0 | 0 | 259 | 0 | 1 |
| QQLAVDT | 0 | 0 | 234 | 0 | 1 |
| LPVRLDW | 0 | 0 | 230 | ** | 1 |
| LTPPNDF | 0 | 0 | 218 | 0 | 0 |
| VHYVKGP | 0 | 0 | 198 | 0 | 0 |
| IDNSHTH | 0 | 1 | 189 | ** | 1 |
| LKHSHHY | 0 | 0 | 182 | 0 | 0 |
| TLWGEHP | 0 | 0 | 181 | 0 | 0 |
| YASAARV | 0 | 0 | 181 | 0 | 1 |
| VPWSVTR | 0 | 0 | 179 | 0 | 0 |
| WSHTPPR | 0 | 0 | 176 | 0 | 0 |

**Table S3**. Top 50 sequences from three replicates of one round of selection against HEK cell line using the BA method. The numbers indicate the copy number of each of the top 50 sequences. The sequences are sorted by copy number in the corresponding screen. If sequences were found in other screens, the corresponding copy number is indicated. The last column identifies the sequence as a parasite ('1') or not ('0'), additionally, all parasite sequences are colored red. Asterisks '**' indicate sequences that we identified and validated to be hits in the top 20 against the MDA-MB-231 cell line.

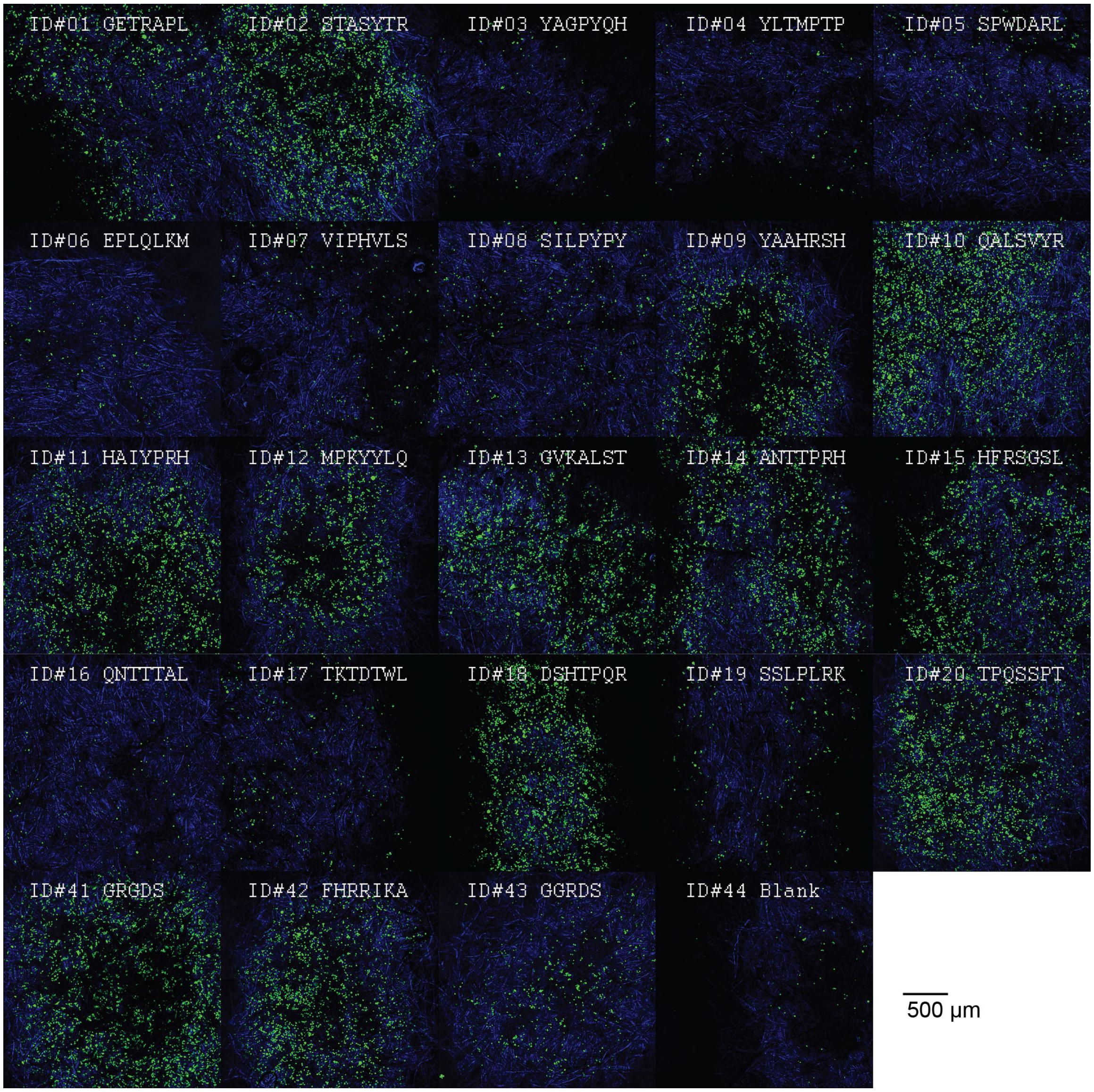


**Figure S5**. Representative confocal images of MDA-MB-231-GFP cells on peptide array after short-term adhesion (3 hours). Peptides were selected in a BA screen. Confocal images correspond to the fluorescent gel image represented in Figure 1C and Figure S2A. All images were acquired using identical microscopy settings (laser intensity, PMT gain). Colors: blue – paper fibers (imaged by reflectance); green – GFP.

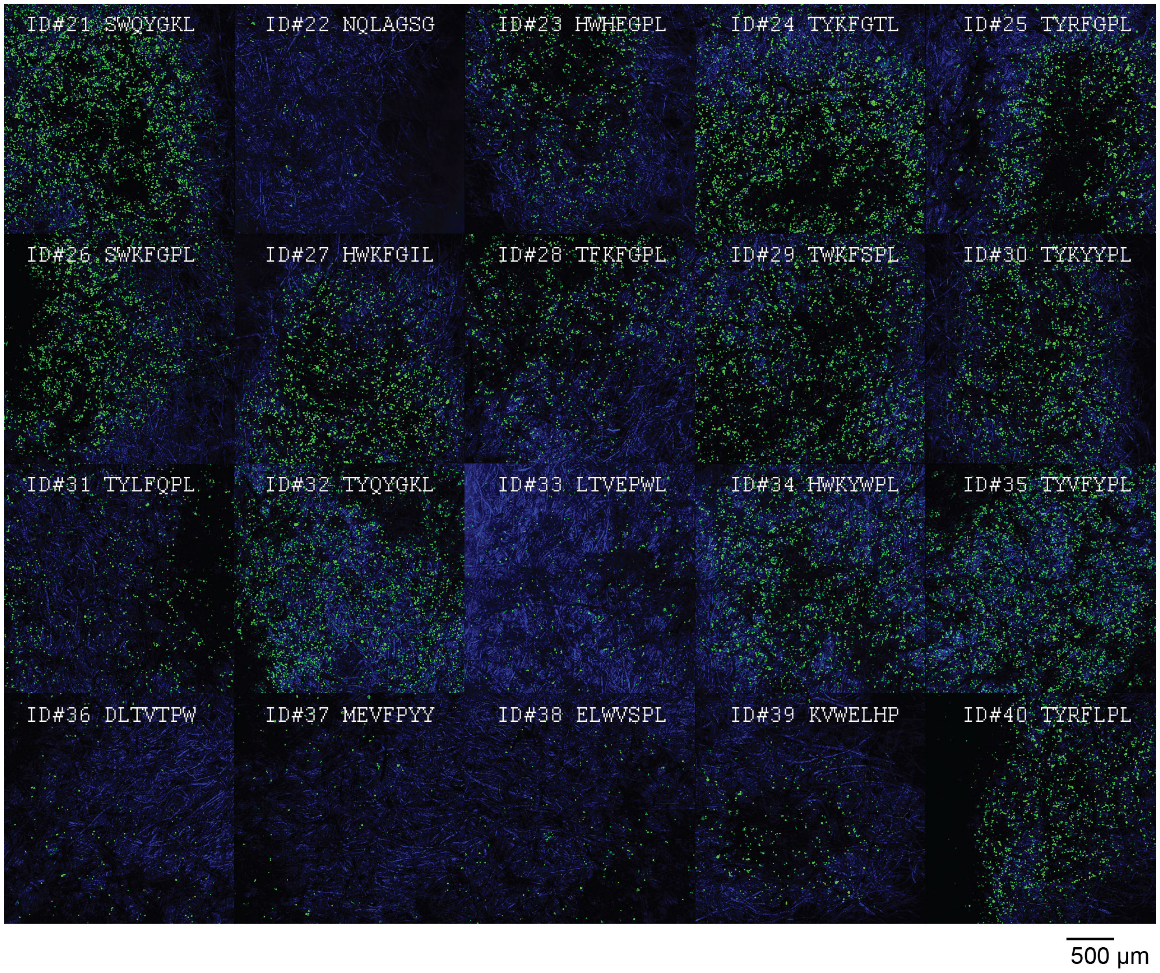


**Figure S6**. Representative confocal images of MDA-MB-231-GFP cells on peptide array after short-term adhesion (3 hours). Peptides were selected in a EmA screen. Confocal images correspond to the fluorescent gel image represented in Figure 1C and Figure S2A. All images were acquired using identical microscopy settings (laser intensity, PMT gain). Colors: blue – paper fibers (imaged by reflectance); green – GFP.

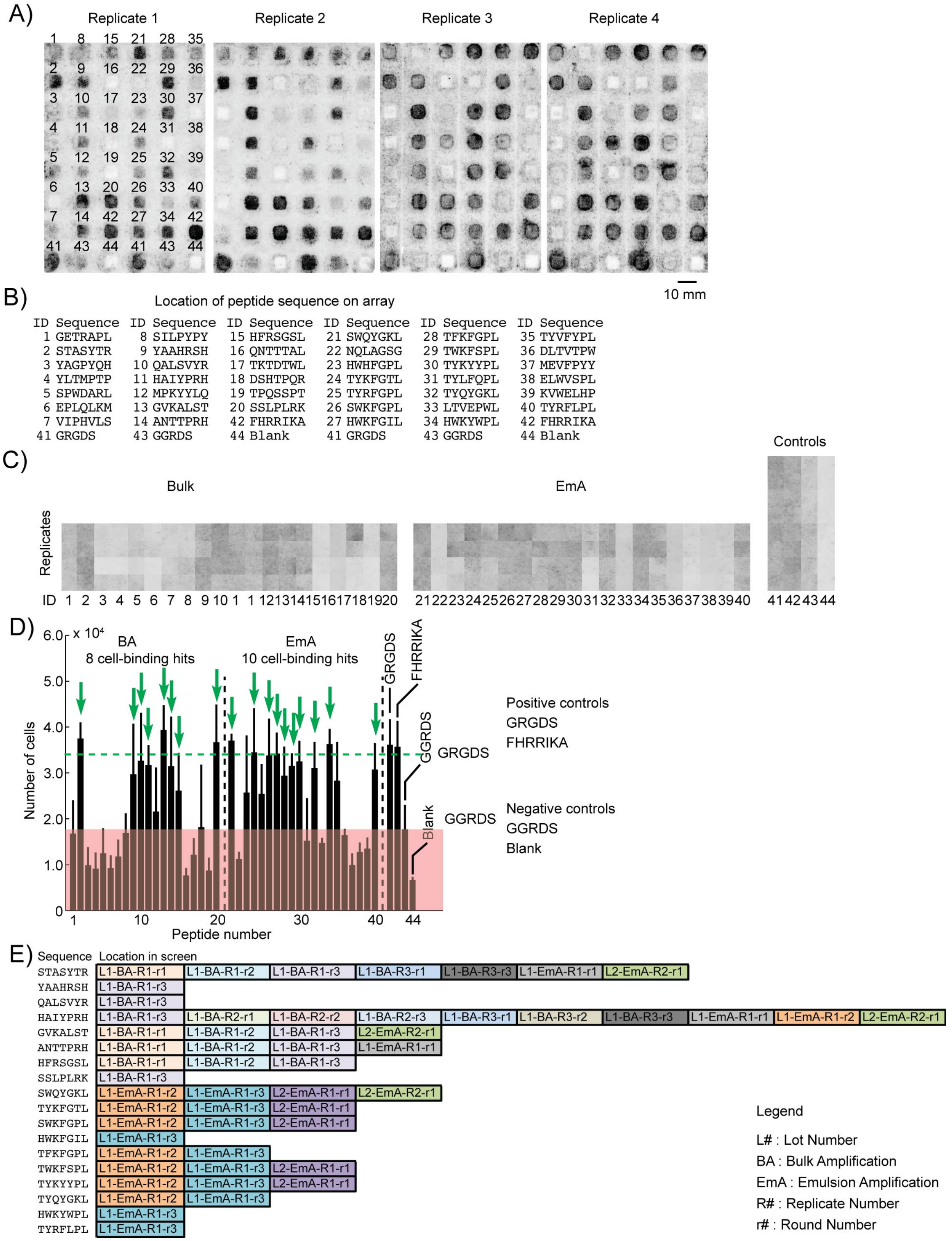


**Figure S7**. Work flow of cell adhesion analysis. (A) Top 20 sequences identified from selection with BA and EmA were synthesized on paper. MDA-MB-231-GFP cells were seeded onto the array and incubated at 37°C in a $CO_2$ incubator. After 3 hours we imaged the arrays with a fluorescent gel scanner. Grey-scale intensities were adjusted in this figure to simplify visualization (same level for all images). Original .gel files were used in processing without any adjustments to grey-scale intensities. (B) The location of each peptide sequence. (C) Matlab script identified the middle of each peptide zone, extracted the grey-scale intensity and organized the replicates. (D) The average greyscale intensity from each replicate was converted to the number of cells using a calibration curve (Figure S8). Data represent an average from 4-8 experiments; error bar is one standard deviation. Cell-binding hits were peptides that supported adhesion of significantly higher ($p < 0.05$) number of cells than the negative control (GGRDS, red box). Green arrows indicate eighteen cell-binding hits. (E) Some cell-binding peptide hit were identified in multiple screens (the color and abbreviation are identical to those used in Table S1.

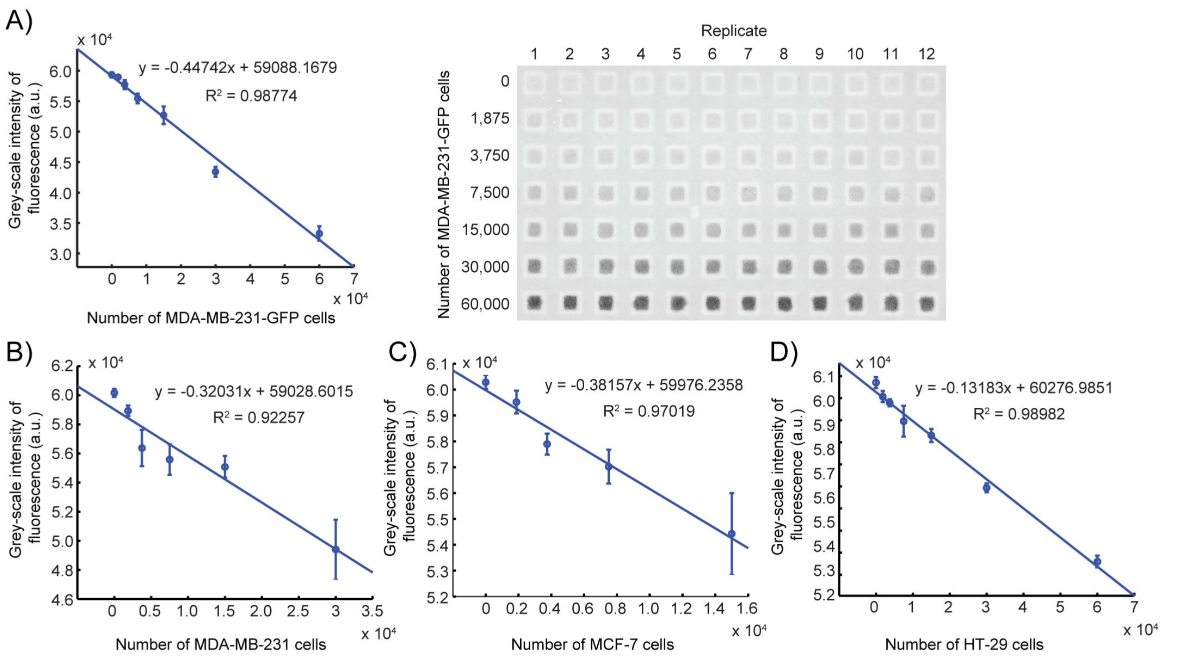


**Figure S8**. Calibration curves used for extrapolating number of cells from grey-scale intensity. (A) Calibration curve converting the number of MDA-MB-231-GFP cells to the corresponding grey-scale intensity per peptide zone (A=0.16 cm$^2$) measured by gel scanner. The curve was generated by seeding a predefined number of cells inside paper-based array; after a brief incubation (15 minutes), the arrays were imaged by gel scanner. (B-D) Standard curves for the number of MDA-MB-231, MCF-7, and HT-29 cell lines stained with 4 μM Cell Tracker Green (1-hour prior the cell adhesion assay) and the corresponding grey-scale intensity per peptide zone (A=0.16 cm$^2$) used in each array. Substrates were scanned using a Typhoon FLA9500 fluorescent gel scanner; with the following settings: LPB filter, 50 μm resolution, 400 V PMT.

A)

| | | | | | |
|---|---|---|---|---|---|
| TPPTMDH | SPQMTLS | QPSMLNP | MTVQPWP | HHSLTVT | AASSLTI |
| TTNLSPW | SPTQPKS | QPTHPTR | NHWASPR | HIPPGSP | ALAHRIL |
| TVRHLQL | SSALLLP | QRLPQTA | QATHRSH | IPTLPSS | ALEVTFW |
| TWYFGPL | SSLVRTA | QSLALQP | QEPLTAR | KAVHPLR | AMSSRSL |
| TYQNPVH | SSSVVTH | QTGYATR | QLHNDAT | LEVYALV | GKPMPPM |
| GGRDSAA | STFTKSP | QTSMATV | QLMNASR | MDAHHAL | GKVQAQS |
| GRGDSAA | STPMQNL | SHTAPLR | QLTVMSW | MHAPPFY | GPMLARG |
| FHRRIKA | TARYPSW | SLSLIQT | QPPRSTS | MPGSLPS | HALGPSS |

B)

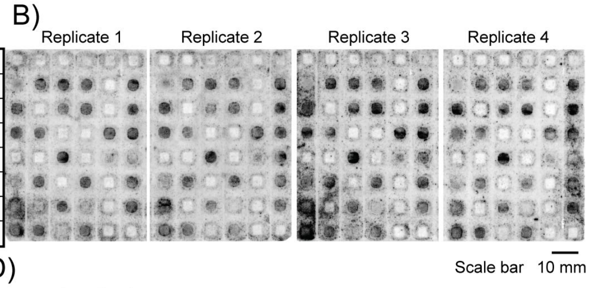


C)

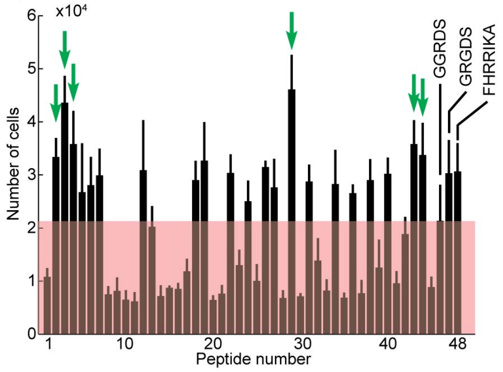


D)

| Sequence | Location in screen | | |
|---|---|---|---|
| ALAHRIL | L1-BA-R2-r1 | L1-BA-R2-r2 | L1-BA-R2-r3 |
| ALEVTFW | L1-EmA-R1-r2 | | |
| AMSSRSL | L1-BA-R2-r2 | L1-EmA-R1-r1 | |
| QTGYATR | L1-EmA-R1-r1 | | |
| TVRHLQL | L1-BA-R1-r1 | L1-BA-R1-r2 | |
| TWYFGPL | L1-BA-R2-r1 | L1-BA-R2-r2 | |

**Figure S9**. Testing short-term cell adhesion of top 20 peptides from BA and EmA selections on a second peptide array. (A) Peptide sequences tested for cell-adhesion. (B) Representative fluorescent gel scanner images of MDA-MB-231-GFP adhering to peptide array (after 3 hours). Grey-scale intensities were adjusted in this figure to simplify visualization (same level for all images). Original .gel files were used in processing without and adjustments to grey-scale intensities. (C) The average greyscale intensity from each replicate was converted to the number of cells using a calibration curve (Figure S8). Data represent an average from 4 experiments; error bar is one standard deviation. Cell-binding hits were peptides that supported adhesion of significantly higher ($p < 0.05$) number of cells than the negative control (GGRDS, red box). Green arrows indicate eighteen cell-binding hits. (D) List of cell-binding peptide hits and their location in the screen(s). The abbreviation and color for each screen is the same as in Table S1.

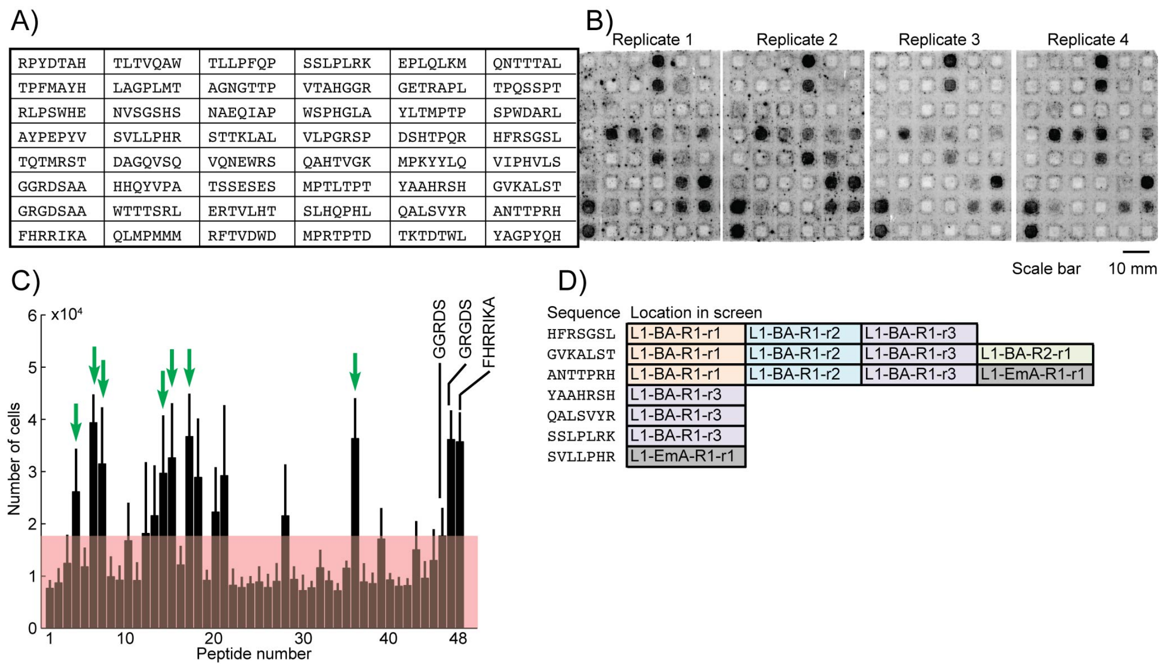

| | | | | | |
|---|---|---|---|---|---|
| RPYDTAH | TLTVQAW | TLLPFQP | SSLPLRK | EPLQLKM | QNTTTAL |
| TPFMAYH | LAGPLMT | AGNGTTP | VTAHGGR | GETRAPL | TPQSSPT |
| RLPSWHE | NVSGSHS | NAEQIAP | WSPHGLA | YLTMPTP | SPWDARL |
| AYPEPYV | SVLLPHR | STTKLAL | VLPGRSP | DSHTPQR | HFRSGSL |
| TQTMRST | DAGQVSQ | VQNEWRS | QAHTVGK | MPKYYLQ | VIPHVLS |
| GGRDSAA | HHQYVPA | TSSESES | MPTLTPT | YAAHRSH | GVKALST |
| GRGDSAA | WTTTSRL | ERTVLHT | SLHQPHL | QALSVYR | ANTTPRH |
| FHRRIKA | QLMPMMM | RFTVDWD | MPRTPTD | TKTDTWL | YAGPYQH |



**Figure S10**. Testing short-term cell adhesion of top 20 peptides from BA and EmA selections on a third peptide array. (A) Peptide sequences tested for cell-adhesion. (B) Representative fluorescent gel scanner images of MDA-MB-231-GFP adhering to peptide array (after 3 hours). Grey-scale intensities were adjusted in this figure to simplify visualization (same level for all images). Original .gel files were used in processing without and adjustments to grey-scale intensities. (C) The average greyscale

A)

| | | | | | |
|---|---|---|---|---|---|
| ELGTTQT | SSVSWLN | WSLSELH | GWRTTWP | QNMQQQI | TPMTRAL |
| AMTALDL | TTQVLEA | LPVRLDW | ATHQRPA | AWSAVMR | VMSQPHP |
| AGSVIDT | QFTQLHQ | QTWLEMG | WMASMAV | APIWMHV | GSTVFTA |
| QAYHVSA | DAIPTSV | GPHNPTQ | HEQPMHR | ATWQLGT | EQGRPLP |
| SNMTRWH | VGKTSFQ | NDRPHMP | SLDVRMW | LHRQSSA | MAANGAR |
| GGRDSAA | GQVALLD | VPNIVTQ | QHMPLTR | ASWIPLP | QVLLTAA |
| GRGDSAA | VENVHVR | LRSDPVV | TMTEHRQ | QQQYMAH | AGRELCC |
| FHRRIKA | VPVTMYW | VPASPWT | STPATLI | STAMDGR | ARAVLQL |

B)

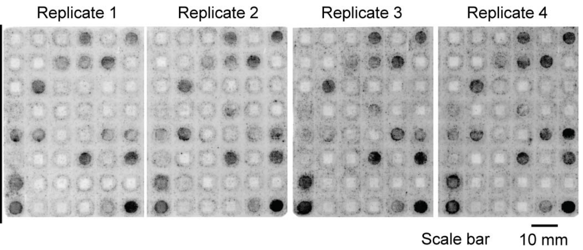


C)

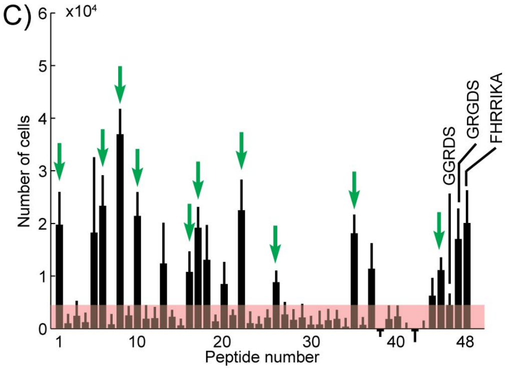


D)

| Sequence | Location in screen |
|---|---|
| TPMTRAL | L2-EmA-R2-r1 |
| QVLLTAA | L2-EmA-R3-r1 |
| ARAVLQL | L2-EmA-R3-r1 |
| AWSAVMR | L2-EmA-R3-r1 |
| STAMDGR | L2-EmA-R3-r1 |
| GWRTTWP | L2-EmA-R3-r1 |
| QHMPLTR | L2-EmA-R3-r1 |
| LPVRLDW | L2-BA-R1-r1 |
| QFTQLHQ | L2-BA-R1-r1 |
| SNMTRWH | L2-BA-R2-r1 |

**Figure S11**. Testing short-term cell adhesion of top 20 peptides from BA and EmA selections on a forth peptide array. (A) Peptide sequences tested for cell-adhesion. (B) Representative fluorescent gel scanner images of MDA-MB-231-GFP adhering to peptide array (after 3 hours). Grey-scale intensities were adjusted in this figure to simplify visualization (same level for all images). Original .gel files were used in processing without and adjustments to grey-scale intensities. (C) The average greyscale intensity from each replicate was converted to the number of cells using a calibration curve (Figure S8). Data represent an average from 4 experiments; error bar is one standard deviation. Cell-binding hits were peptides that supported adhesion of significantly higher ($p < 0.05$) number of cells than the negative control (GGRDS, red box). Green arrows indicate eighteen cell-binding hits. (D) List of cell-binding peptide hits and their location in the screen(s). The abbreviation and color for each screen is the same as in Table S1.

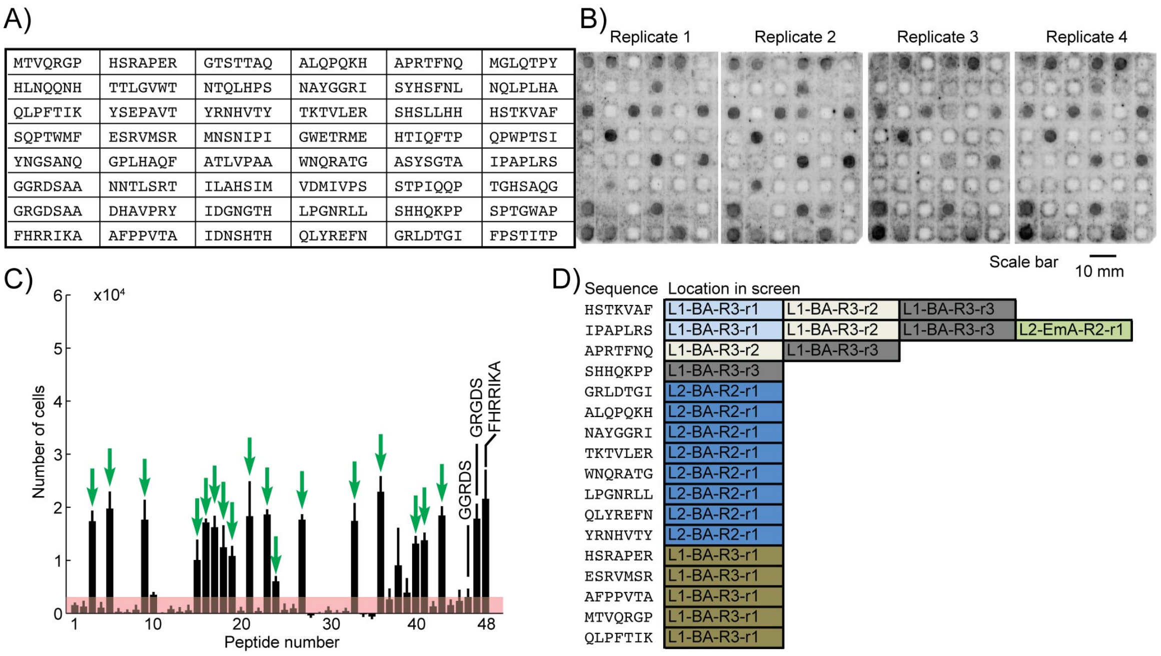


**Figure S12**. Testing short-term cell adhesion of top 20 peptides from BA and EmA selections on a fifth peptide array. (A) Peptide sequences tested for cell-adhesion. (B) Representative fluorescent gel scanner images of MDA-MB-231-GFP adhering to peptide array (after 3 hours). Grey-scale intensities were adjusted in this figure to simplify visualization (same level for all images). Original .gel files were used in processing without and adjustments to grey-scale intensities. (C) The average greyscale intensity from each replicate was converted to the number of cells using a calibration curve (Figure S8). Data represent an average from 4 experiments; error bar is one standard deviation. Cell-binding hits were peptides that supported adhesion of significantly higher ($p < 0.05$) number of cells than the negative control (GGRDS, red box). Green arrows indicate eighteen cell-binding hits. (D) List of cell-binding peptide hits and their location in the screen(s). The abbreviation and color for each screen is the same as in Table S1.

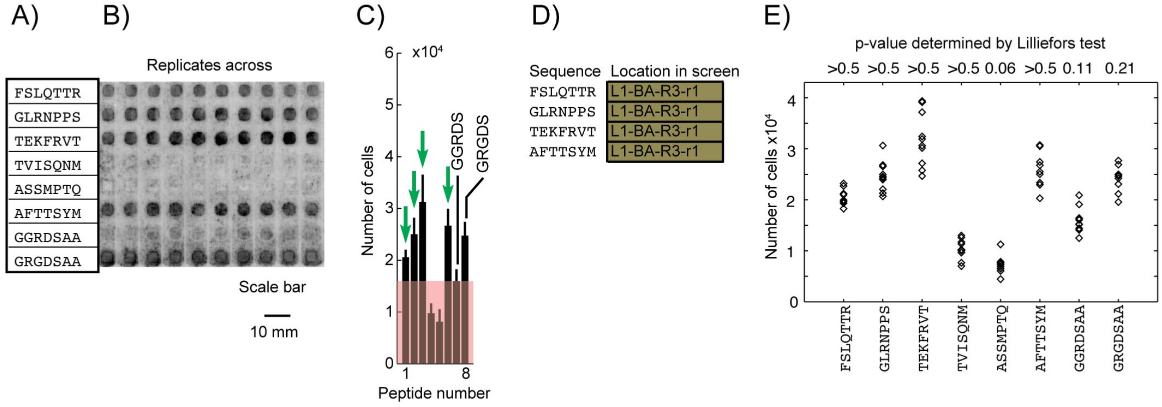


**Figure S13**. Adhesion of cells to peptide array is reproducible in 10 replicates. (A) Testing short-term cell adhesion of top 20 peptides from BA and EmA selections on a sixth peptide array. (B) Representative fluorescent gel scanner images of MDA-MB-231-GFP adhering to peptide array (after 3 hours). Grey-scale intensities were adjusted in this figure to simplify visualization (same level for all images). Original .gel files were used in processing without and adjustments to grey-scale intensities. (C) The average greyscale intensity from each replicate was converted to the number of cells using a calibration curve (Figure S8). Data represent an average from 10 experiments; error bar is one standard deviation. Cell-binding hits were peptides that supported adhesion of significantly higher ($p < 0.05$) number of cells than the negative control (GGRDS, red box). Green arrows indicate eighteen cell-binding hits. (D) List of cell-binding peptide hits and their location in the screen(s). The abbreviation and color for each screen is the same as in Table S1. (E) Cell adhesion data for each peptide was tested for normality using the Lilliefors test. P-values > 0.05 indicate data originates from a normally distributed population.

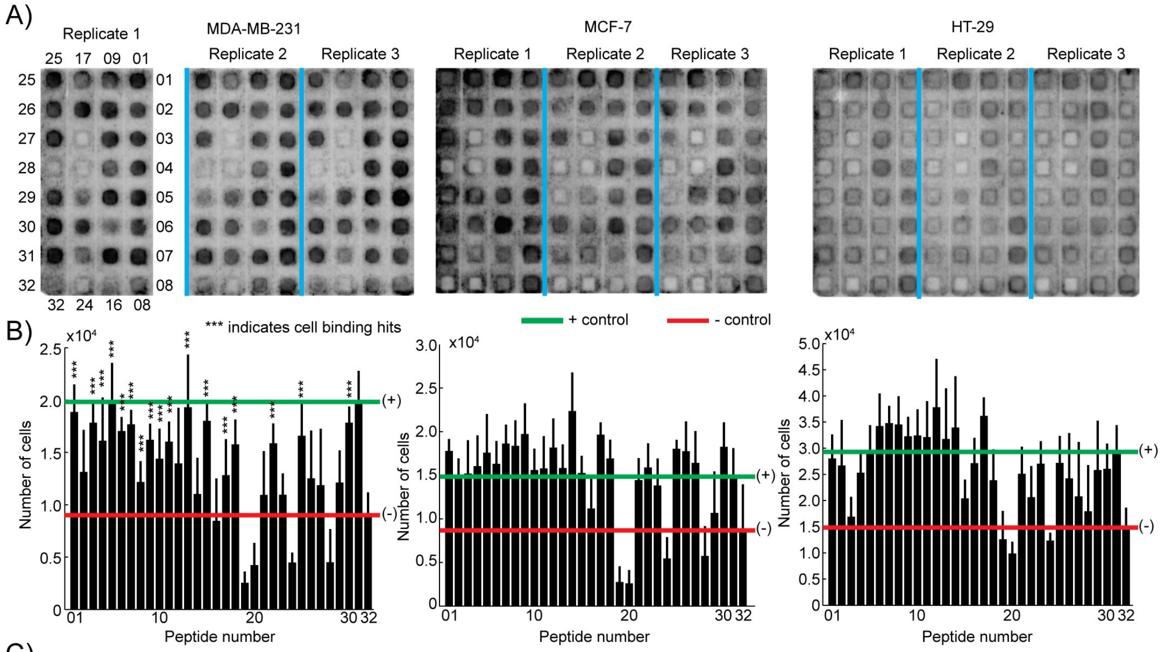


C)

Location of peptide sequence on array

| ID | Sequence | ID | Sequence | ID | Sequence | ID | Sequence |
|---|---|---|---|---|---|---|---|
| 01 | YAAHRSH | 09 | TWKFSPL | 17 | TWYFGPL | 25 | SNMTRWH |
| 02 | QALSVYR | 10 | TYKYYPL | 18 | SVLLPHR | 26 | HSTKVAF |
| 03 | ANTTPRH | 11 | TYQYGKL | 19 | QVLLTAA | 27 | APRTFNQ |
| 04 | HFRSGSL | 12 | TYRFLPL | 20 | STAMDGR | 28 | SHHQKPP |
| 05 | SSLPLRK | 13 | ALAHRIL | 21 | GWRTTWP | 29 | GRLDTGI |
| 06 | SWKFGPL | 14 | ALEVTFW | 22 | QHMPLTR | 30 | ALQPQKH |
| 07 | HWKFGIL | 15 | QTGYATR | 23 | LPVRLDW | 31 | GRGDSAA |
| 08 | TFKFGPL | 16 | TVRHLQL | 24 | QFTQLHQ | 32 | GGRDSAA |

**Figure S14.** Representative images of the peptide arrays tested for adhesion of MDA-MB-231, MCF-7, and HT-29 cells on the first focused array of cell-binding hits (A). Cell lines were stained for one hour with 4 μM Cell Tracker green in MEM media prior to the assay. (B) Plots of the average number of cells per peptide zone. The number of cells was calculated using calibration curves for each cell line (Figure S4). Data represent an average from 6 experiments; error bar is one standard deviation. Asterisks (***) indicate peptides binding to significantly more MDA-MB-231 cells than the negative control (GGRDSAA). (C) Location of each peptide sequence on the peptide arrays.

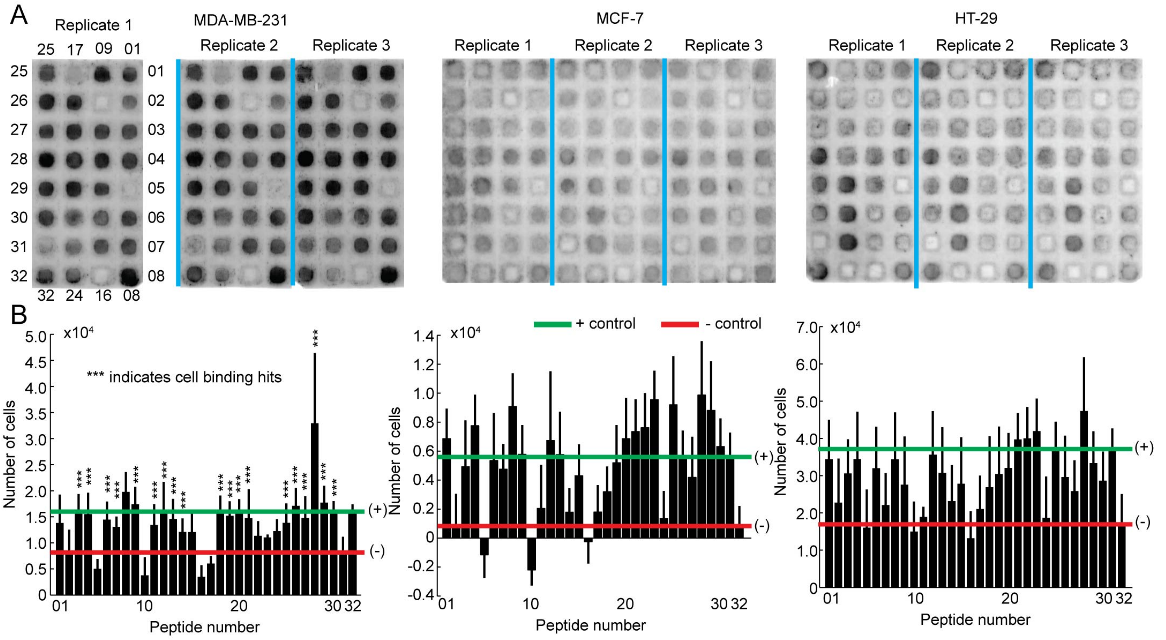


C

Location of peptide sequence on array

| ID | Sequence | ID | Sequence | ID | Sequence | ID | Sequence |
|---|---|---|---|---|---|---|---|
| 01 | NAYGGRI | 09 | ESRVMSR | 17 | AFTTSYM | 25 | HWKYWPL |
| 02 | TKTVLER | 10 | AFPPVTA | 18 | STASYTR | 26 | AMSSRSL |
| 03 | WNQRATG | 11 | MTVQRGP | 19 | HAIYPRH | 27 | TPMTRAL |
| 04 | LPGNRLL | 12 | QLPFTIK | 20 | GVKALST | 28 | ARAVLQL |
| 05 | QLYREFN | 13 | FSLQTTR | 21 | SWQYGKL | 29 | AWSAVMR |
| 06 | YRNHVTY | 14 | GLRNPPS | 22 | TYKFGTL | 30 | IPAPLRS |
| 07 | HSRAPER | 15 | TEKFRVT | 23 | TYRFGPL | 31 | GGRDSAA |
| 08 | FHRRIKA | 16 | MQLPLAT | 24 | VRPMPLQ | 32 | GRGDSAA |

**Figure S15.** Representative images of the peptide arrays tested for adhesion of MDA-MB-231, MCF-7, and HT-29 cells on the second focused array of cell-binding hits (A). Cell lines were stained for one hour with 4 µM Cell Tracker green in MEM media prior to the assay. (B) Plots of the average number of cells per peptide zone. The number of cells was calculated using calibration curves for each cell line (Figure S4). Data represent an average from 6 experiments; error bar is one standard deviation. Asterisks (***) indicate peptides binding to significantly more MDA-MB-231 cells than the negative control (GGRDSAA). (C) Location of each peptide sequence on the peptide arrays.

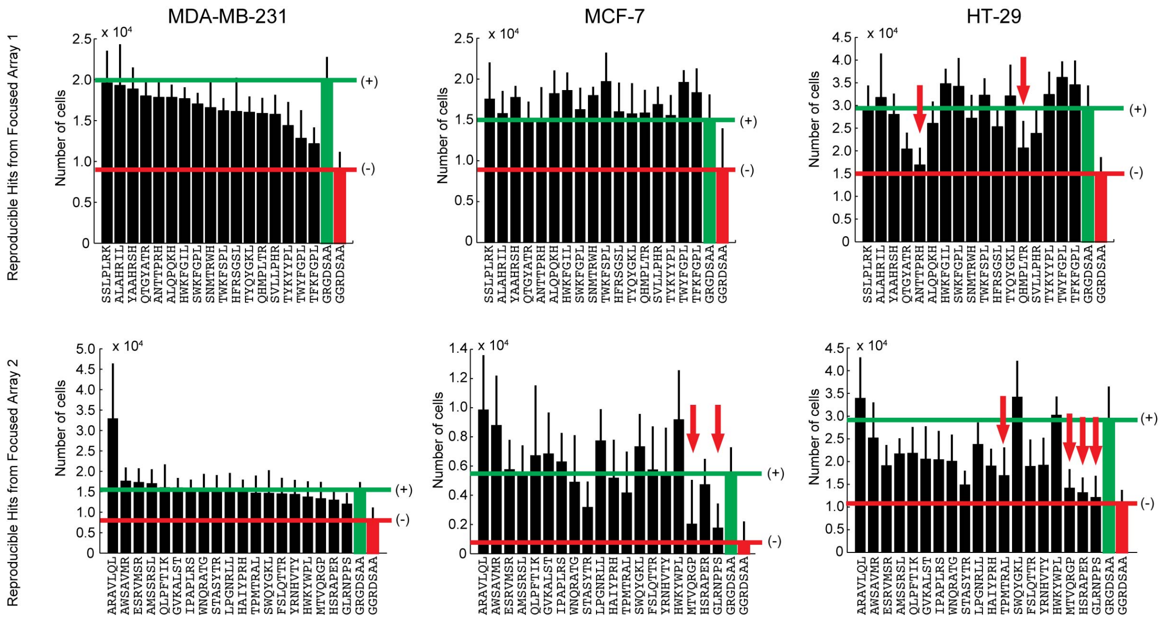


**Figure S16.** Comparison between MDA-MB-231, MCF-7, and HT-29 cell binding to a focused set of peptides. Peptides that bind MDA-MB-231 are sorted in identical order in bar charts describing adhesion of MCF-7 and HT-29 cells. The positive (GRGDSAA, highlighted by green lines) and negative (GGRDSAA, highlighted by red lines) controls were included as references in each bar graph. Red arrows indicate peptides that do not support adhesion of MCF-7 or HT-29 cells. MCF-7 binds to 34, and HT-29 binds to 29 out of 36 MDA-MB-231-binding peptides.

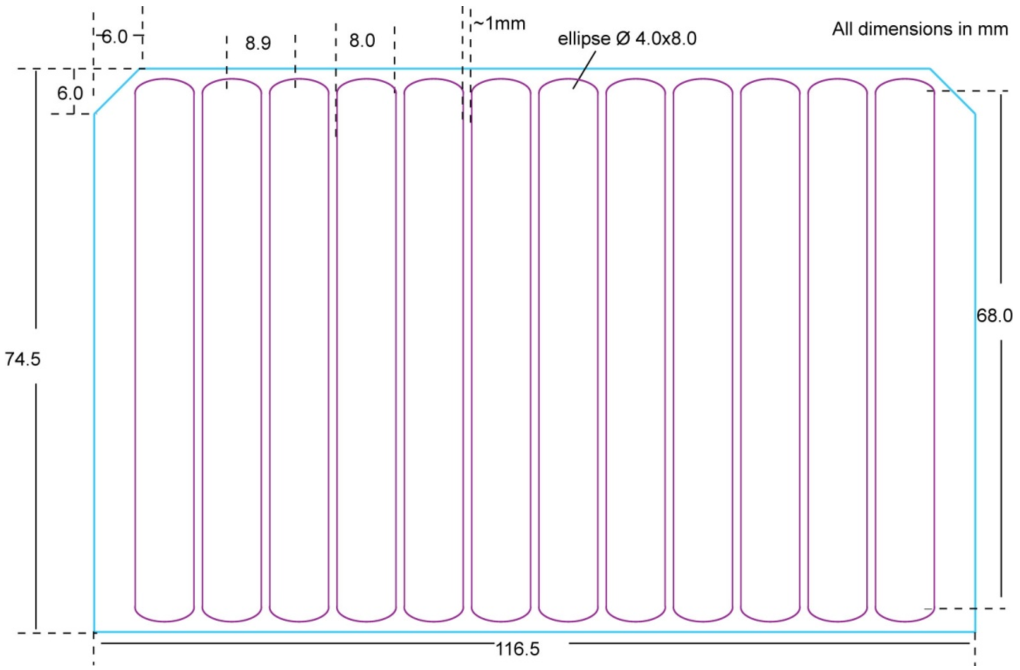


**Figure S17**. Scheme for an aluminium grid insert used to hold peptide paper arrays submerged in a Nunc Omni-Tray for cell adhesion assays.

Description of the files available as part of **Supporting_Files.RAR**

File: "**lot2_Naive_and_Amplified.txt**" includes the copy number of all sequences found in Naïve and amplified Ph.D.-7 libraries for Lots 1 and 2. Snapshot:

```
Nucleotide                 Peptide    L2A1 L2A2 L2A3 L2A4 L2A5 L2N1 L2N2 L2N3 L2N4 L2N5

TGGGGGCGGATTAGTCATGTT      WGRISHV     183  158  221  371  230   34   47   34   16   33
TGGTCGCTGTCGGAGCTTCAT      WSLSELH     220  105  177  278  192   50   57   42   28   43
CTGCTGTATAGGGAGTTTAAT      LLYREFN     142  143  164  303  207   38   66   36   13   60
ACTACGCAGGTGCTGGAGGCT      TTQVLEA     157  123  150  216  151   55   81   50   34   61
GGGTGGGAGACGCGGATGGAG      GWETRME     110  124  112  193  159   20   35   12   11   37
GTTGCGATTCCTACGTCTGTG      VAIPTSV     123  104   94  170  115   31   43   27   11   39
etc
```

Labels:
L2N1, L2N2, L2N3: lot #2 naïve library (replicates 1, 2, 3).
L2A1, L2A2, L2A3: lot #2 library amplified in bulk $10^9$ PFU to $10^{15}$ PFU (rep. 1, 2, 3).

File: "**all_seq_selections.txt**" includes the copy number of all sequences found in all selection using the Ph.D.-7 libraries for Lots 1 and 2. Snapshot:

```
Nucleotide sequence    Peptide  L1B1.1 L1B2.1 L1B3.1 L1B1.2 L1B2.2 L1B3.2 L1B1.3 L1B2.3 L1B3.3 L1E1.1 L1E2.1 L1E3.1 L2B1.1 L2B1.2 L2B1.3 L2E1.1 L2E1.2 L2E1.3

TCGACGGCGTCTTATACTCGT  STASYTR     645   3722   4203      1      4      0    807   3490   1957     17      6      1      0      0      0     38   1695     53
GGGGAGACTCGTGCGCCGCTT  GETRAPL     191   2868  11859    111     36      1      5      0      0     11      6     14      0      0      0     29    478     40
TATGCTGGTCCTTATCAGCAT  YAGPYQH     215   1028   1907    190    141     63      1      2      0      4      2      0      0      0      0      4    437     16
CAGAATACGACTACGGCTCTG  QNTTTAL     482    629    229      0      0      0      0      0      0      3      0      0      0      0      0      7    575     32
```

| label | Library | Condition | Replicate | Round |
|---|---|---|---|---|
| L1B1.1 | PhD7-Lot1 | BA | Replicate 1 | Round 1 |
| L1B2.1 | PhD7-Lot1 | BA | Replicate 1 | Round 2 |
| L1B3.1 | PhD7-Lot1 | BA | Replicate 1 | Round 3 |
| L1B1.2 | PhD7-Lot1 | BA | Replicate 2 | Round 1 |
| L1B2.2 | PhD7-Lot1 | BA | Replicate 2 | Round 2 |
| L1B3.2 | PhD7-Lot1 | BA | Replicate 2 | Round 3 |
| L1B1.3 | PhD7-Lot1 | BA | Replicate 3 | Round 1 |
| L1B2.3 | PhD7-Lot1 | BA | Replicate 3 | Round 2 |
| L1B3.3 | PhD7-Lot1 | BA | Replicate 3 | Round 3 |
| L1E1.1 | PhD7-Lot1 | EmA | Replicate 1 | Round 1 |
| L1E2.1 | PhD7-Lot1 | EmA | Replicate 1 | Round 2 |
| L1E3.1 | PhD7-Lot1 | EmA | Replicate 1 | Round 3 |
| L2B1.1 | PhD7-Lot2 | BA | Replicate 1 | Round 1 |
| L2B1.2 | PhD7-Lot2 | BA | Replicate 2 | Round 1 |
| L2B1.3 | PhD7-Lot2 | BA | Replicate 3 | Round 1 |
| L2E1.1 | PhD7-Lot2 | EmA | Replicate 1 | Round 1 |
| L2E1.2 | PhD7-Lot2 | EmA | Replicate 2 | Round 1 |
| L2E1.3 | PhD7-Lot2 | EmA | Replicate 3 | Round 1 |

File: “**Summary_Lot1.txt**” contains the copy number of all sequences found in the Naïve Ph.D.-7 library for Lot 1 and identification of parasite sequences. This file was obtained from our previous publication.[1] Snapshot:

```
GGGAAGCCTATGCCTCCGATG     GKPMPPM     5548     1
CAGCCTTGGCCGACGAGTATT     QPWPTSI     4478     1
TGGCCTACGCCGCCTTATGCG     WPTPPYA     4044     0
CATGCTCTGGGTCCGTCTTCG     HALGPSS     3066     1
GAGCCGCTGCAGCTGAAGATG     EPLQLKM     2869     1
Etc..
```

Third column contains copy number of sequence. Fourth column indicates whether sequence is a parasite (1) or not (0).

File: “**Summary_Lot2.txt**” contains the copy number of all sequences found in the Naïve Ph.D.-7 library for Lot 2 and identification of parasite sequences. This file was generated by MatLab script **Make_FigureS2b.m** using **lot2_Naive_and_Amplified.txt** as input. Snapshot:

```
ACTACGCAGGTGCTGGAGGCT     TTQVLEA     281    1
TGGTCGCTGTCGGAGCTTCAT     WSLSELH     220    1
CTGCTGTATAGGGAGTTTAAT     LLYREFN     213    1
TGGGGGCGGATTAGTCATGTT     WGRISHV     164    1
GTTGCGATTCCTACGTCTGTG     VAIPTSV     151    1
Etc..
```

Third column contains copy number of sequence. Fourth column indicates whether sequence is a parasite (1) or not (0).

File: “**Parasites_Lot2.txt**” contains the sequences of all parasites found in the Ph.D.-7 library for Lot 2. This file was generated by MatLab script **Make_FigureS2b.m** using **lot2_Naive_and_Amplified.txt** as input. Snapshot:

```
TGGGGGCGGATTAGTCATGTT     WGRISHV     1
TGGTCGCTGTCGGAGCTTCAT     WSLSELH     1
CTGCTGTATAGGGAGTTTAAT     LLYREFN     1
ACTACGCAGGTGCTGGAGGCT     TTQVLEA     1
GGGTGGGAGACGCGGATGGAG     GWETRME     1
GTTGCGATTCCTACGTCTGTG     VAIPTSV     1
ATGATGATTAGTGCGACTGAG     MMISATE     1
Etc..
```

Script: “**Make_FigureS2b.m**”. Running this file will produce Supplementary Figure 2b (volcano plot) and files “**Parasites_Lot2.txt**”, which contains all parasite sequences in the Ph.D.-7 Lot 2 library, and “**Summary_Lot2.txt**”, which contains the copy number of all sequences found in the Naïve Ph.D.-7 library for Lot 2 and identification of parasite sequences (see above).

Script: “**MakeFigure1e.m**”. Running this file will produce plots for Figure 1e and Supplementary Figure 3.

File: “**MakeFigure1f.m**”. Running this file will provide values for the P-, V-, and I-populations from selections using the Ph.D.-7 Lot 1 library. These values were used to make Figure 1f and g. This Script requires “**all_seq_selections.txt**” and “**Summary_Lot1.txt**” as inputs.

File: “**MakeFigure1h.m**”. Running this file will provide values for the P-, V-, and I-populations from selections using the Ph.D.-7 Lot 2 library. These values were used to make **Figure 1h**. This Script requires “**all_seq_selections.txt**” and “**Summary_Lot2.txt**” as inputs.

File: “**readMulticolumn.m**”. This file is a function used to open and read text files.

**References**:


[1] W. L. Matochko, S. C. Li, S. K. Y. Tang, R. Derda, *Nucleic Acids Res.* **2014**, *42*, 1784-1798.

[2] R. Derda, S. K. Y. Tang, G. M. Whitesides, *Angew. Chem. Int. Ed.* **2010**, *49*, 5301-5304.

[3] F. Deiss, W. L. Matochko, N. Govindasamy, E. Y. Lin, R. Derda, *Angew. Chem. Int. Ed.* **2014**, *53*, 6374-6377.
