## Supporting Data (Sequencing, Scripts) for "Reproducible Discovery of Cell-Binding Peptides “Lost” in Bulk Amplification via Emulsion Amplification in Phage Display Panning": Make_FigureS2b.m

```
clear all; close all

Dir='';
File='lot2_Naive_and_Amplified.txt';

Naive = 6:10;
Amplified = 1:5;

skip=2;

[Nuc, AA, Fr0] = readMulticolumn(Dir, File, [Amplified Naive], skip);

FrR=Fr0;
IX=find(Fr0==0);
FrR(IX)=0.3;

% normalize the files by total frequency.
for i=1:size(Fr0,2)
    Fr(:,i)  = 1000*Fr0(:,i)  / (sum(Fr0(:,i)));
    FrR(:,i) = 1000*FrR(:,i) / (sum(Fr(:,i)));
end

Fr1=Fr(:,1:5);
Fr2=Fr(:,6:10);

FrR1=FrR(:,1:5);
FrR2=FrR(:,6:10);

%%

ratio = log2 (mean(FrR1,2)./mean(FrR2,2));
[~,confi] = ttest2(Fr1',Fr2',0.05,'right','unequal');
IX = find(confi < 0.05);


confi(confi<0.0001)=0.0001;
confiL=-log10(confi);

plot(ratio,confiL','.k'); hold on;

line([-1 8],[-log10(0.05) -log10(0.05)]);


plot(ratio(IX),confiL(IX),'.r');
disp(['Found ' num2str(numel(IX)) '  parasites']);

%%
disp('Saving the parasite file');
savename = 'Parasites_Lot2.txt';

IX2 = transpose(IX);

Nucs = Nuc(IX2,:);
AAs = AA(IX2,:);

SP = char(32*ones(size(IX2),5));
NUM = char(49*ones(size(IX2),1));
RET = char(10*ones(size(IX2),1));

toSave = [char(Nucs) SP char(AAs) SP NUM RET];
fprintf( fopen(savename, 'wt'), '%s\r\n', toSave');

fclose all;

%% 
disp('Saving the summary file');
savename = 'Summary_Lot2.txt';

Fre1 = sum(Fr0(:,6:10),2);
IX   = (confi<0.05)';
NUM  = num2str([Fre1 IX]);

SP =  char(32*ones(size(Fre1,1),5));
RET = char(10*ones(size(Fre1,1),1));

toSave = [char(Nuc) SP char(AA) SP NUM RET];
%%
[~,IX3] = sort(Fre1,'descend');

cutoff = min(find(Fre1(IX3)==0));

toSave = toSave(IX3(1:cutoff-1),:);

fprintf( fopen(savename, 'wt'), '%s\r\n', toSave');

fclose all;
```
