## Supporting Data (Sequencing, Scripts) for "Reproducible Discovery of Cell-Binding Peptides “Lost” in Bulk Amplification via Emulsion Amplification in Phage Display Panning": MakeFigure1e.m

```
clear all; close all


Dir = '';
input1='all_seq_selections.txt';
input2='Summary_Lot1.txt';

disp('reading naive sequences and parasites');
[Naive, ~, Fr000] = readMulticolumn(Dir, input2, [1 2], 0);

range= {[1 2 3]; [4 5 6]; [7 8 9]; [10 11 12]};
    
    
for i=1:4
    
    disp('reading the selected sequences')
    [Nuc, AA, Fr] = readMulticolumn(Dir, input1, range{i}, 2);


    % normalize the files by total frequency; those that are 0, make 0.3
    for ii=1:size(Fr,2)
        IX=find(Fr(:,ii)==0);
        M(ii) = (sum(Fr(:,ii)));
        Fr2(:,ii)  = 1*Fr(:,ii)  / M(ii);
        Fr2(IX,ii) = 0.3/M(ii);
    end
    
    figure(i)

    [~,IX]=sort(Fr(:,end),'descend');

    N=1:20;

    isparasite = zeros(size(N));
    isvisible  = zeros(size(N));

    Fr3=[ zeros(numel(N),1) Fr2(IX(N),:) ];
    SUM = sum(Fr000(:,1));

    for j=1:numel(N)
        disp('looking for selected sequences in the naive population')
        where = find(strcmp(Nuc(IX(j)), Naive));

        if where
            Fr3(j,1)      = Fr000(where,1) / SUM;
            isparasite(j) = Fr000(where,2);
            isvisible(j)  = 1;
        else
            isvisible(j)  = 0;
            Fr3(j,1)      = 0.3 / SUM;
        end
    end


    NN = numel(range{i})+1;

    plot(1:NN, Fr3','-k', 'LineWidth', 2); hold on
    
    if sum(find(isvisible))
        plot(1:NN, Fr3(find(isvisible),:)','-b', 'LineWidth', 2); hold on
    end
    
    if sum(find(isparasite))
        plot(1:NN, Fr3(find(isparasite),:)','-r', 'LineWidth', 2); hold on
    end

    set(gca,'YScale','log', 'YLim',[0.00000001 1], 'Box','Off');

    % %%%%%%%%%%%%%%%% USE THIS TO PLACE LIMIT OF SEQUENCING %%%%%%%%%%%
    XXXX = 2 * [0.3/SUM, min(Fr2(:,1)), min(Fr2(:,2)), min(Fr2(:,3))];
    plot(1:NN,  XXXX,'-y', 'LineWidth', 3);
    set(gca, 'YScale','log', 'YLim',[0.00000001 1], 'Box','Off');

end  
```
