## Supporting Data (Sequencing, Scripts) for "Reproducible Discovery of Cell-Binding Peptides “Lost” in Bulk Amplification via Emulsion Amplification in Phage Display Panning": MakeFigure1f.m

```
clear all; close all


Dir = '';
input1='all_seq_selections.txt';
input2='Summary_Lot1.txt';

disp('reading naive sequences and parasites');
[Naive, ~, Fr000] = readMulticolumn(Dir, input2, [1 2], 0);

disp('reading the selected sequences')
    [Nuc, AA, Fr] = readMulticolumn(Dir, input1, 1:12, 2);
    
    
for i=1:12

    [~,IX]=sort(Fr(:,i),'descend');

    N20=1:20;
    N50=1:50;
    
    top20 = Nuc(IX(N20));
    top50 = Nuc(IX(N50));

    isparasite = zeros(size(N20));
    isvisible  = zeros(size(N20));
    disp('looking for selected sequences in the naive population')

    for j=1:numel(N50)
        
        where = find(strcmp(top50(j), Naive));

        if where
            isparasite(j) = Fr000(where,2);
            isvisible(j)  = 1;
        else
            isvisible(j)  = 0;
            isparasite(j) = 0;
        end
    end
    
    disp(['parasite in top20/50  ' num2str(sum(isparasite(1:20))) ...
           ' / ' num2str(sum(isparasite))]);
    disp(['visible in top20/50   '  num2str(sum(isvisible(1:20))) ...
           ' / ' num2str(sum(isvisible))  ]);
    disp(['invisible in top20/50 ' num2str(sum(~isvisible(1:20))) ...
           ' / ' num2str(sum(~isvisible))]);
       
   disp(' ****************   make a bar: ****************************');
   disp(['Large dots red ' num2str(sum(isparasite(1:20))) ...
        ' blue ' num2str(sum(isvisible(1:20)) - sum(isparasite(1:20))) ...
        ' white ' num2str(sum(~isvisible(1:20))) ...
        ' Small dots red ' num2str(sum(isparasite(21:50))) ...
        ' blue ' num2str(sum(isvisible(21:50)) - sum(isparasite(21:50))) ...
        ' white ' num2str(sum(~isvisible(21:50))) ]);
   disp(' *********************************************************'); 
        

end  
```
