## Supporting Data (Sequencing, Scripts) for "Reproducible Discovery of Cell-Binding Peptides “Lost” in Bulk Amplification via Emulsion Amplification in Phage Display Panning": readMulticolumn.m

```
function [Nuc, AA, Fr] = readMulticolumn(Dir, File, column, skip)

FORM = '%s %s';

for i=1:max(column)
    if sum(i==column)
        FORM = [FORM ' %f'];
    else
        FORM = [FORM ' %*f'];
    end
end

FORM = [FORM ' %*[^\n]'];

fh = fopen(fullfile(Dir, File),'r');
for i=1:skip
    temp=fgetl(fh);
end
 
AllVar = textscan(fh,FORM);
Nuc = AllVar{1};
AA  = AllVar{2};

Fr=[];
for i=1:numel(column)    
    Fr = [Fr AllVar{2+i}];
end

% %remove the NaNs
% 
% [r,~] = find(~isnan(Fr));
% 
% r=unique(r);
% 
% Fr=Fr(r,:);
% Nuc=Nuc(r);
% AA=AA(r);

clear AllVar;
fclose all;
end
```
